# Temporal filters in response to presynaptic spike trains: Interplay of cellular, synaptic and short-term plasticity time scales

**DOI:** 10.1101/2021.09.16.460719

**Authors:** Yugarshi Mondal, Rodrigo F. O. Pena, Horacio G. Rotstein

## Abstract

Temporal filters, the ability of postsynaptic neurons to preferentially select certain presynaptic input patterns over others, have been shown to be associated with the notion of information filtering and coding of sensory inputs. Short-term plasticity (depression and facilitation; STP) has been proposed to be an important player in the generation of temporal filters. We carry out a systematic modeling, analysis and computational study to understand how characteristic postsynaptic (low-, high- and band-pass) temporal filters are generated in response to periodic presynaptic spike trains in the presence STP. We investigate how the dynamic properties of these filters depend on the interplay of a hierarchy of processes, including the arrival of the presynaptic spikes, the activation of STP, its effect on the excitatory synaptic connection efficacy, and the response of the postsynaptic cell. These mechanisms involve the interplay of a collection of time scales that operate at the single-event level (roughly, during each presynaptic interspike-interval) and control the long-term development of the temporal filters over multiple presynaptic events. These time scales are generated at the levels of the presynaptic cell (captured by the presynaptic interspike-intervals), short-term depression and facilitation, synaptic dynamics and the post-synaptic cellular currents. We develop mathematical tools to link the single-event time scales with the time scales governing the long-term dynamics of the resulting temporal filters for a relatively simple model where depression and facilitation interact at the level of the synaptic efficacy change. We extend our results and tools to account for more complex models. These include multiple STP time scales and non-periodic presynaptic inputs. The results and ideas we develop have implications for the understanding of the generation of temporal filters in complex networks for which the simple feedforward network we investigate here is a building block.

## 1 Introduction

The synaptic communication between neurons involves a multiplicity of interacting time scales and is affected by a number of factors including short-term plasticity [1–3], primarily involved in information filtering, long-term plasticity [4,5], involved in learning and memory [6], homeostatic plasticity [7], involved in the maintenance of function in the presence of changing environments, neuromodulation [8, 9], and astrocyte regulation [10, 11], in addition to the temporal properties of the presynaptic spikes, the intrinsic currents of the postsynaptic neurons and background noise activity.

Short-term plasticity (STP) refers to the increase (synaptic facilitation) or decrease (synaptic depression) of the efficacy of synaptic transmission (strength of the synaptic conductance) in response to repeated presynaptic spikes with a time scale in the range of hundreds of milliseconds to seconds [1–3, 12, 13]. STP is ubiquitous both in invertebrate and vertebrate synapses, and has been shown to be important for neuronal computation [14–18] and information filtering (temporal and frequency-dependent) [2, 12, 19–41], and related phenomena such as burst detection [27, 38], temporal coding and information processing [27, 28, 42–45], gain control [15, 46, 47], information flow [16, 36, 48] given the presynaptic history-dependent nature of STP, the prolongation of neural responses to transient inputs [49–51], the modulation of network responses to external inputs [52, 53], hearing and sound localization [54, 55], direction selectivity [56], attractor dynamics [57] (see also [47]), the generation of cortical up and down states [58], navigation (e.g., place field sensing) [30, 33], vision (e.g., microsacades) [59], working memory [51, 60] and decision making [61].

The notion of information filtering as the result of STP is associated with the concept of temporal filters [12, 22–24] at the synaptic and postsynaptic levels, which are better understood in response to periodic presynaptic inputs [38, 62–64] for a wide enough range of input frequencies. (See the schematic diagrams in Figs. 1-A and 2-A where *x* and *z* describe the evolution over time of synaptic depression and facilitation, respectively, and their product describes their combined activity.) In spite of the ubiquitousness of STP and the consequences for information filtering [12, 22, 65], the mechanisms of generation of postsynaptic temporal filters in response to presynaptic input spikes are not well understood.

**Figure 1:**
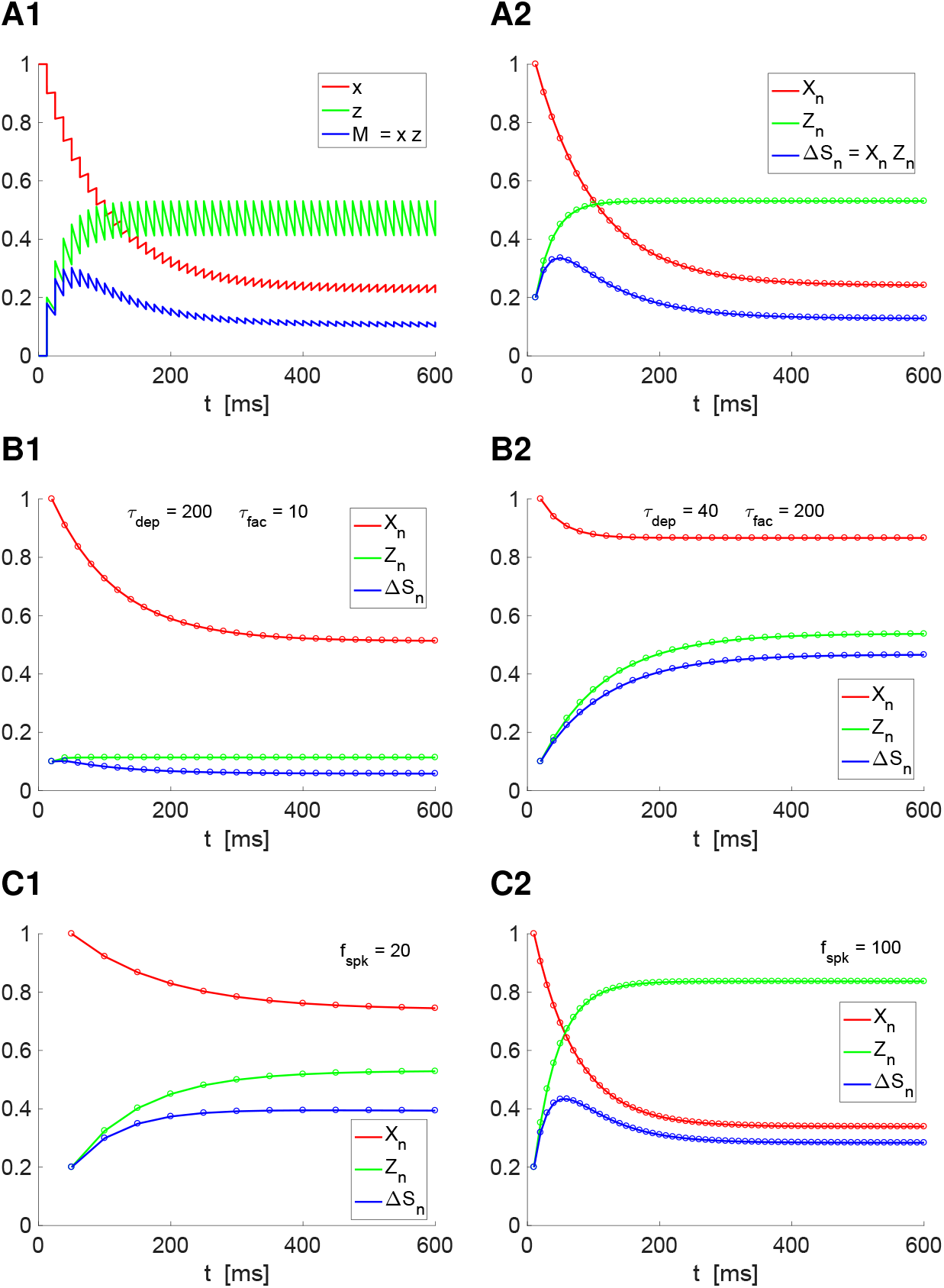
Short-term depression and facilitation and the generation of temporal filters in response to periodic presynaptic inputs. **A1.***x*-, *z*- and *M*-traces (curves of *x*, *z* and *M* = *xz* as a function of *t*). **A2.** Circles: *X_n_*-, *Z_n_*- and Δ*S_n_* = *X_n_Z_n_*-peak sequence computed using (7)-(8). Solid curves: join the *X_n_*-, *Z_n_*- and Δ*S_n_* = *X_n_Z_n_*-envelope peak sequences computed using the caricature (descriptive) model (31)-(34). The values of the envelope peaks decay constants are *σ_d_* ~ 91.5 and *σ_f_* ~ 26.4. We used the simplified model (4)-(6) and the following parameter values: *τ_dep_* = 400, *τ_fac_* = 50, *a_d_* = 0.1, *a_f_* = 0.2, *x*_∞_ = 1, *z*_∞_ = 0, *f_spk_* = 80 Hz (presynaptic input frequency). **B.** Depression- and facilitation-dominated peak sequences. **B1.** Depression-dominated temporal filter regime. **B2.** Facilitation-dominated temporal filter regime. We used the simplified model (4)-(6) and the following parameter values: *τ_dep_* = 200 (B1), *τ_dep_* = 40 (B2), *τ_fac_* = 10 (B1), *τ_fac_* = 200 (B2), *a_d_* = 0.1, *a_f_* = 0.1, *x*_∞_ = 1, *z*_∞_ = 0, *f_spk_* = 50 Hz. **C.** Input frequency-dependent temporal filters. **C1.** High-pass temporal filter for low spiking input frequencies (*f_spk_* = 20). **C2.** Band-pass temporal filter for higher spiking input frequencies (*f_spk_* = 100). We used the simplified model (4)-(6) and the following parameter values: *τ_dep_* = 200, *τ_fac_* = 200, *a_d_* = 0.1, *a_f_* = 0.2, *x*_∞_ = 1, *z*_∞_ = 0.

**Figure 2:**
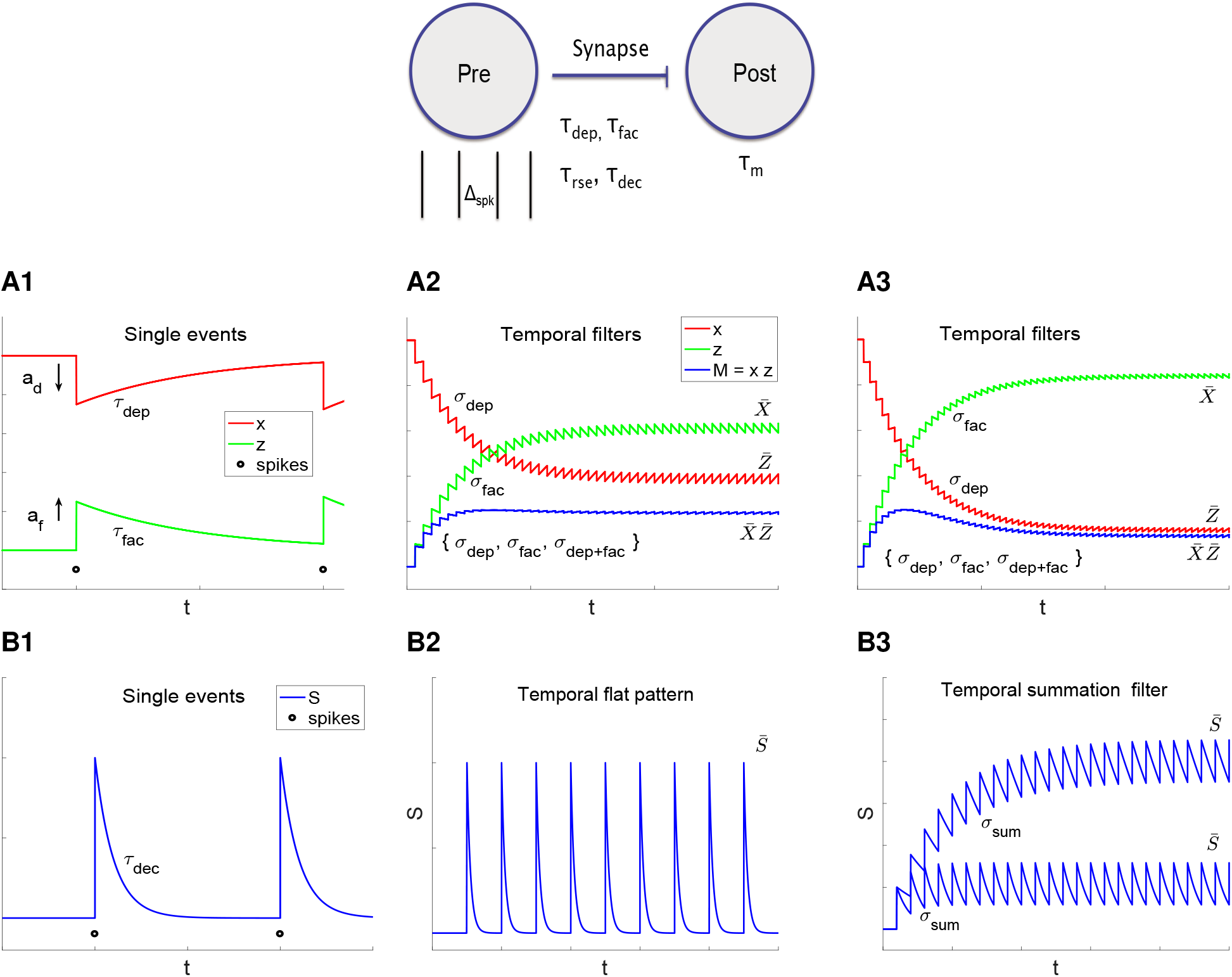
Single event and temporal filters’ time scales and other attributes in response to presynaptic spike trains in the presence of synaptic depression (*x*) and facilitation (*z*). The presynaptic cell is modeled as a periodic spike train with period Δ_*spk*_. The postsynaptic cell is modeled as a passive cell (capacitive and leak currents) with a membrane time constant *τ_m_*. The excitatory synaptic function *S* raises and decays with time constants *τ_rse_* and *τ_dec_*, respectively. The synaptic depression and facilitations are *τ_dep_* and *τ_fac_*, respectively. **A.** Depression and facilitation. **A1.** Single events. At the arrival of each presynaptic spike (black dots), the depression (*x*) and facilitation (*z*) variables decrease and increase, respectively. In the models we use in this paper, they are discretely updated. They decay towards their (single event) steady-states (*x*_∞_ = 1 and *z*_∞_ = 0) with the (single event) time scales *τ_dep_* and *τ_fac_*, respectively. **A2, A3.** Temporal patterns (filters) generated by presynaptic spike trains with different ISIs Δ_*spk*_ (or frequencies *f_spk_*) and the dynamics of the single events (controlled by *τ_dep_* and *τ_fac_*). Depression and facilitation always give rise to low- (red) and high- (green) pass filters respectively. Their product can be a depression-dominated (low-pass) filter, facilitation-dominated (high-pass) filter (A2), or a band-pass filter (A3). The (emergent, long term) filter time scales *σ_dep_*, *σ_fac_* and *σ_dep_*_+*fac*_ depend on the interplay of *τ_dep_*, *τ_fac_* and Δ_*spk*_. The temporal filter steady-states are captured by the peak sequence steady-states 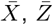 and 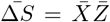. **B.** Synaptic dynamics. **B1.** Single events. At the arrival of each presynaptic spike (black dots) the synaptic variable *S* increases instantaneously (*τ_rse_* = 0) and then decreases with a time constant *τ_dec_*, which defines the decay time scale. In the models we use in this paper, *S* is discretely updated. **B2, B3.** Temporal patterns (filters) generated by presynaptic spike trains with different ISIs Δ_*spk*_ (or frequencies *f_spk_*) and the dynamics of the single events (controlled by *τ_dec_*). For small *τ_dec_* and Δ_*spk*_, the *S* pattern is flat (B2). For larger values of *τ_dec_* and Δ_*spk*_, summation generates a high-pass filter. The emergent time scales depend on the interplay of *τ_dec_* and Δ_*spk*_. The temporal filter steady-states are captured by the peak sequence steady-states 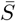.

One difficulty is that the notion of temporal filters has not been precisely defined. Temporal filters have been broadly characterized as biological systems that allow certain information carried out by the presynaptic spike pattern to pass to the postsynaptic neuron with possibly a modification (attenuation or amplification) in the firing rate, while other information is rejected [22,66]. A systematic mechanistic study requires a more precise characterization that takes into account the underlying complexity. First, postsynaptic temporal filters result from the concatenation of various processes: the structure of the presynaptic spike patterns, STP, synaptic dynamics and the intrinsic dynamics of the postsynaptic cell resulting from the intrinsic currents (diagram in Fig. 2). It is not well understood how the time scales associated with the dynamics of synaptic depression and facilitation interact with the presynaptic spike train time scales (interspike intervals, ISIs) and the membrane potential time scales to generate the resulting temporal filters. Second, temporal filters are a transient phenomenon in the time domain, in addition to being frequency-dependent [12]. Therefore, the steady-state postsynaptic membrane potential profiles (curves of the postsynaptic membrane potential amplitudes or peaks as a function of the presynaptic input frequency) [38, 67, 68] does not necessarily capture the system’s filtering properties. (These steady-state profiles are the natural extensions of the impedance profiles for subthreshold resonance in neurons.) Third, the STP’s history-dependent properties generate a significant amount of variability in the STP-mediated temporal patterns due to the multiple possible arrangements of ISIs’ durations in non-periodic presynaptic input patterns.

In this paper, we adopt the use of periodic presynaptic spike patterns as the reference presynaptic spike trains to define and characterize the various types of temporal filters that emerge and investigate the mechanisms by which they are generated. This can serve as the reference point for the investigation of the filtering properties of temporal patterns in response to more complex presynaptic patterns (e.g., bursting, Poisson distributed). Periodic presynaptic spike trains have been used by other authors [38, 62–64] to illustrate the emergence of temporal patterns in the presence of STP.

We focus on the feedforward network described in the diagram in Fig. 2, which is the minimal model that can show postsynaptic membrane potential temporal filters in response to presynaptic spike trains in the presence of STP. We leave out the postsynaptic firing rate responses. In some cases, they can be directly derived from the membrane potential responses.

Phenomenological models of synaptic depression and facilitation [21, 43, 62, 63, 67–73] describe the evolution of two variables that abruptly decrease and increase, respectively, by a certain amount in response to each presynaptic spike and relax towards their steady-state values during the presynaptic ISIs (see Fig. 2-A1 for the depression and facilitation variables *x* and *z*, respectively). At the arrival of each presynaptic spike, the synaptic function (*S*) is updated by an amount Δ*S* equal to the appropriate product of *x* and *z* at the arrival time. The cumulative effect of these single-spike events along the sequence of presynaptic spikes generates temporal patterns in the variables *x*, *z* and *S* (Figs. 1 and 2), which are transmitted to the postsynaptic cell (diagram in Fig. 2) to produce postsynaptic temporal filters.

The temporal filters for the variables *x* and *z* are better captured by the sequences of peak values *X_n_* and *Z_n_* (for the spike index *n*) (Figs. 1) whose evolution is characterized by the (long-term) time scales (*σ_dep_* and *σ_fac_*) and the steady state values (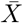 and 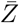) Fig. (2). Because of their monotonic decreasing (*X_n_*) and increasing (*Z_n_*) properties, we refer to them as temporal low-pass (*X_n_*) and high-pass (*Z_n_*) filters, respectively. The synaptic update is the product Δ*S_n_* = *X_n_Z_n_* and the corresponding filter can have a transient peak, which we refer to as a temporal band-pass filter and, as we show, it involves an additional (long-term) time scale (*σ_dep_*_+*fac*_). These time scales depend on the single-event time scales (*τ_dep_* and *τ_fac_*) and the presynaptic ISI (Δ_*spk*_) in complex ways. In addition, the phenomenon of summation in a postsynaptic cell in response to presynaptic inputs may develop an additional (postsynaptic) high-pass temporal filter (Fig. 2-B), which is independent of the ones described above, and is characterized by the (long-term) time scale (*σ_sum_*) and the steady state value 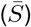. They depend on the membrane time constant, the synaptic decay time *τ_dec_* and the presynaptic Δ_*spk*_. For relatively fast synapses (e.g., AMPA), summation is not observed at the synaptic level, but at the postsynaptic level, and depends on the time scale of the postsynaptic cell (*τ_m_*) and the presynaptic Δ_*spk*_.

A key idea we develop in this paper is that of the communication of time scales (i) across levels of organization (presynaptic, STP, synaptic, postsynaptic; Fig. 2) and (ii) from these operating at the single event level (e.g., *τ_dep_*, *τ_fac_*, *τ_dec_*, *τ_m_*, Δ_*spk*_) to the (long-term) ones operating at the filter level (e.g., *σ_dep_*, *σ_fac_*, *σ_sum_*). This notion of communication involves the complex interaction of time scales and generation of new time scales. We use this framework to organize our mechanistic understanding of the temporal filtering phenomena. However, we note that while in some cases the time scales can be easily represented by time constants, in other cases they are more difficult to be precisely characterized.

More specifically, we use biophysically plausible (conductance-based) mathematical modeling and dynamical systems tools to systematically understand how the postsynaptic low-, high- and band-pass temporal filters are generated in response to presynaptic spike trains in the presence of STP. Using a combination of analytical and computational tools, we describe the dependence of the dynamic properties of these filters, captured by the long-term time scales, on the interplay of the hierarchy of processes, ranging from the arrival of the presynaptic spike trains, to the activation of STP to the activation of the synaptic function to the response of the postsynaptic cell (Fig. 2, diagram). In particular we describe how all this depends on the time scales of the building blocks (*τ_dep_*, *τ_fac_*, *τ_dec_*, *τ_m_* and the presynaptic Δ_*spk*_). We then extend our results and tools to account for more complex models. These include synaptic depression and facilitation processes with multiple time scales and non-periodic presynaptic synaptic inputs.

The conceptual and mathematical framework we introduce to develop these ideas and identify the contribution of each of the network components to the generation of temporal filters can be extended to understand the filtering and coding properties of more complex scenarios. These involve more realistic description of the participating processes at the various levels of organization and the presynaptic input spike trains (e.g., bursting patterns). Finally, the results and ideas we develop have implications for the understanding of the generation of temporal filters in complex networks for which the simple feedforward network we investigate here is a building block.

## 2 Methods

### 2.1 Models

#### 2.1.1 Postsynaptic cell: leaky integrate-and-fire model

The current-balance equation for the post-synaptic cell is given by

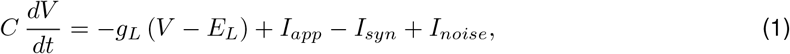

where *t* is time (ms), *V* represents the voltage (mV), *C* is the specific capacitance (*μ*F/cm^2^), *g_L_* is the leak conductance (mS/cm^2^), *I_app_* is the tonic (DC) current (*μ*A/cm^2^)), 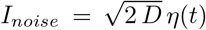 represents white noise (delta correlated with zero mean), and *I_syn_* is an excitatory synaptic current of the form

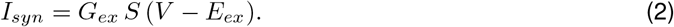

Here *G_ex_* is the maximal synaptic conductance (mS/cm^2^), *E_ex_* = 0 is the reversal potential for AMPA excitation, and the synaptic variable *S* obeys a kinetic equation of the form

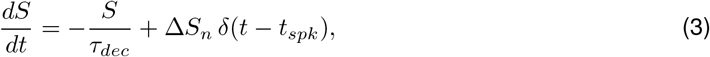

where *τ_dec_* (ms) is the decay time of excitation. Each presynaptic spike instantaneously raises *S* to some value Δ*S_n_* which varies depending on the properties of the short-term dynamics (depression and/or facilitation) and defined below. We refer the reader to [69, 74] for additional details on biophysical (conductance-based) models.

#### 2.1.2 Presynaptic spike-trains

We model the spiking activity of the presynaptic cell as a spike train with presynaptic spike times *t*_1_, *t*_2_*,…, t_N_*. We consider two types of input spike-trains: uniformly and Poisson distributed. The former is characterized by the interspike interval (ISI) of length Δ_*spk*_ (or its reciprocal, the spiking frequency *f_spk_*) and the latter are characterized by the mean spiking rate (or the associated exponential distribution of ISIs).

#### 2.1.3 The DA (Dayan-Abbott) phenomenological model for short-term dynamics: synaptic depression and facilitation

This simplified phenomenological model has been introduced in [69] by Dayan and Abbott. It is relatively simpler than the well-known phenomenological MT (Markram-Tsodkys) model described below [63]. In particular, the depression and facilitation processes are independent (the updates upon arrival of each presynaptic spike are uncoupled). We use it for its tractability and to introduce some conceptual ideas.

The magnitude Δ*S* of the synaptic release per presynaptic spike is assumed to be the product of the depressing and facilitating variables

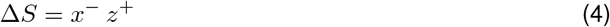

where

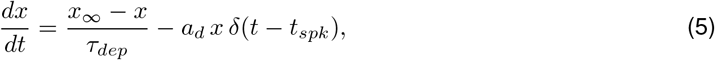

and

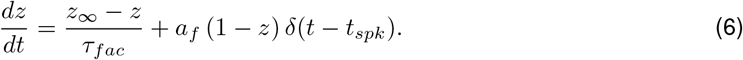

Each time a presynaptic spike arrives (*t* = *t_spk_*), the depressing variable *x* is decreased by an amount *a_d_ x* (the release probability is reduced) and the facilitating variable *z* is increased by an amount *a_f_* (1 − *z*) (the release probability is augmented). During the presynaptic interspike intervals (ISIs) both *x* and *z* decay exponentially to their saturation values *x*_∞_ and *z*_∞_ respectively. The rate at which this occurs is controlled by the parameters *τ_dep_* and *τ_fac_*. Following others we use *x*_∞_ = 1 and *z*_∞_ = 0. The superscripts “ ± ” in the variables *x* and *z* indicate that the update is carried out by taking the values of these variables prior (^−^) or after (^+^) the arrival of the presynaptic spike.

Fig. 1-A1 illustrates the *x*-, *z*- and *M* = *xz*-traces (curves of *x*, *z* and *M* as a function of time) in response to a periodic presynaptic input train for representative parameter values. (Note that *M* = *x z* is defined for all values of *t*, while Δ*S*= *x*^−^*z*^+^ is used for the update of *S* after the arrival of spikes and Δ*S_n_* = *X_n_Z_n_* is the sequence of peaks.)

#### 2.1.4 DA model in response to presynaptic inputs

##### Peak dynamics and temporal filters

By solving the differential equations (5)–(6) during the presynaptic ISIs and appropriately updating the solutions at *t* = *t_n_* (occurrence of each presynaptic spike), one arrives at the following recurrent formula for the peak sequences in terms of the model parameters

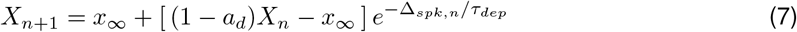

and

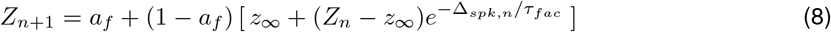

where 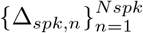 represents the lengths of the presynaptic ISIs.

Fig. 1-A2 illustrates the peak envelopes (curves joining the peak sequences for *X_n_*, *Z_n_* and Δ*S_n_* = *X_n_Z_n_*, circles) for the parameter values in Fig. 1-A1. These are sequences indexed by the input spike number, which we calculate analytically below. As expected, *X_n_* is a decreasing sequence (temporal low-pass filter) and *Z_n_* is an increasing sequence (temporal high-pass filter). Their product (computed so that the peak of the product is the product of the peaks) exhibits a transient peak (temporal band-pass filter).

##### Steady-state frequency-dependent filters

For periodic inputs, Δ_*spk,n*_ is independent of *n* (Δ_*spk*_) and eqs. (7)–(8) are linear 1D difference equations. Therefore both the sequences *X* and *Z* obey linear discrete dynamics (e.g., see [75]), decaying to their steady state values

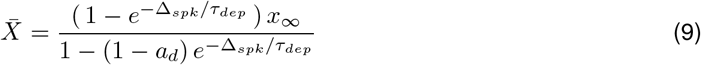

and

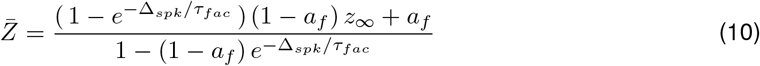

as shown in Figs. 1-B and -C (red and green).

#### 2.1.5 The MT (Markram-Tsodkys) phenomenological model for short-term dynamics: synaptic depression and facilitation

This model was introduced in [63] as a simplification of earlier models [1, 43, 76]. It is more complex and more widely used than the DA model described above [38, 77], but still a phenomenological model.

As for the DA model, the magnitude Δ*S* of the synaptic release per presynaptic spike is assumed to be the product of the depressing and facilitating variables:

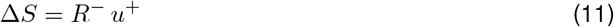

where, in its more general formulation,

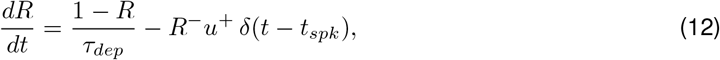

and

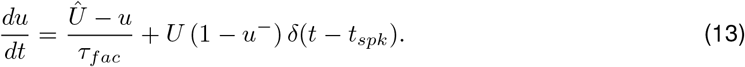

Each time a presynaptic spike arrives (*t* = *t_spk_*), the depressing variable *R* is decreased by *R*^−^*u*^+^ and the facilitating variable *u* is increased by *U* (1 − *u*^−^). As before, the superscripts “ ± ” in the variables *R* and *u* indicate that the update is carried out by taking the values of these variables prior (^−^) or after (^+^) the arrival of the presynaptic spike. In contrast to the DA model, the update of the depression variable *R* is affected by the value of the facilitation variable *u*^+^. Simplified versions of this model include making 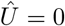 [21, 62, 63, 67, 73, 78] and 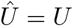 [38].

#### 2.1.6 MT model in response to presynaptic inputs

##### Peak dynamics and temporal filters

By solving the differential equations (12)–(13) during the presynaptic ISIs and appropriately updating the solutions at *t* = *t_n_* (occurrence of each presynaptic spike), one arrives at the following recurrent formula for the peak sequences in terms of the model parameters

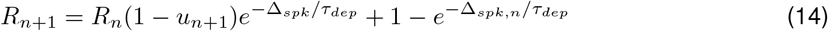

and

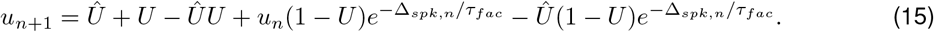

##### Steady-state frequency-dependent filters

As before, for presynaptic inputs Δ_*spk,n*_ is independent of *n* and these equations represent a system two 1D difference equations, which are now nonlinear. The steady-state values are given by

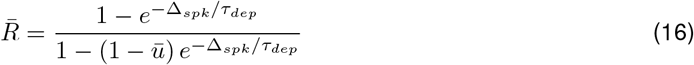

and

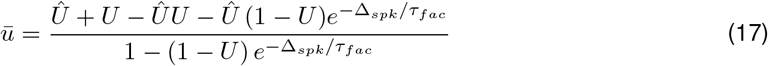

#### 2.1.7 Synaptic dynamics in response to periodic presynaptic inputs and a constant update

##### Peak dynamics

By solving the differential equation (3) for a constant value of Δ*S_n_* = Δ*S* during the presynaptic ISIs and updating the solution at each occurrence of the presynaptic spikes at *t* = *t_n_*, *n* = 1*,…, N_spk_*, one arrives to the following discrete linear differential equation for the peak sequences in terms of the model parameters

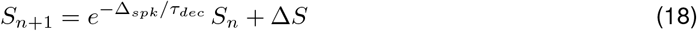

##### Steady-states and frequency filters

The steady state values of (18) are given by

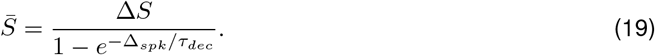

### 2.2 Numerical simulation

The numerical solutions were computed using the modified Euler method (Runge-Kutta, order 2) [79] with a time step Δ*t* = 0.01 ms (or smaller values of Δ*t* when necessary) in MATLAB (The Mathworks, Natick, MA). The code is available at \https://github.com/BioDatanamics-Lab/temporal_filters_p20_01.

## 3 Results

### 3.1 Temporal summation filters for linear synaptic dynamics and constant updates: the single-event and the (long-term) filter time scales coincide

As discussed above, the mechanisms of generation of temporal filters involve the communication of the time scales from the single event (the *τ*’s) to the filter levels (the *σ*’s) (Fig. 2). These two classes of time scales are generally different reflecting the complexity of the process (e.g., the updates of the corresponding variables at the arrival of each presynaptic spike are non-constant, state-dependent). Here we discuss the special case of synaptic summation (linear single event dynamics and constant update) for which both types of time scales coincide. This is relevant both as a reference case and because *S* is a component o the feedforward network we investigate here.

Temporal summation filters (SFs, Fig. 2-B3) refer to the long-term patterns generated in the response of a dynamical system to periodic stimulation with constant amplitude by the accumulation of the responses produced by the single events (cycles). Temporal summation synaptic filters are high-pass filters (HPFs) and naturally develop in the response of linear systems such as eq. (18) with a constant update (independent of the spike index) where the activity *S* decays during the presynaptic ISI and it is updated in an additive manner at the arrival of each presynaptic spike. If the quotient Δ_*spk*_/τ_*dec*_ is finite, *S* will not be able to reach a small enough vicinity of zero before the next presynaptic spike arrives and then the *S*-peak envelope will increase across cycles. While summation does not require the presynaptic inputs to be periodic or the input to have constant amplitude, the notion of summation filter we use here does.

The solution to equation (18) is given by

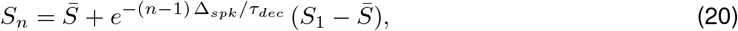

where *S*_1_ = Δ*S* (see Appendix A). This equation describes the temporal filter in response to the presynaptic spike train. For technical purposes, one can extend eq. (20) to include the point (0, 0), obtaining

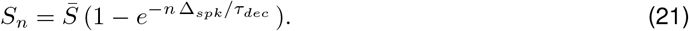

A further extension to the continuous domain yield

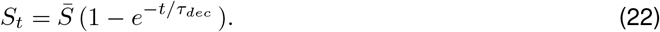

which is the solution to

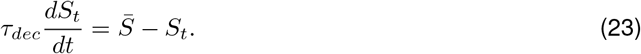

We use the notation *S_t_* instead of *S*(*t*) to emphasize the origin of *S_t_* as the continuous extension of a discrete sequence rather than the evolution of Eq. (3).

Together these results show the temporal SF and the single events are controlled by the same time constant *τ_dec_*. While the time scale is independent of Δ_*spk*_, the steady-state 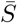 is Δ_*spk*_-dependent.

### 3.2 Dynamics of the depression (*x*) and facilitation (*z*) variables and their interaction: emergence of temporal low-, high-and band-pass filters

#### 3.2.1 From single events (local in time) to temporal patterns and filters (global in time)

The dynamics of single events for the variables *x* (depression) and *z* (facilitation) are governed by eqs. (5)–(6), respectively. After the update upon the arrival of a spike, *x* and *z* decay towards their saturation values *x*_∞_(= 1) and *z*_∞_(= 0), respectively.

The response of *x* and *z* to repetitive input spiking generates patterns for these variables in the temporal domain (e.g., Fig. 1-A1) and for the peak sequences *X_n_* and *Z_n_* in the (discrete) presynaptic spike-time domain (e.g., Fig. 1-A2). The latter consist of the transition from the initial peaks *X*_1_ and *Z*_1_ to 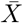 and 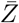, respectively, as *n* → ∞. The properties of these patterns depend not only on the parameters for the single events (*τ_dep/fac_* and *a_d/f_*), but also on the input frerquency *f_spk_* (or the presynaptic ISI Δ_*spk*_) as reflected by eqs. (9)–(10) describing the peak-envelope steady-state values. We note that we use the notation *X_n_* and *Z_n_* for the peak envelope sequences to refer to the sequences 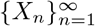 and 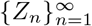.

The peak envelope patterns have emergent, long-term time constants, for which we use the notation *σ_dep_* and *σ_fac_*. As we show in more detail later in the next section, *σ_dep/fac_* depend on *τ_dep/fac_*, Δ_*spk*_ and *a_d/f_* in a relatively complex way. This is in contrast to our discussion in the previous section for the synaptic dynamics where the single event and long-term time scales coincide.

The Δ*S_n_* envelope patterns combine these time scales in ways that involve different levels of complexity. We refer to the Δ*S_n_* patterns that are monotonically decreasing (e.g., Fig. 1-B1) and increasing (e.g., Fig. 1-B2) as temporal low- and high-pass filters (LPFs and HPFs), respectively. We refer to the Δ*S_n_* patterns that exhibit a peak in the temporal domain (e.g., Fig. 1-A2) as temporal band-pass filters (BPFs). This terminology is extended to the peak envelopes *X_n_* (temporal LPFs) and *Z_n_* (temporal HPFs).

#### 3.2.2 Depression- / facilitation-dominated regimes and transitions between them

In the absence of either facilitation or depression, the Δ*S_n_* temporal LPFs and HPFs reflect the presence of depressing or facilitating synapses, respectively. However, Δ*S_n_* temporal LPFs and HPFs need not be generated by pure depression and facilitation but can reflect different balances between these processes where either depression (Fig. 1-B1) or facilitation (Fig. 1-B2) dominates.

It is instructive to look at the limiting cases. A small enough value of *τ_dep/fac_* causes a fast recovery to the saturation value (*x*_∞_ or *z*_∞_) and therefore the corresponding sequence (*X_n_* or *Z_n_*) is almost constant. In contrast, a large enough value of *τ_dep/fac_* causes a slow recovery to the saturation value and therefore the corresponding sequence shows a significant decrease (*X_n_*) or increase (*Z_n_*) as the result of the corresponding underlying variables (*x* and *z*), being almost constant during the ISI. Therefore, when *τ_dep_ ≫ τ_fac_*, depression dominates (Fig. 1-B1) and when *τ_dep_ ≪ τ_fac_*, facilitation dominates (Fig. 1-B2). In both regimes, the exact ranges depend on the input frequency *f_spk_*. An increase in *f_spk_* reduces the ability of *x* and *z* to recover to their saturation values within each presynaptic ISI, and therefore amplifies the depression and facilitation effects over the same time interval and over the same amount of input spikes (compare Figs. 1-C1 and -C2). Therefore, the different balances between *X_n_* and *Z_n_* as *f_spk_* generate different types of Δ*S_n_* patterns and may cause transitions between qualitatively different Δ*S_n_* patterns.

### 3.3 Dependence of the temporal (depression and facilitation) LPFs and HPFs on the single event time scales

#### 3.3.1 Communication of the single event time scales to the (long-term) history-dependent filters

Here we focus on understanding how the (long-term) time scales of the peak envelope sequences *X_n_* and *Z_n_* (*σ_dep_* and *σ_fac_*, respectively) result from the interaction between the time constants for the corresponding single events (*τ_dep_* and *τ_fac_*) and the presynaptic spike input time scales Δ_*spk*_.

Standard methods (see Appendix A with Δ_*spk,n*_ = Δ_*spk*_, independent of *n*) applied to difference eqs. (7)–(8) yield

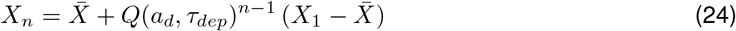

and

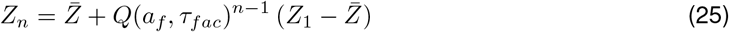

for *n* = 1*,…, N_spk_*, where

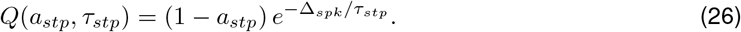

The evolution of the temporal patterns *X_n_* and *Z_n_* are controlled by the behavior of *Q*(*a_d_, τ_dep_*)^*n*−1^ and *Q*(*a_f_, τ_fac_*)^*n*−1^ as *n* → ∞. Because both approach zero as *n* → ∞ (e.g., Figs. 3, gray), the *X_n_* and *Z_n_* patterns decrease and increase monotonically to 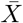 and 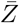, respectively (e.g., Figs. 1 and 3, red and green dots, respectively). The convergence for *a_stp_* < 1 is guaranteed by the fact that *Q*(*a_stp_, τ_stp_*) < 1. Biophysically plausible values of *a_d_* and *a_f_* are well within this range.

**Figure 3:**
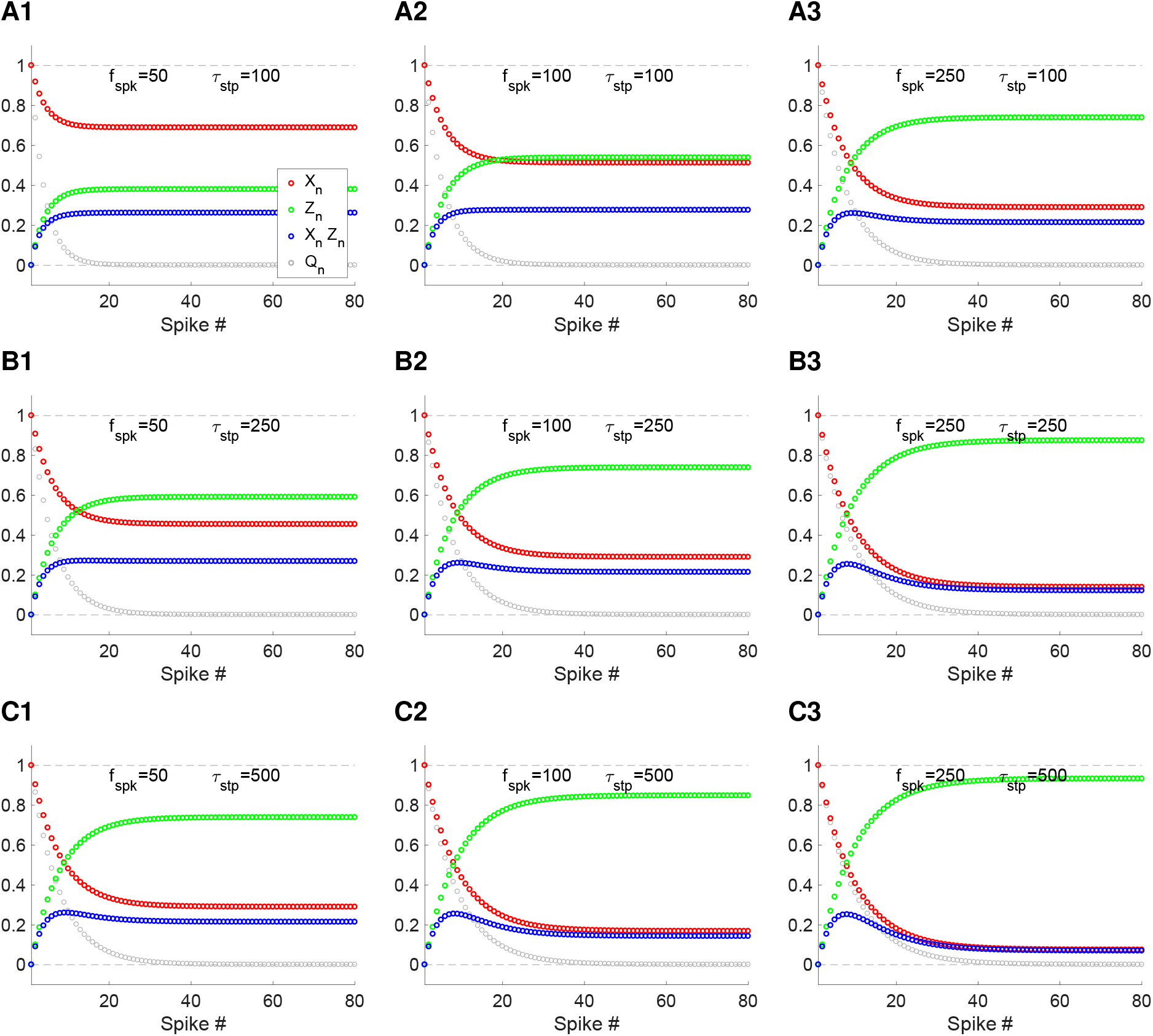
Low-, high- and band-pass filters in response to periodic presynaptic inputs in the presence of synaptic depression and facilitation: peak envelope dynamics. The evolution of the peak sequences *X_n_* (depression, red) and *Z_n_* (facilitation, green), respectively are governed by eqs. (24)–(26) and the sequence *Qn* = *Q*(*a_stp_, τ_stp_*)^*n*−1^ (light gray) is given by eq. (26). We used the same parameter values for depression and facilitation: *τ_dep_* = *τ_fac_* = *τ_stp_* and *a_d_* = *a_f_* = *a_stp_* = 0.1. **A.** *τ_stp_* = 100. **A1.***f_spk_* = 50. **A2.***f_spk_* = 100. **A3.***f_spk_* = 250. **B.***τ_stp_* = 250. **B1.***f_spk_* = 50. **B2.***f_spk_* = 100. **B3.***f_spk_* = 250. **C.***τ_stp_* = 500. **C1.***f_spk_* = 50. **C2.***f_spk_* = 100. **C3.***f_spk_* = 250. We used the folowing additional parameter values: *x*_∞_ = 1 and *z*_∞_ = 0.

The effective time scale of the sequence *Q_n_* = *Q^n^*^−1^ can be quantified by calculating the approximated time it takes for *Q_t_* (where the index *n* is substituted by *t*) to decrease from *Q*_1_ = 1 to 0.37 (decay by 63 % of the total decay range) and multiply this number by Δ_*spk*_. This yields

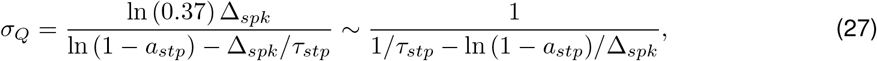

where *σ_Q_* is expressed in decimal numbers and has units of time. The time scales *σ_dep_* and *σ_fac_* for the sequences *X_n_* and *Z_n_* are obtained by substituting *τ_stp_* and *a_stp_* by *τ_dep_* and *a_d_* (*X_n_*) and by *τ_fac_* and *a_f_* (*Z_n_*), respectively. These time scales quantify the time it takes for *X_n_* to decrease from *X*_1_ to 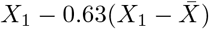 and for *Z_n_* to increase from *Z*_1_ to 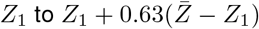, respectively.

#### 3.3.2 Additional properties of the depression and facilitation temporal LPFs and HPFs

The properties of the temporal LPFs and HPFs generated by *X_n_* and *Z_n_* are primarily dependent on the properties of the corresponding functions *Q*. For each value of *n*, *Q^n^*^−1^ is an increasing function of *Q* and for each fixed value of *Q*, *Q^n^*^−1^ is a decreasing function of *n*. Together, the larger *Q*, the larger the sequence *Q_n_* = *Q^n^*^−1^ and the slower *Q_n_* = *Q^n^*^−1^ converges to zero. From eq. (26), all other parameters fixed, *Q^n^*^−1^ decreases slower the smaller Δ_*spk*_ (the larger *f_spk_*) (compare Fig. 3 columns 1 to 3), the larger *τ_stp_* (compare Fig. 3 rows 1 to 3) and the smaller *a_stp_* (not shown in the figure). An extended analysis of the dependence of *Q*(*a_stp_, τ_stp_*) on both parameters can be found in Fig S1.

While the dynamics of *Q*(*a_stp_, τ_stp_*), *X_n_* and *Z_n_* depend on the quotient Δ_*spk*_/τ_*stp*_ (the two interacting time scales for the single events), the long-term time scales (*σ_Q_* = *σ_dep_, σ_fac_*) depend on these quantities in a more complex way (Fig.5-A). For the limiting case Δ_*spk*_ → ∞, *σ_dep/fac_ → τ_dep/fac_*. For the limiting case Δ*spk* → 0, *σ_dep/fac_* → 0. For values of Δ*spk* in between (see Appendix B), *σ_dep_* and *σ_fac_* decrease between these extreme values (*dσ_dep/fac_/d*Δ_*spk*_ < 0). The dependence of *σ_dep_* and *σ_fac_* with *τ_dep_* and *τ_fac_* follows a different pattern since (*dσ_dep/fac_/dτ_dep/fac_* > 0). Both *σ_dep_* and *σ_fac_* increase with *τ_dep_* and *τ_fac_*, respectively. The dependence of *σ_dep_* and *σ_fac_* with *a_d_* and *a_f_* follows a similar pattern (*dσ_d/f_ /da_d/f_* > 0). Details for these calculations are presented in the Appendix B.

One important feature is the dependence of the sequences *X_n_* and *Z_n_* on Δ_*spk*_ for fixed values of *τ_dep/fac_*. This highlights the fact the depression/facilitation-induced history-dependent filters are also frequency-dependent. A second important feature is that multiple combinations of *τ_dep/fac_* and Δ_*spk*_ give rise to the same sequence *X_n_* and *Z_n_*, which from eqs. (7)–(8) depend on the ratios

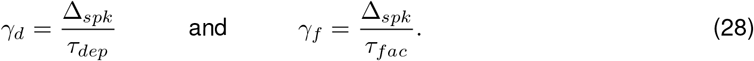

Constant values of *γ_d_* and *γ_f_* generate identical sequences *X_n_* and *Z_n_*, respectively, which will be differently distributed in the time domain according to the rescaling provided by Δ_*spk*_, reflecting the long-term time scales *σ_dep/fac_*. This degeneracy also occurs for the steady state values 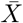 and 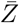 (9)-(10), generating the 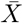 - and 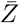-profiles (curves of 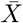 and 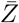 as a function of the input frequency 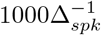). This type of degeneracies may interfere with the inference process of short-term dynamics from experimental data. A third important feature is that the update values *a_d_* and *a_f_* that operate at the single events contribute to the long-term time scale for the filters and do not simply produce a multiplicative effect on the filters uniformly across events.

#### 3.3.3 Descriptive envelope model for short-term dynamics in response to periodic presynaptic inputs

The relative simplicity of the DA model allows for the analytical calculation of the temporal patterns *X_n_* and *Z_n_* (24)-(26) (for constant values of Δ_*spk*_) and the subsequent analytical calculation of the (long-term) time scales *σ_dep_* (LPF) and *σ_fac_* (HPF) (27) in terms of the single event time constants *τ_dep_* and *τ_fac_*, respectively. Here, we develop an alternative approach for the computation of the LPF and HPF time constants, which is applicable to both a more general class of STP-mediated LPFs and HPFs, generated by more complex models for which we have no analytical expressions available, and to data collected following the appropriate stimulation protocols. This model is descriptive, as opposed to mechanistic, in the sense that it consists of functions that capture the shapes of the temporal LPFs and HPFs, but the model does not explain how these temporal filters are generated in terms of the parameters governing the dynamics of the single events, in contrast to the DA model.

We explain the basic ideas using data generated by the DA model. We then use this approach for the MT model in the supplementary material.

The shapes of the temporal LPFs and HPFs suggest an exponential-like decay to their steady states (e.g., Fig. 1). For the DA model this can be computed analytically following a similar approach to the one developed in Section 3.1 (for the synaptic dynamics) and extend the peak sequences (24)-(25) to the continuous domain

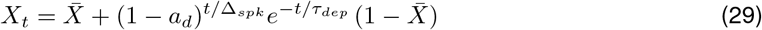

and

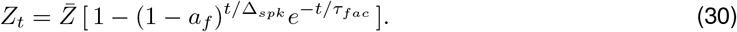

We assume exponential decay and define the following envelope functions

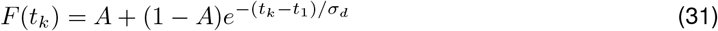

and

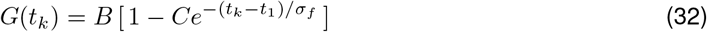

for the LPF and HPF, respectively. The parameters *σ_d_* and *σ_f_* are the filter time scales. We use a different notation than in the previous section to differentiate these time scales from the ones computed analytically for the DA model.

The parameters of the descriptive model (31)-(32) can be computed from the graphs of peak sequences (e.g., *X_n_* and *Z_n_* for the DA model or *R_n_* and *u_n_* for the MT model) by matching the initial values (e.g., *F* (*t*_1_) = *X*_1_ and *G*(*t*_1_) = *Z*_1_), the steady steady state values e. g., (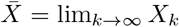 and 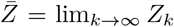) and the intermediate values (*t_c_, X_c_*) and (*t_c_, Z_c_*) chosen to be in the range of fastest increase/decrease of the corresponding sequences (~50% of the gap between the initial and steady state values). This gives

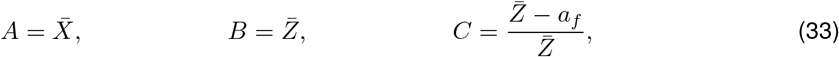

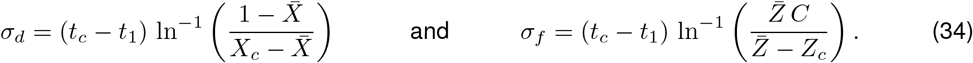

Fig. 1 (solid) shows the plots of *F* (red), *G* (green) and *H* = *FG* (blue) superimposed with the 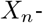, 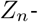 and Δ*S_n_*-sequence values. The error between the sequences and the envelope curves (using a normalized sum of square differences) is extremely small in both cases, consistent with previous findings [75]. For the DA model, *σ_dep_* and *σ_fac_* are well approximated by *σ_d_* and *σ_f_*, respectively.

For *f_spk_* → 0, both *X_n_* and *Z_n_* are almost constant, since the *x*(*t*) and *z*(*t*) have enough time to recover to their steady state values before the next input spike arrives, and therefore *σ_d_* ≫ 1 and *σ_f_* ≫ 1. In contrast, for *f_spk_* ≫ 1, *x*(*t*) and *z*(*t*) have little time to recover before the next input spike arrives and therefore they rapidly decay to their steady state values. In the limit of *f_spk_* → ∞, *σ_d_* = *σ_f_* = 0. In between these two limiting cases, *σ_d_* and *σ_f_* are decreasing functions of *f_spk_* (Figs. 4-A1 and -A2). For fixed-values of *f_spk_*, both *σ_d_* and *σ_f_* are increasing functions of *τ_dep_* and *τ_fac_*, respectively. These results are consistent with the analytical results described above.

**Figure 4:**
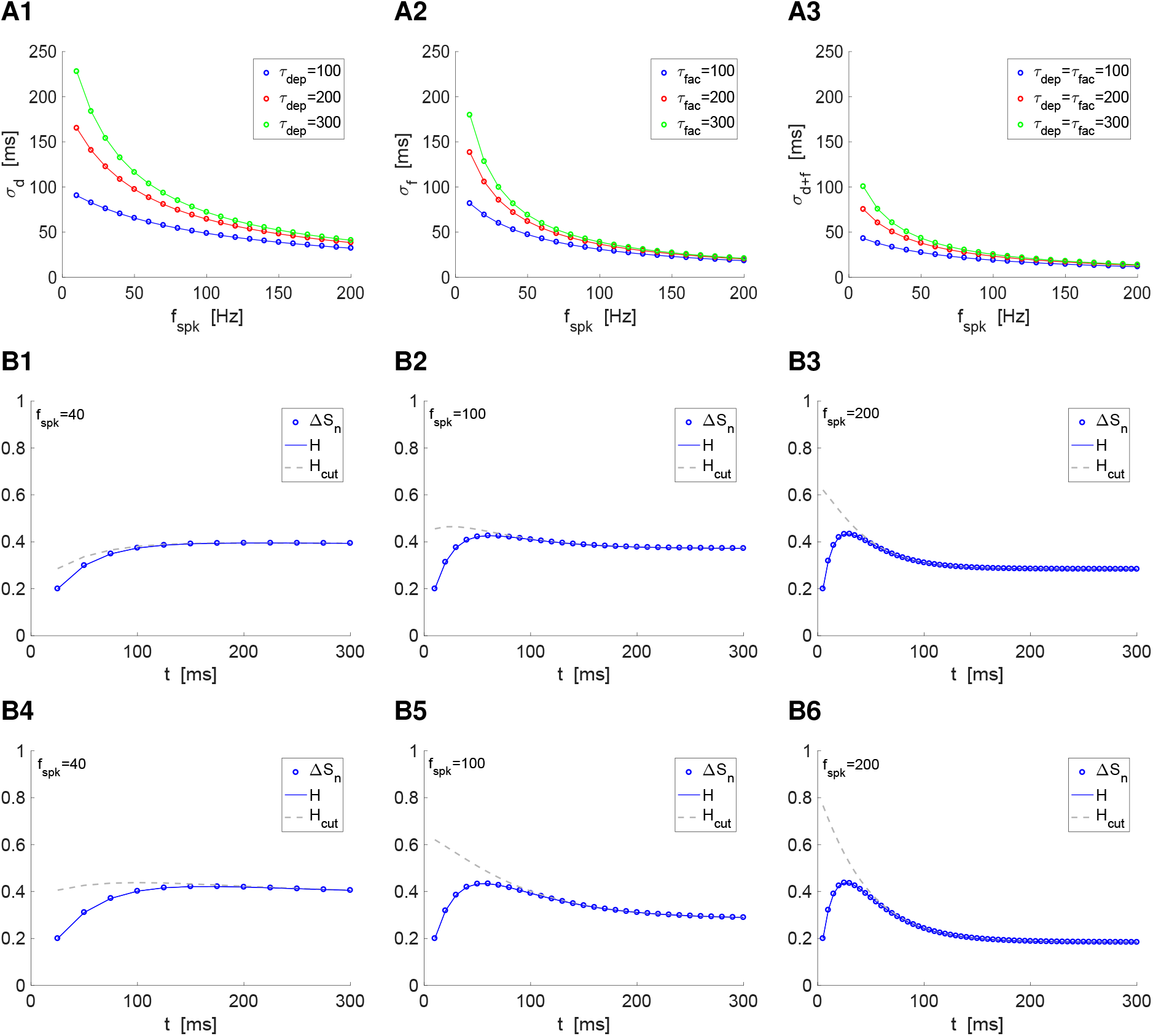
The time scales for the peak envelope responses *X_n_* and *Z_n_* to periodic presynaptic spikes and Δ*S_n_* temporal filters (*σ_d_*,*σ_f_* and *σ_d_*_+*f*_) depend on the interplay of *τ_dep_*,*τ_fac_* and *f_spk_* (or Δ_*spk*_). **A.** Dependence of *σ_d_*, *σ_f_* and *σ_d_*_+*f*_ with the presynaptic input frequency *f_spk_*. **A1.***σ_d_* for *a_dep_* = 0.1. **A2.***σ_f_* for *a_fac_* = 0.2. **A3.***σ_d_*_+*f*_ computed using eq. (107) from *σ_d_* and *σ_f_* in panels A1 and A2. **B.** The contribution of the combined time scale *σ_d_*_+*f*_ increases with *f_spk_*. The function *H*(*t_k_*) is given by (39) and the function *H_cut_* consists of the three first terms in (39). **B1-B3.***τ_dep_* = *τ_fac_* = 100. **B4-B6.***τ_dep_* = *τ_fac_* = 200. We used the simplified model (4)-(6) and the formulas (34) for the (simplified) descriptive model, together with (7)-(10) and the following parameter values: *a_d_* = 0.1, *a_f_* = 0.2, *x*_∞_ = 1, *z*_∞_ = 0.

**Figure 5:**
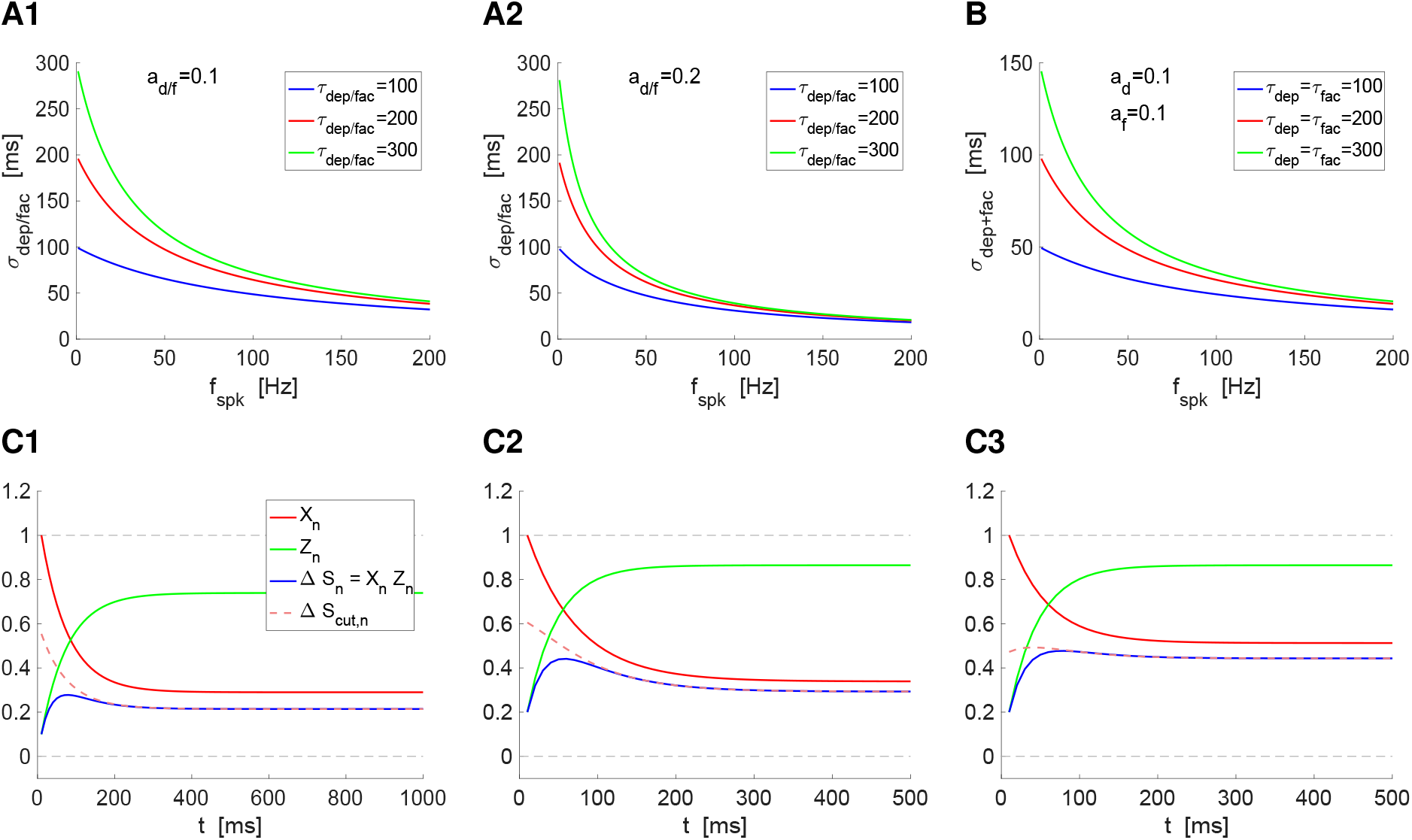
The time scales for the peak envelope responses *X_n_* and *Z_n_* to periodic presynaptic spikes and Δ*S_n_* = *X_n_Z_n_* temporal filters (*σ_dep_*,*σ_fac_* and *σ_dep_*_+*fac*_) depend on the interplay of *τ_dep_*,*τ_fac_* and *f_spk_* (or Δ_*spk*_). The evolution of the peak sequences *X_n_* (depression, red) and *Z_n_* (facilitation, green), respectively are governed by eqs. (24)–(26) and the sequence *Q_n_* = *Q*(*a_stp_, τ_stp_*)^*n*−1^ (light gray) is given by eq. (26). **A.** Dependence of the (long-term) temporal filter time constants *σ_dep_* and *σ_fac_* with the presynaptic input frequency *f_spk_* and (short-term) time constants for the single events *τ_dep_* and *τ_fac_*. We used eq. (27) with *τ_stp_* and *a_stp_* substituted by *τ_dep/fact_* and *a_d/f_*, respectively. **A1.***a_d_* = *a_f_* = 0.1. **A2.***a_d_* = *a_f_* = 0.2. **B.** Dependence of the the (long-term) temporal filter time constants *σ_dep_*_+*fac*_ with the presynaptic input frequency *f_spk_* and (short-term) time constants for the single events *τ_dep_* and *τ_fac_*. We used eq. (27). **C.** Comparison between the filters produced by the “cut” sequence Δ*S_cut,n_* (light coral) and the sequence Δ*S_n_* (blue) for representative parameter values. **A1.***a_d_* = 0.1, *a_f_* = 0.1, *τ_dep_* = 250 and *τ_fac_* = 250. **A2.***a_d_* = 0.1, *a_f_* = 0.2, *τ_dep_* = 200 and *τ_fac_* = 250. **A3.***a_d_* = 0.1, *a_f_* = 0.2, *τ_dep_* = 100 and *τ_fac_* = 250. We used the following additional parameter values: *x*_∞_ = 1 and *z*_∞_ = 0.

### 3.4 Emergence of temporal band-pass filters for Δ*S_n_* = *X_n_Z_n_*: interplay of depression and facilitation

Under certain conditions, the interplay of depression and facilitation generates temporal band-pass filters (BPFs) in response to periodic inputs for (Fig. 1-A), which are captured by the sequence Δ*S_n_* = *X_n_Z_n_* (Fig. 1-A2 and -C2). BPFs represent an overshoot for the sequence Δ*S_n_* (they require 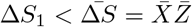 and the existence of a spike index *m* such that 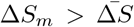). This in turn requires that the two time constants *τ_dep_* and *τ_fac_* are such that they create the appropriate balance between the two temporal filter time constants *σ_dep_* and *σ_fac_* (or *σ_d_* and *σ_f_* when using the descriptive model) to support a BPF.

For the parameter values in Figs. 3, Δ*S_n_* BPFs emerge and become more prominent as the input frequency *f_spk_* increases for fixed values of *τ_dep_* = *τ_fac_*(= *τ_stp_*). This results from both the dependence of the *X_n_* and *Z_n_* time constants *σ_dep/fac_* on the single event time constants *τ_dep/fac_* and the fact that 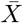 decreases and 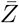 increases with increasing values of *f_spk_*. Fig. 6 further illustrates that temporal Δ*S_n_* BPFs (panels C, D, E) provide a transition mechanism from LPFs for low enough input frequencies (panels A and B) to HPFs for high-enough frequencies (panel F, which is strictly not a HPF, but it is effectively so for the time scale considered). Fig. 3 also illustrates that for fixed values of *f_spk_*, the Δ*S_n_* BPFs emerge and become more prominent as *τ_dep_* = *τ_fac_* increases. This is a consequence of the dependence of the *X_n_* and *Z_n_* time constants *σ_dep/fac_* on the single event time constants *τ_dep/fac_* and the fact that 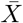 decreases with increasing values of *τ_dep_* and 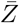 increases with increasing values of *τ_fac_*. Fig. 7-A summarizes the dependences of 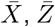 and the quotient between *σ_dep_* and *σ_fac_*. Because of the dependence of *X_n_* and *Z_n_* on the quotients Δ_*spk*_/τ_*dep/fac*_, Δ*S_n_* BPFs can be generated by increasing values of *τ_dep_*, *τ_fac_* or both (not shown).

**Figure 6:**
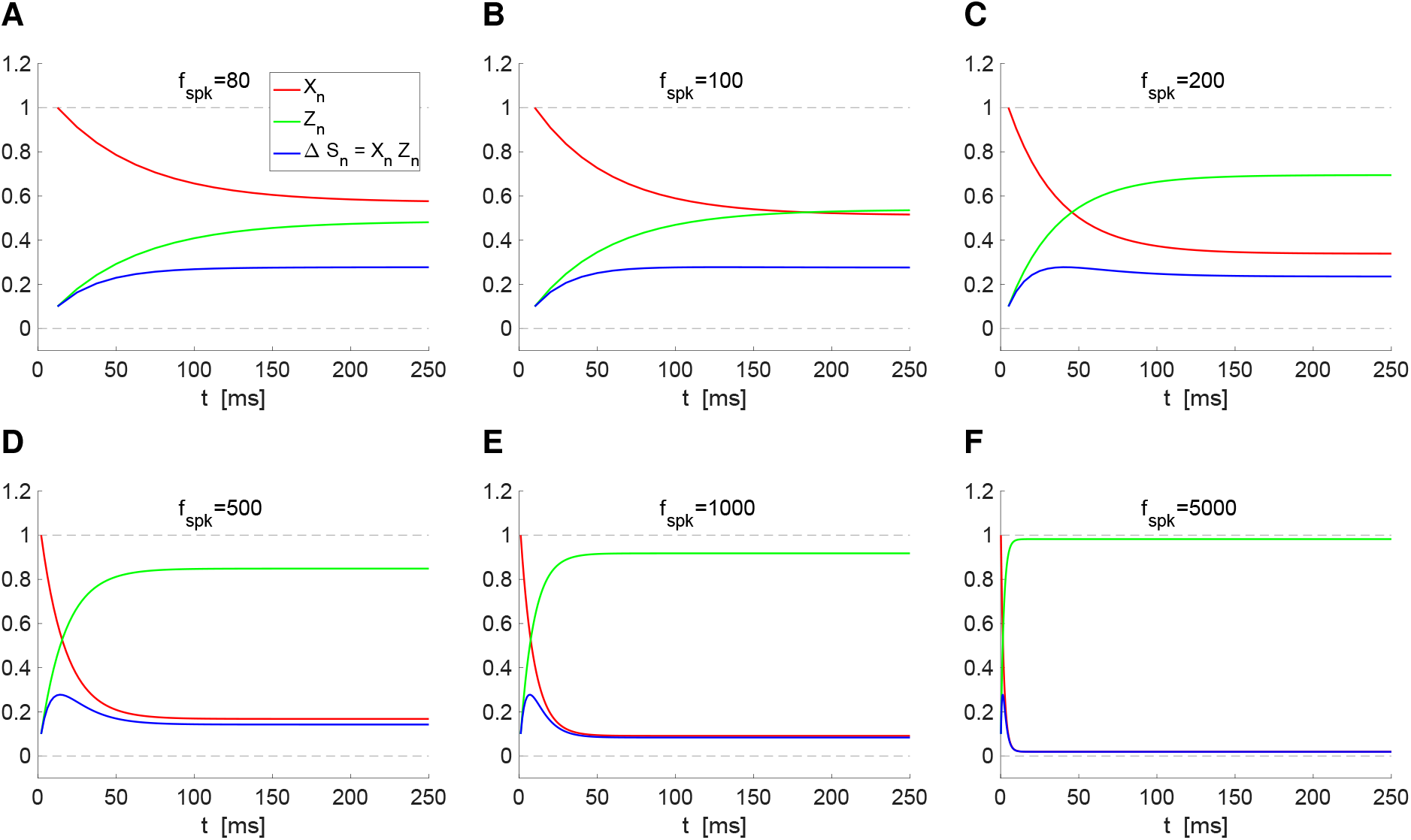
Transition from temporal high- to low-pass filters as the presynaptic frequency increases via a temporal band-pass filtering mechanism for fixed values of the synaptic depression and facilitation time constants. The evolution of the peak sequences *X_n_* (depression, red) and *Z_n_* (facilitation, green), respectively are governed by eqs. (24)–(26). We used the same parameter values for depression and facilitation: *τ_dep_* = *τ_fac_* = 100 and *a_d_* = *a_f_* = 0.1. **A.***f_spk_* = 80. **B.***f_spk_* = 100. **C.** *f_spk_* = 200. **D.***f_spk_* = 500. **E.***f_spk_* = 1000. **F.***f_spk_* = 5000. We used the following additional parameter values: *x*_∞_ = 1 and *z*_∞_ = 0.

**Figure 7:**
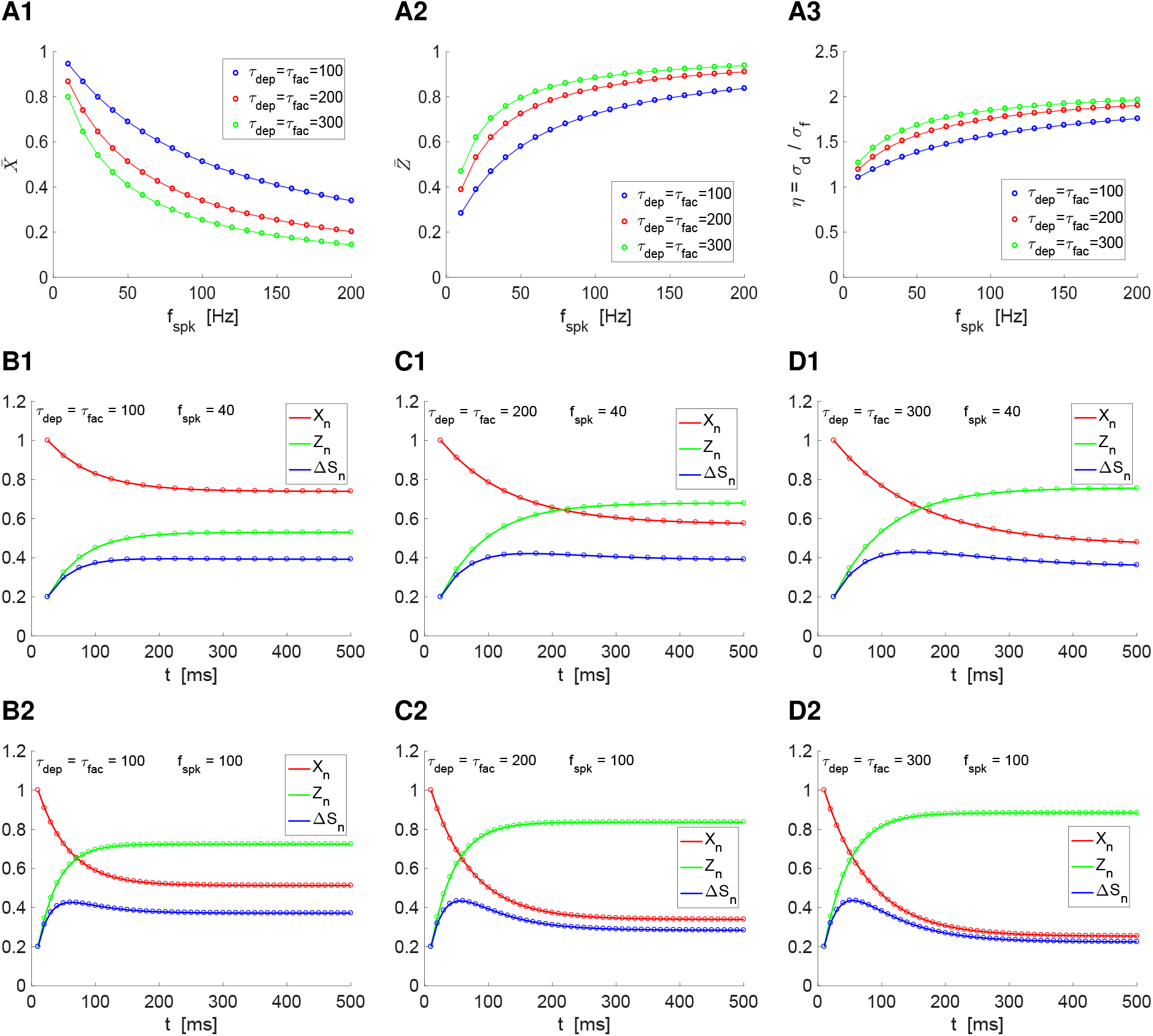
Temporal band-pass filters generated as the result of the multiplicative interaction of temporal low- and high-pass filters: Peak envelope responses to periodic presynaptic inputs. **A.** Dependence of *η* = *σ_dep_/σ_fac_*, 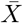 and 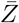 with the presynaptic input frequency *f_spk_*. We used formulas (9)-(10) together with (34) and the following parameter values: *a_d_* = 0.1, *a_f_* = 0.2, *x*_∞_ = 1, *z*_∞_ = 0. **B-D.** Variables *X_n_*, *Z_n_*, and Δ*S_n_* for different presynaptic input frequencies and combinations of *τ_dep_* and *τ_fac_* (see legend). **B1.***τ_dep_* = *τ_fac_* = 100 and *f_spk_* = 40. **C1.***τ_dep_* = *τ_fac_* = 200 and *f_spk_* = 40. **D1.***τ_dep_* = *τ_fac_* = 300 and *f_spk_* = 40. **B2.***τ_dep_* = *τ_fac_* = 100 and *f_spk_* = 100. **C2.***τ_dep_* = *τ_fac_* = 200 and *f_spk_* = 100. **D2.***τ_dep_* = *τ_fac_* = 300 and *f_spk_* = 100.

From a purely geometric perspective (devoid of any biophysical meaning), it is expected that the product of two exponential-like decaying functions, one increasing and the other decreasing, has a peak in certain parameter regimes (see Appendix E). By design, the geometric/dynamic mechanism described in the Appendix E is based on the assumption that all the parameters are free and independent. However, from eqs. (9)–(10) and (33)–(34) the geometric parameters that describe the LPF and HPF are not independent and therefore it is not clear how Δ*S_n_* BPF are generated and how they depend on the single event time constants *τ_dep_* and *τ_fac_*.

### 3.5 *ΔS_n_*-band pass filters require a third (emergent) time scale whose contribution is independent from the low- and high-pass filters’ time scales

The linear one-dimensional dynamics for both depression and facilitation at the single event level generate linear one-dimensional discrete dynamics at the (long-term) temporal (low- and high-pass) filters level where the long-term time constants (*σ_dep_* and *σ_fac_*) depend on the short-term time constants (*τ_dep_* and *τ_fac_*) and the input frequency (Δ_*spk*_). From the discrete dynamics point of view, the temporal BPFs obtained as the product of the temporal LPF and HPF are considered overshoots where the sequence evolves by peaking at a higher value than the steady-state (without exhibiting damped oscillations). Over-shoots require at least two-dimensional dynamics (generated by a difference equation where each value of the resulting sequence depends on the previous two) with the appropriate time scales. In [75] we showed that in certain circumstances the generation of temporal BPFs requires three-dimensional dynamics. Here we investigate how the time scales giving rise to temporal BPF depend on the time scales of the temporal LPF and HPF and these of the corresponding single events.

#### 3.5.1 Mechanistic DA model

From (24)-(26),

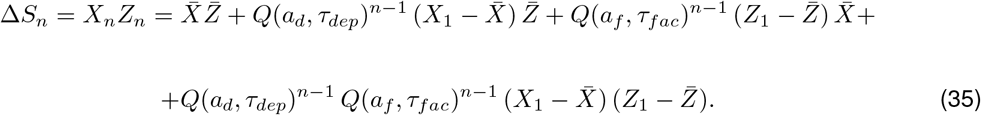

The dynamics of the three last terms in eq. (35) are governed by *Q*(*a_d_, τ_dep_*)^*n*−1^, *Q*(*a_f_, τ_fac_*)^*n*−1^ and [*Q*(*a_d_, τ_dep_*) *Q*(*a_f_, τ_fac_*)]^*n*−1^, respectively, and 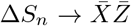 as *n* → ∞. The first two are given eq. (26) with *τ_stp_* substituted by *τ_dep_* and *τ_fac_*, and *a_stp_* substituted by *a_d_* and *a_f_*, accordingly. The corresponding time scales are given by eq. (27) with the same substitutions. The last one is given by

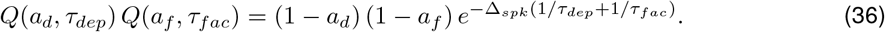

The product [*Q*(*a_d_, τ_dep_*) *Q*(*a_f_, τ_fac_*)]^*n*−1^ cannot be expressed in terms of a linear combination of *Q*(*a_d_, τ_dep_*)^*n*−1^ and *Q*(*a_f_, τ_fac_*)^*n*−1^ and therefore the four terms in (35) are linearly independent.

The time scale associated to the fourth terms in (35) can be computed from (36) (as we did before) by calculating the time it takes for [*Q*(*a_d_, τ_dep_*) *Q*(*a_f_, τ_fac_*)]^*n*−1^ to decrease from [*Q*(*a_d_, τ_dep_*) *Q*(*a_f_, τ_fac_*)]^*n*−1^ from 1 (is value for *n* = 1) to 0.37 and multiply this number by Δ_*spk*_. This yields

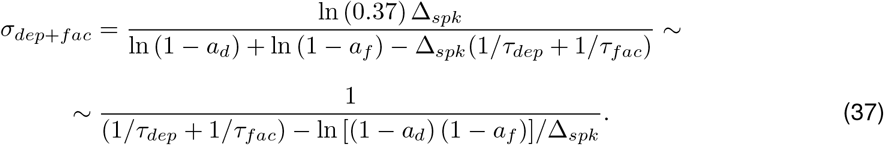

This long-term time scale has similar properties as *σ_dep_* and *σ_fac_*. In particular, for *f_spk_* → 0, *σ*_*dep*+*fac*_ → τ_*dep*_τ_*fac*_/(*τ_dep_* + *τ_fac_*). For *f_spk_* → ∞, *σ*_*dep*+*fac*_ → 0. For values of *f_spk_* in between, *σ_dep_*_+*fac*_ decrease between these extreme values. This is illustrated in Fig. 5-B along the time other two times scales, *σ_dep_* and *σ_fac_* (Fig. 5-B).

To say that the three time scales (*σ_dep_*, *σ_fac_* and *σ_dep_*_+*fac*_) are independent is equivalent to state that the dynamics is three dimensional, while the dynamics of the depression and facilitation sequences are one-dimensional. It also means that erasing one of the terms in Δ*S_n_* is equivalent to projecting the three-dimensional signal into a two-dimensional space with the consequent loss of information if it does not provide a good approximation to the original signal, and this loss of information may in principle be the loss of the band-pass filter (overshoot). On the other hand, two-dimensional systems are able to produce overshoots. So the question arises of whether the signal Δ*S_n_* without the last term (that combines the time scales of the two filters *X_n_* and *Z_n_*) preserves the temporal band-pass filter and, if yes, under what conditions.

In order to test the necessity of this third time scale for the generation of temporal band-pass filters, we consider the “cut” sequence

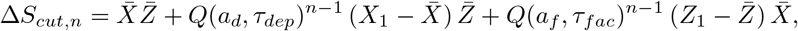

where 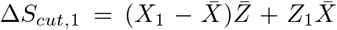 and 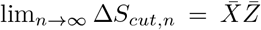. For *a_d_* = *a_f_* and *τ_dep_* = *τ_fac_*, *Q*(*a_d_, τ_dep_*) = *Q*(*a_f_, τ_fac_*), and therefore Δ*S_cut,n_* has the same structure as *X_n_* and *Z_n_*, and therefore Δ*S_cut,n_* cannot generate a temporal band-pass filter regardless of the value of Δ_*spk*_. This includes the examples presented in Fig. 3. For other parameter values, standard calculations show that the parameter ranges for which Δ*S_cut,n_* shows a peak are very restricted and when it happens, they rarely provide a good approximation to the temporal band-pass filter exhibited by Δ*S_n_*. Fig. 5-C illustrates this for a number of representative examples.

#### 3.5.2 Descriptive envelope model

Here we address similar issues using the descriptive models described in the previous section. We consider the function

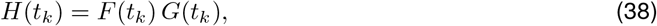

which approximates the behavior of Δ*S_n_*. From (31)-(32),

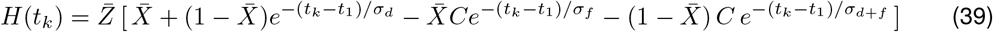

where

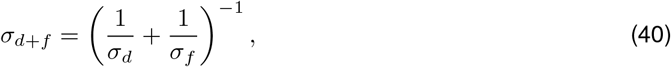

and *σ_d_* and *σ_f_* are given by (34). Note that *σ_d_*, *σ_f_* and *σ_d_*_+*f*_ are different quantities from *σ_dep_*, *σ_fac_* and *σ_dep_*_+*fac*_ discussed above. Note also that, formally, the dependences of the third time scales (*σ_dep_*_+*fac*_ and *σ_d_*_+*f*_) on the corresponding LPF and HPF time scales are different. In contrast to the analytical expression for the DA model, the third time scale for the descriptive model (*σ_d_*_+*f*_) can be explicitely computed in terms of the time scales for the LPF and HPF.

Together, these results and the results from the previous section shows that while *X_n_* and *Z_n_* are generated by a 1D (linear) discrete dynamical systems (1D difference equations), Δ*S_n_* is generated by a 3D (linear) discrete dynamical system (3D discrete difference equation). Under certain conditions, a 2D (linear) discrete dynamical system will produce a good approximation. Δ*S_n_* is able to exhibit a band-pass filter because of the higher dimensionality of its generative model.

In order to understand the contribution of the combined time scale *σ_d_*_+*f*_, we look at the effect of cutting the fourth term in H (39). We call this function *H_cut_*. Fig. 4-B shows that *H_cut_* does not approximate Δ*S_n_* well during the transients (response to the first input spikes) and fails to capture the transient peaks and the temporal band-pass filter properties of Δ*S_n_*. This discrepancy between Δ*S_n_* (or H) and *H_cut_* is more pronounced for low than for high input frequencies. In fact 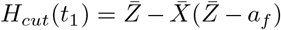, while *H*(*t*_1_) = *a_f_*. The question remains of whether there could be a 2D (linear) dynamical system able to reproduce the temporal filters for Δ*S_n_* with (emergent) time scales different from *σ_d_* and *σ_f_*. In other words, whether Δ*S_n_* can be captured by a function of the form

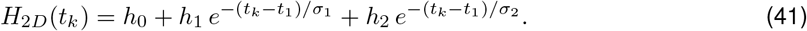

where *h*_0_, *h*_1_, *h*_2_, *σ*_1_ and *σ*_2_ are constants where by necessity,

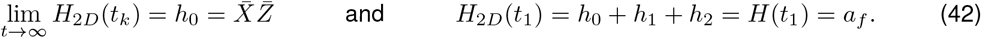

The fact that the function *H* is a sum of exponentials indicate that the answer is negative.

The dependence of the estimated time scales *σ_d_*, *σ_f_* and *σ_d_*_+*f*_ on the parameters *f_spk_*, *τ_dep_* and *τ_fac_* needs to be computed numerically. Our results are presented in Fig. 4-A, and are consistent with our previous results.

### 3.6 Interplay of short-term synaptic and cellular postsynaptic dynamics: temporal BPFs generated within and across levels of organization

Earlier models of synaptic dynamics consider the postsynaptic potential (PSP) peak sequence to be proportional to Δ*S_n_* = *X_n_Z_n_* [21, 63, 67]. This approach does not take into consideration the dynamic interaction between the synaptic function *S* and the postsynaptic membrane potential, particularly the membrane time scale. Subsequent models consider synaptic currents such as *I_syn_* in eq. (2). The synaptic function *S*, which controls the synaptic efficacy, obeys a first linear kinetic equation (see Appendix C). The presence of additional times scales further in the line (e.g., membrane potential time scale) gives rise to the phenomenon of temporal summation and the associated HPF in response to periodic synaptic inputs (see Fig. 2-B), which interacts with the LPFs and HPFs associated with synaptic depression and facilitation, respectively. The resulting PSP temporal filters reflect these interactions and therefore are expected to depart from the proportionality relationship with the Δ*S_n_* filters.

Here we address these issues by following a dual approach. We first consider postsynaptic dynamics governed by eq. (3). We interpret the variable *S* as the postsynaptic membrane potential and the decay time *τ_dec_* as reflecting the membrane potential dynamics (Section 3.6.1). The simplified model has the advantage of being analytically solvable and it allows us to understand the effects of the temporal filtering (depression, facilitation and summation) time scales in terms of the single event time constants (*τ_dep_*, *τ_fac_* and *τ_dec_*). Then, we consider the more biophysically realistic approach by using eqs. (1)–(3) where *τ_dec_* is chosen to be relatively small, of the order of magnitude of the AMPA decay time (Section 3.6.2). In this model, the membrane time constant is *C/G_L_* and the interplay between the synaptic input and the postsynaptic cell is multiplicative (nonlinear). In both approaches, a systematic analysis of the generation and properties of temporal PSP filters would require the consideration of an enormous amount of cases given the increase in the number of building blocks and the consequent increase in the number of time scales involved. Therefore, in our study we consider a number of representative guided by mechanistic questions. For conceptual purposes, we also considered the case where rise times are non-zero (see eq. (91) in Appendix C.1).

#### 3.6.1 PSP temporal filters in the simplified model

We use eq. (3) with decay times reflecting the membrane potential time scales of postsynaptic cells. As mentioned above, in this simplified intermediate approach *S* is interpreted as the postsynaptic membrane potential. Our goal is to understand how the (global) time scales of the *S*-response patterns to periodic inputs depend on the time constant *τ_dec_* and the depression and facilitation time scales *τ_dep_* and *τ_fac_* through the Δ*S_n_* filter time scales *σ_dep_* and *σ_fac_*.

In the absence of depression and facilitation (*τ_dep_, τ_fac_* → 0), Δ*S_n_* is constant across cycles and *S* generates temporal summation HPFs as described in Section 3.1 (Fig. 2-B) with *σ_sum_* = *τ_dec_*. We use the notation 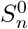 for the corresponding peak sequences. In the presence of either depression or facilitation, the update Δ*S_n_* is no longer constant across cycles and therefore, the STP LPFs and HPFs interact with the summation HPFs.

By solving the differential equation (3) where *S* is increased by Δ*S_n_* at the arrival of each spike (*t* = *t_n_*, *n* = 1*,…, N_spk_*) one arrives to the following discrete linear differential equation for the peak sequences in terms of the model parameters

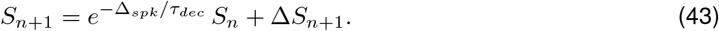

The solution to eq. (43) is given by the following equation involving the convolution between the STP input Δ*S_n_* and an exponentially decreasing function

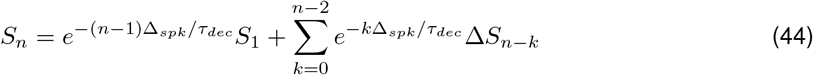

with *S*_1_ = Δ*S*_1_ = *a_f_*. The evolution of *S_n_* is affected by the history of the STP inputs Δ*S_n_* weighted by an exponentially decreasing function of the spike index and a coefficient

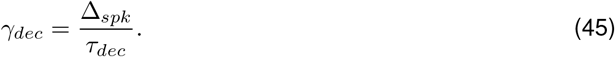

The steady state is given by

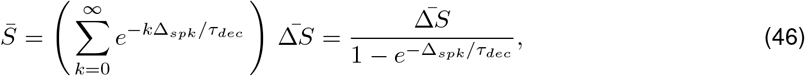

where 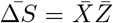 given by (9) and (10). Note that Eq. (19) is a particular case of eq. (46) when Δ*S_n_* is a constant sequence (no STP).

Both *S_n_* and 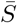 depend on Δ_*spk*_ and the time constants *τ_dep_*, *τ_fac_* and *τ_dec_* through the quotients (28) and (45). Therefore, here we consider temporal patterns for a fixed-value of Δ_*spk*_. The temporal patterns for other values of Δ_*spk*_ will be temporal compressions, expansions and height modulations of these baseline patterns. The values of *τ_dep_*, *τ_fac_* and *τ_dec_* used in our simulations should be interpreted in this context.

For the limiting case *τ_dec_* → 0, *S_n_* → Δ*S_n_* for 1 = 2*,…, N_spk_*. The *S*-temporal filter reproduces (is equal to or a multiple of) the Δ*S_n_* as in [21, 63, 67]. For the limiting case 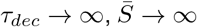 reflecting the lack of convergence of the sum in eq. (44). As the presynaptic spike number increases, the *S*-temporal filter reproduces the *S*^0^-temporal filter since the 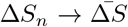. In the remaining regimes, changes in *τ_dec_* affect both the steady state and the temporal properties of *S* in an input frequency-dependent manner as the *S*-temporal filter transitions between the two limiting cases. Here we focus on the temporal filtering properties. The former will be the object of a separate study.

##### Emergence of *S_n_* temporal BPFs: Interplay of synaptic depression (LPF) and postsynaptic summation (HPF)

In the Δ*S_n_* facilitation-dominated regime (Fig. 8-A), the PSP *S_n_* patterns result from the interaction between two HPFs, the Δ*S_n_* facilitation and the 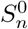 summation ones. The filter time constant increases with increasing values of *τ_dec_* reflecting the dependence of the summation (global) time constant *σ_sum_* with *τ_dec_*.

**Figure 8:**
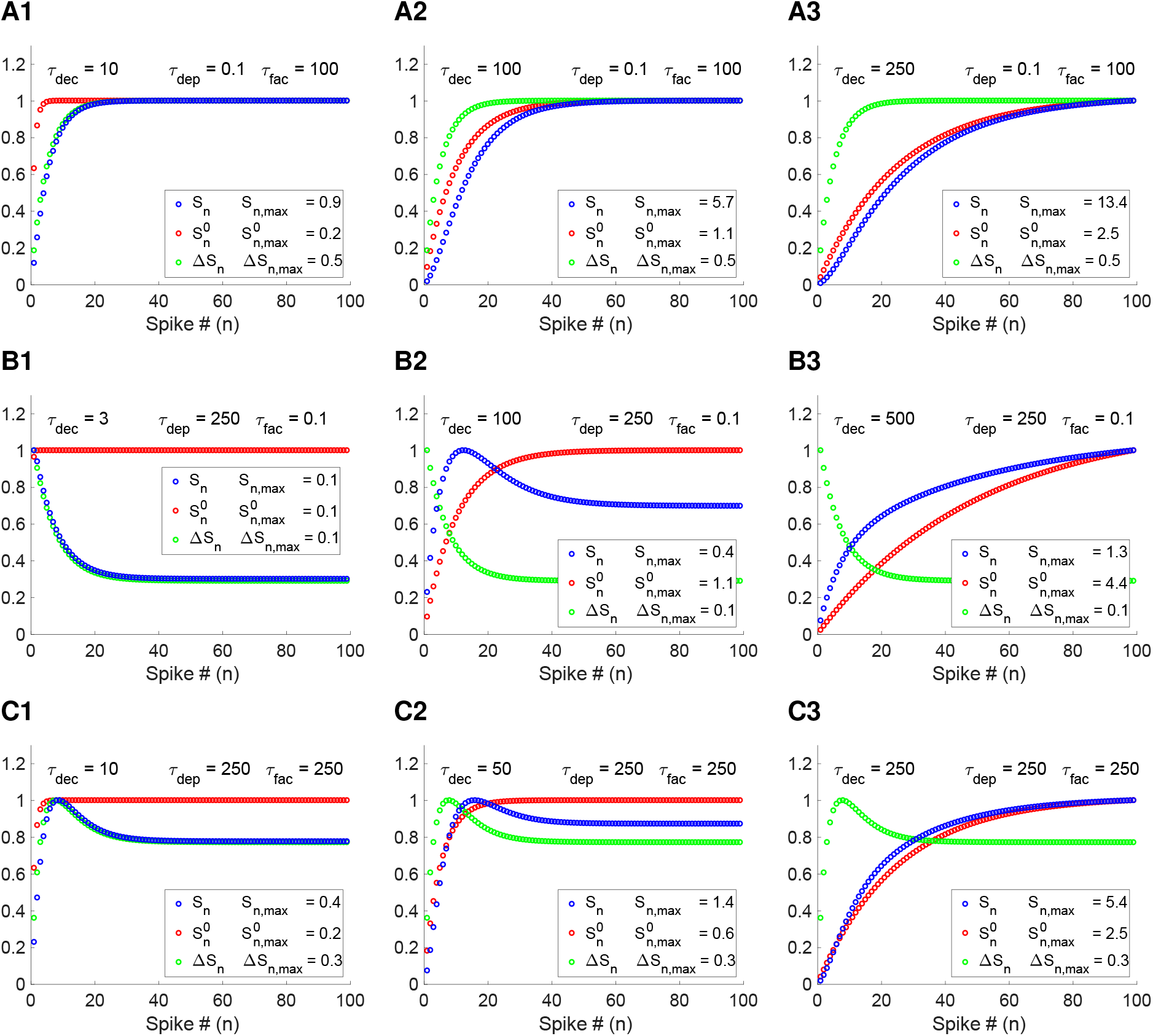
Temporal *S*-filters in response to periodic presynaptic inputs in the presence of STP. We used the DA model for synaptic depression and facilitation to generate the sequences Δ*S_n_* and eq. (3) to generate the *S_n_* sequences. The 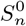 sequences were computed using eq. (3) with a constant value of Δ*S_n_* = Δ*S* = *a_fac_*. The curves are normalized by their maxima *S_n,max_*, Δ*S_n,max_* and 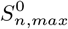. **A.** Facilitation-dominated regime. *S_n_* is a HPF (transitions between two HPFs). The time constant increases monotonically with *τ_dec_*. **B.** Depression-dominated regime. A *S_n_* BPF is created as the result of the interaction between the presynaptic (depression) LPF and the temporal summation HPF. **C.***S_n_* and Δ*S_n_* temporal BPFs peak at different times. We used the following parameter values: *a_d_* = 0.1, *a_f_* = 0.2, *x*_∞_ = 1, *z*_∞_ = 0. For visualization purposes and to compare the (global) time constants of the *S*, Δ*S* and *S*^0^ temporal filters, we present the *S_n_*, Δ*S_n_* and 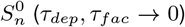 curves normalized by their maxima.

In the Δ*S_n_* depression-dominated regime (Fig. 8-B), the temporal PSP *S_n_* BPFs emerge as the result of the interaction between the Δ*S_n_* depression LPF and a 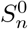 HPF for intermediate values of *τ_dec_* (Fig. 8-B2). *S_n_* BPFs are not present for small enough values of *τ_dec_* (Fig. 8-B1) since this would require Δ*S_n_* to be a BPF, and are also not present for large enough values of *τ_dec_* (Fig. 8-B3) since this would require 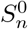 to be a BPF. As for the depression/facilitation BPFs discussed above, the *S_n_* BPFs are a balance between the two other filters and emerge as *S_n_* transitions in between them as *τ_dec_* changes.

##### Dislocation of the (output)*S_n_* temporal BPF from the (input)Δ*S_n_* temporal BPFs

In this scenario, a depression-facilitation Δ*S_n_* BPF is generated at the synaptic level and interacts with the summation 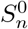 HPF (Fig. 8-C). The Δ*S_n_* BFP evokes a PSP *S_n_* BFP for low enough values of *τ_dec_* (Fig. 8-C1). As *τ_dec_* increases, the *S_n_* pattern transitions to the 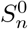 HPF (Fig. 8-C3). As this transition happens, the *S_n_* BPF moves to the right and increases in size before entering the summation-dominated HPF regime. While the PSP *S_n_* BPF is inherited from the synaptic regime, its structure results from the interplay of the synaptic BPF and PSP temporal summation.

#### 3.6.2 Biophysically realistic models reproduce the above PSP temporal filters with similar mechanisms

Here we test whether the results and mechanisms discussed above remain valid when using the more realistic, conductance-based model (1)-(3). Here *S* has its original interpretation as a synaptic function with relatively small time constants, consistent with AMPA excitatory synaptic connections. Because the interaction between the synaptic variable *S* and *V* are multiplicative, the model is not analytically solvable. The *V* temporal patterns generated by this model (Fig. 9) are largely similar to the ones discussed above (Fig. 8) and are generated by similar mechanisms described in terms of the interplay of the membrane potential time scale *τ_m_* (= *C/gL*) and the depression/facilitation time scales (*τ_dep_* and *τ_fac_*) through the (global) Δ*S_n_* filter time scales (*σ_dep_* and *σ_fac_*). Because *τ_dec_* is relatively small, *S* largely reproduces the temporal properties of the Δ*S_n_* pattern. As before, 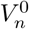 refer to the voltage response to presynaptic periodic inputs in the absence of STD (*S* is updated to Δ*S_n_* constant). Fig. 9-A illustrates the generation of *V* temporal BPFs as the result of the interaction between synaptic depression (LPF) and postsynaptic summation (HPF). Fig. 9-B illustrates the dislocation of postsynaptic BPF inherited from the synaptic input Fig. 9-A. We limited our study to realistic values of *τ_m_*.

**Figure 9:**
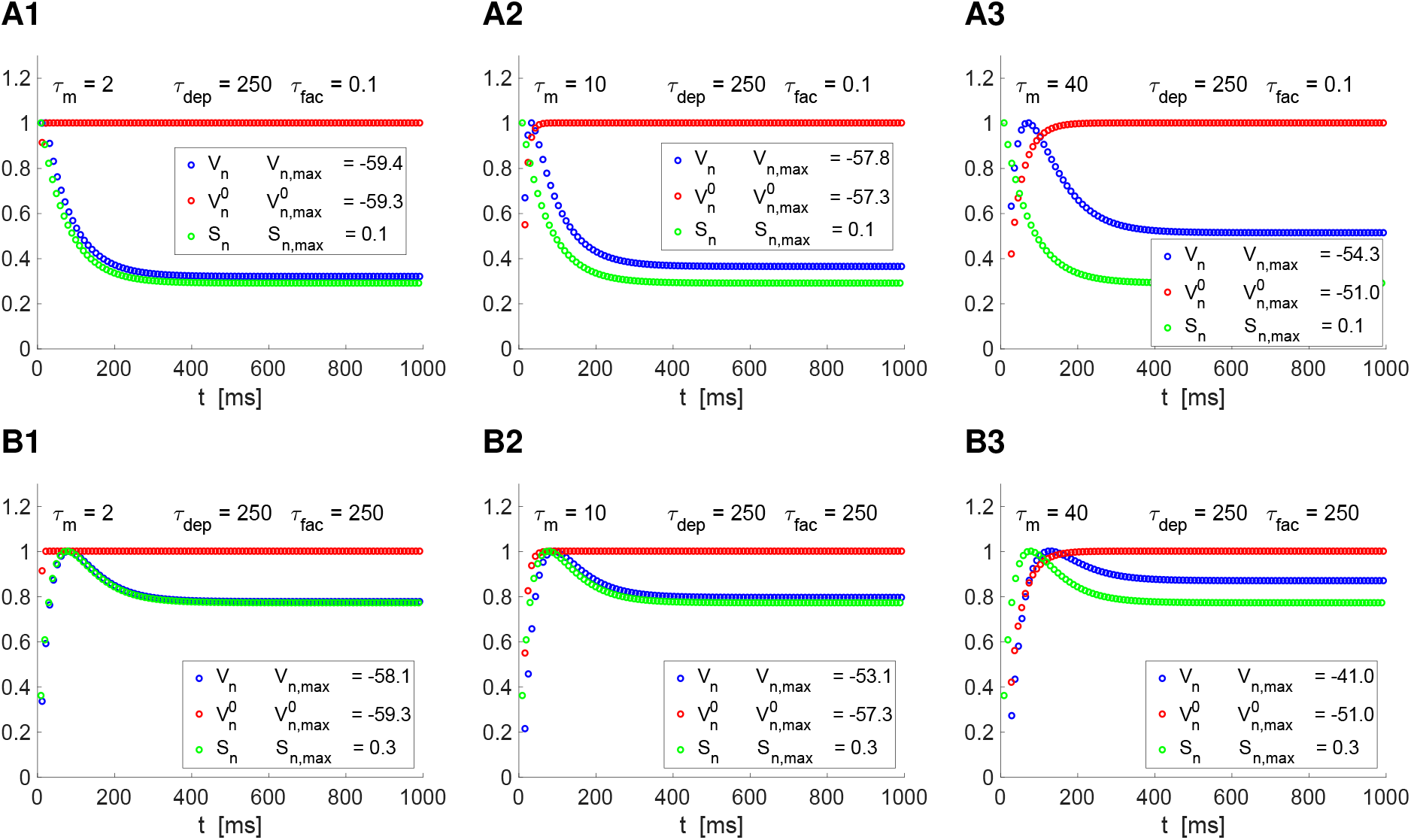
Temporal *S*-filters in response to periodic presynaptic inputs in the presence of STP. We used the DA model for synaptic depression and facilitation to generate the sequences Δ*S_n_*, eq. (3) (*τ_dec_* = 5) to generate the *S_n_* sequences, and the passive membrane equation (1)–(2) to generate the *V_n_* sequences. The 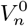 sequences were computed using eq. (3) with a constant value of Δ*S_n_* = Δ*S* = *a_fac_*. The curves are normalized by their maxima *S_n,max_*, §_*n,max*_ and 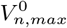. **B.** Depression-dominated regime. A *S_n_* BPF is created as the result of the interaction between the presynaptic (depression) LPF and the temporal summation HPF. **C.***V_n_* and *S_n_* temporal BPFs peak at different times. We used the following parameter values: *a_d_* = 0.1, *a_f_* = 0.2, *x*_∞_ = 1, *z*_∞_ = 0, *C* = 1, *E_L_* = −60, *E_syn_* = 0, *G_syn_* = 0.1.

#### 3.6.3 Attenuation of the filtering properties as the rise time increases

A conceptual question that arises in this context is whether and how *S* interacts with the membrane potential in the presence of longer rise times than the ones considered here. We did not take this into account, in geneeral since rise times for the type of synapses we use are assumed to be fast (typical for *AMPA* and also *GABA_A_*) for which the instantaneous approximation is justified. We illustrate the effect of longer rise times in Fig. S2 in the context of the DA model. We used presynaptic spikes of 1 ms width. (see explanation in Appendix C.1 and eq. (91)). In the first row of Fig. S2, we see a comparison of how the rise time affects *S* and *V* alone and in the second line we see panels of the same simulation above but in the presence of the DA model. As *τ_rise_* increases, the presynaptic spikes take longer and longer to increase. In the first row, we see that the steady-state is lowered by such an effect and in the second row, we see that the type of BPF from the DA models persists qualitatively but is suppressed quantitatively.

### 3.7 Interplay of multiple depression and facilitation processes with different time scales

In the models discussed so far both short-term depression and facilitation involve one time scale (*τ_dep_* and *τ_fac_*) that governs the evolution of the corresponding variables (*x* and *z*) in between presynaptic spikes. These single-event time scales are the primary component of the temporal filter time scales (*σ_dep_* and *σ_fac_*) that develop in response to the periodic repetitive arrival of presynaptic spikes.

Here we extend these ideas to include scenarios where depression and facilitation involve more than one time scale. We interpret this as the coexistence of more than one depression or facilitation process whose dynamics are governed by a single single-event time scale each. Similar to the standard model, the independent filters that develop in response to the presynaptic spike train inputs interact to provide an input to the synaptic dynamics. In principle, this interaction may take various forms. Here, for exploratory purposes and to develop ideas, we consider a scenario where the processes of the same type (depression or facilitation) are linearly combined and the interaction between depression and facilitation is multiplicative as for the single depression-facilitation processes. We refer to it as the distributive or cross model. In the Appendix D we discuss other possible formulations. The ideas we develop here can be easily extended to more than two STD processes.

#### 3.7.1 The cross (distributive) model

In this formulation, the depression and facilitation variables *x_k_* and *z_k_*, *k* = 1, 2 obey equations of the form (5)-(6) with parameters *τ_dep,k_*, *τ_fac,k_*, *a_d,k_* and *a_f,k_* for *k* = 1, 2. The evolution of these variables generate the sequences *X_k,n_* and *Z_k,n_* (*k* = 1, 2) given by (24)-(25) with the steady-state values 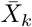 and 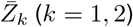 given by (9)-(10). For simplicity, we consider *a_d,_*_1_ = *a_d,_*_2_ and *a_f,_*_1_ = *a_f,_*_2_ (and omit the index *k*) so the differences between two depression or facilitation filters depend only on the differences of the single-event time constants. This can be easily extended to different values of these parameters for the different processes. In what follows, we will not specify the range of the index *k* = 1, 2 unless necessary for clarity.

In the cross (or distributive) model, the variable *M* is given by

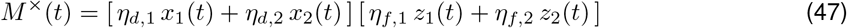

where *η_d,_*_1_ + *η_d,_*_2_ = 1 and *η_f,_*_1_ + *η_f,_*_2_ = 1. Correspondingly, the synaptic update is given by

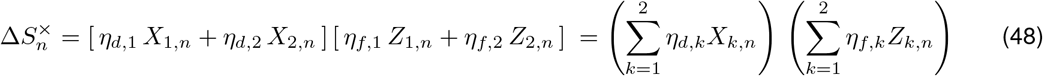

for *n* = 1*,…, N_spk_*. This model allows for all possible interactions between the participating depression and facilitation processes. It reduces to the single depression-facilitation process for *η_d,_*_2_ = *η_f,_*_2_ = 0 (or *η_d,_*_1_ = *η_f,_*_1_ = 0) and allows for independent reductions of depression and facilitation by making *η_d,_*_2_ = 0 or *η_f,_*_2_ = 0, but not both simultaneously.

From (24)-(26),

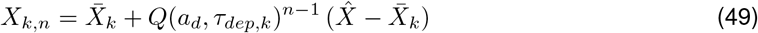

and

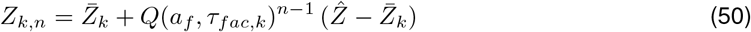

for *k* = 1, 2 with

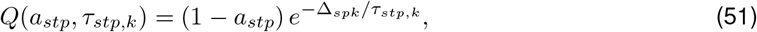

where for use the notation 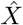 and 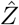 to refer to the first elements in the sequences, which, for simplicity, are assumed to be independent of *k*.

We use the notation

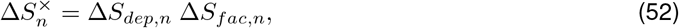

where after algebraic manipulation,

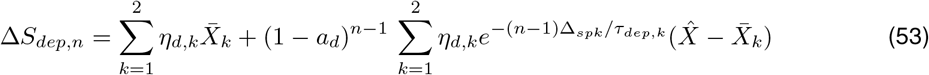

and

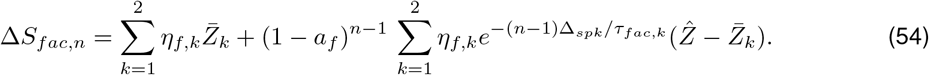

#### 3.7.2 Depression (Δ*S_dep,n_*), facilitation (Δ*S_fac,n_*) and 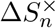 temporal filters

The history-dependent temporal filter Δ*S_dep,n_* transitions from 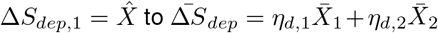 as *n* → ∞, and Δ*S_fac,n_* transitions from 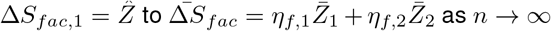 as *n* → ∞. Because the individual filters are monotonic functions, the linearly combined filters represented by the sequences Δ*S_dep,n_* and Δ*S_fac,n_* are also monotonic functions lying in between the corresponding filters for the individual filter components (Figs. 10-A1 and 11-A1 for depression and Figs. 10-A2 and 11-A2 for facilitation). As a consequence, the 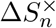 filters also lie in between the product of the corresponding individual filter components Δ*S*_1*,n*_ and Δ*S*_2*,n*_ (Figs. 10-A3 and Figs. 11-A3). In these figures, all parameter values are the same except for *τ_dep,_*_1_ and *τ_fac,_*_1_, which are *τ_dep,_*_1_ = *τ_fac,_*_1_ = 100 in Fig. 10 and *τ_dep,_*_1_ = *τ_fac,_*_1_ = 10 in Fig. 11. In both figures, the values of the facilitation time constants are *τ_dep,_*_2_ = *τ_fac,_*_2_ = 1000.

**Figure 10:**
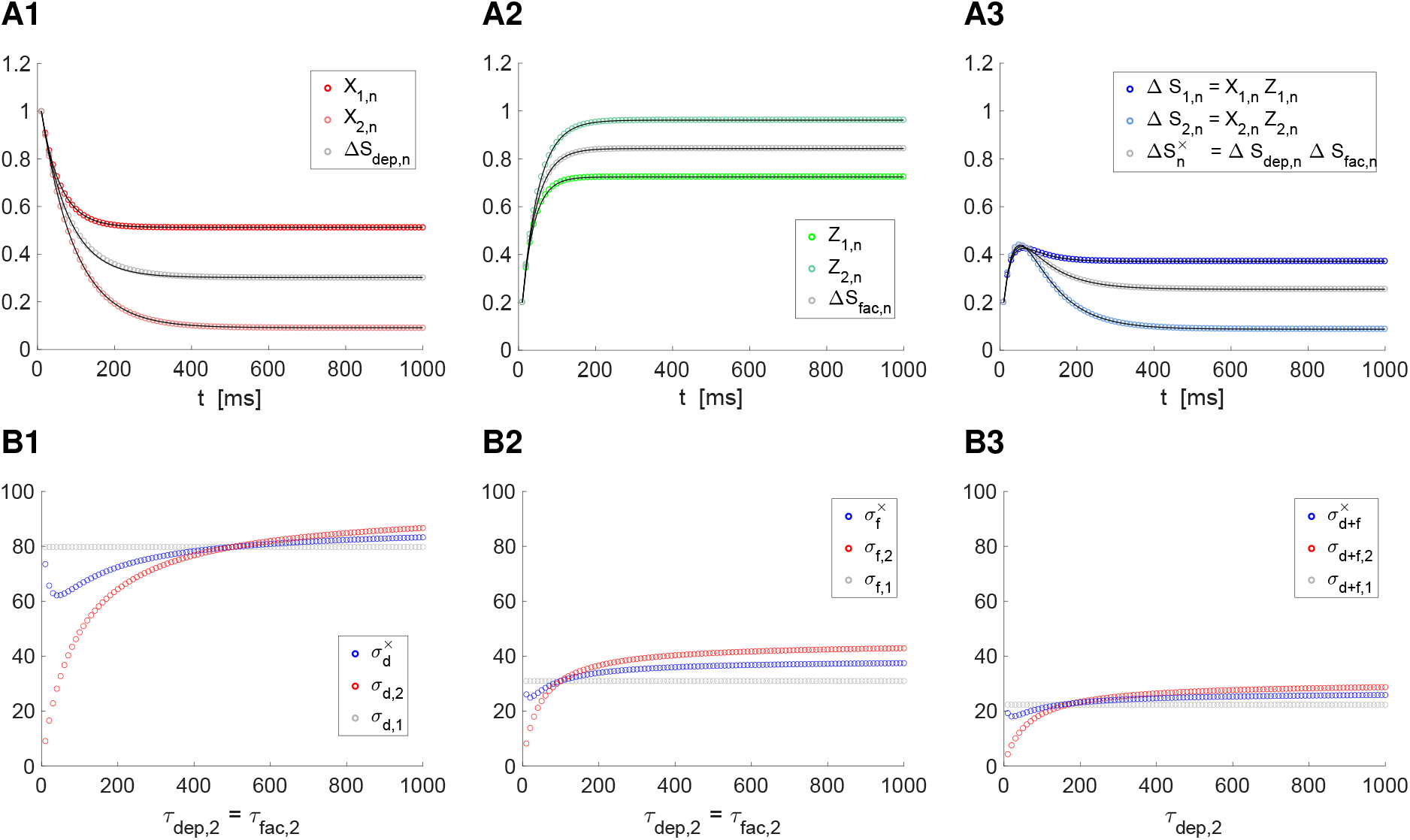
Temporal filters generated by the interplay of multiple depression and facilitation processes with different single-event time scales. **A.** Depression (*X*), facilitation (*Z*) and Δ*S* = *XZ* filters for representative parameter values. We use the distributive model (47)-(52) for the synaptic updates 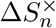. The factors Δ*S_dep,n_* and Δ*S_fac,n_* in 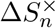 are given by eqs. (53)–(54). The depression and facilitation individual filters *X_k,n_* and *Z_k,n_* (*k* = 1, 2) are given by eqs. (49)–(51). These and the Δ*S_dep,n_* and Δ*S_fac,n_* filters were approximated by using the descriptive envelope model for STD in response to periodic presynaptic inputs (solid curves superimposed to the dotted curves) described in Section 3.3.3 by eqs. (31)–(33). The filter 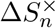 was approximated by using with eq. (39) with *F* and *G* substituted by the corresponding approximations to Δ*S_dep,n_* and Δ*S_fac,n_*. **B.** Dependence of the filter (global) time constants on the single events time constants. We used fixed values of *τ_dep,_*_2_ = *τ_fac,_*_2_ and *τ_dep,_*_1_. Fig. 10 uses a different value of *τ_dep,_*_1_. The filter (global) time constants were computed using eq. (34). We used the following parameter values: *a_d_* = 0.1, *a_f_* = 0.2, *x*_∞_ = 1, *z*_∞_ = 0, *τ_dep,_*_2_ = *τ_fac,_*_2_ = 1000, *η_dep_* = *η_fac_* = 0.5, *τ_dep,_*_1_ = *τ_fac,_*_1_ = 100, and Δ_*spk*_ = 10.

**Figure 11:**
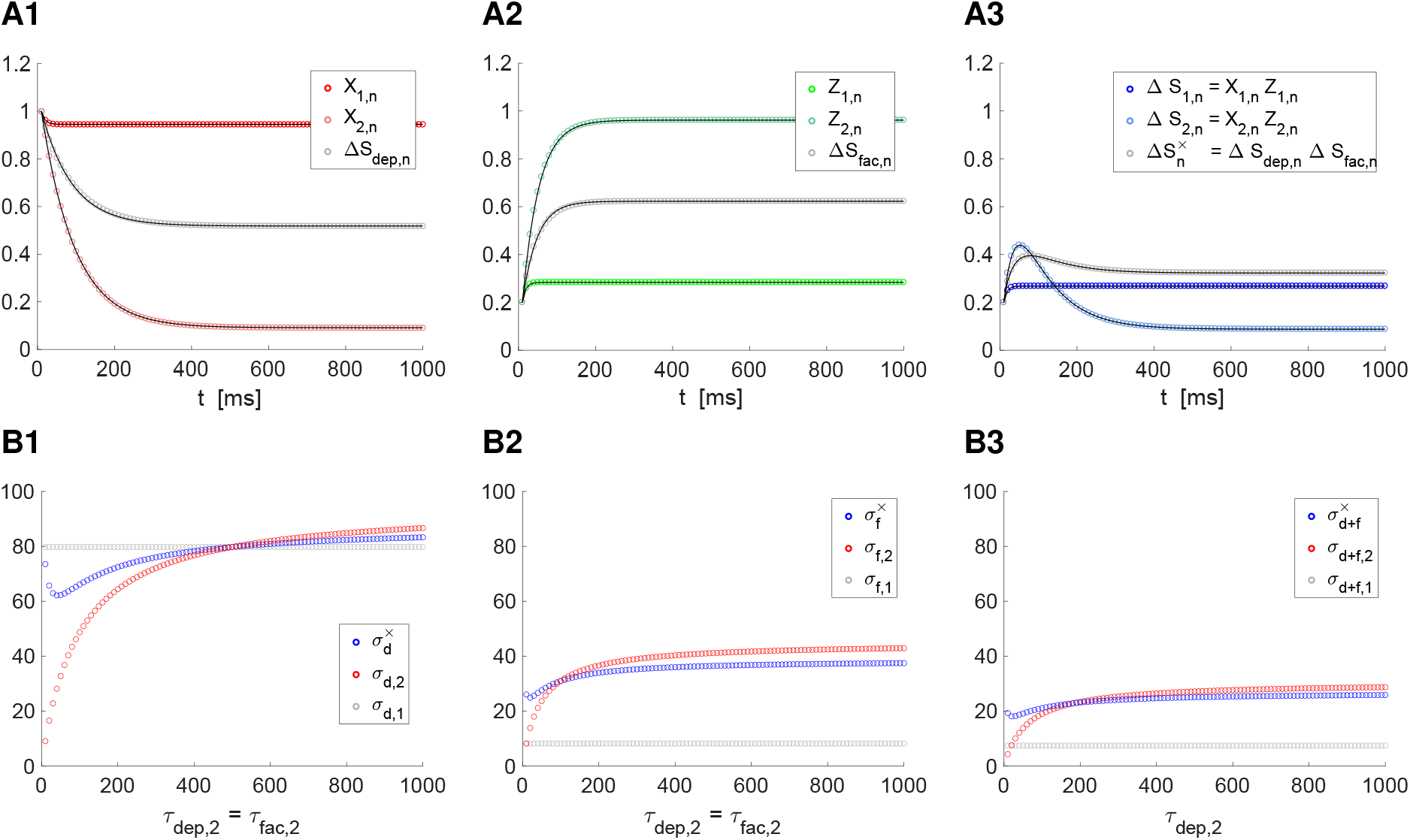
Temporal filters generated by the interplay of multiple depression and facilitation processes with different single-event time scales. **A.** Depression (*X*), facilitation (*Z*) and Δ*S* = *XZ* filters for representative parameter values. We use the distributive model (47)-(52) for the synaptic updates 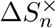. The factors Δ*S_dep,n_* and Δ*S_fac,n_* in 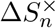 are given by eqs. (53)–(54). The depression and facilitation individual filters *X_k,n_* and *Z_k,n_* (*k* = 1, 2) are given by eqs. (49)–(51). These and the Δ*S_dep,n_* and Δ*S_fac,n_* filters were approximated by using the descriptive envelope model for STD in response to periodic presynaptic inputs (solid curves superimposed to the dotted curves) described in Section 3.3.3 by eqs. (31)–(33). The filter 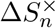 was approximated by using with eq. (39) with with *F* and *G* substituted by the corresponding approximations to Δ*S_dep,n_* and Δ*S_fac,n_*. **B.** Dependence of the filter (global) time constants on the single events time constants. We used fixed values of *τ_dep,_*_2_ = *τ_fac,_*_2_ and *τ_dep,_*_1_. Fig. 10 uses a different value of *τ_dep,_*_1_. The filter (global) time constants were computed using eq. (34). We used the following parameter values: *a_d_* = 0.1, *a_f_* = 0.2, *x*_∞_ = 1, *z*_∞_ = 0, *τ_dep,_*_2_ = *τ_fac,_*_2_ = 1000, *η_dep_* = *η_fac_* = 0.5, *τ_dep,_*_1_ = *τ_fac,_*_1_ = 10, and Δ_*spk*_ = 10.

In Fig. 10-A3 both Δ*S*_1*,n*_ and Δ*S*_2*,n*_ are temporal BPFs peaking almost at the same time. Consequently 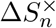 is also temporal BPFs lying strictly in between the individual ones and peaking almost at the same time. In Fig. 11-A3, in contrast, Δ*S*_1*,n*_ is a temporal HPF, while Δ*S*_2*,n*_ is a temporal BPF. The resulting 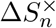 is also a temporal BFP, but the two temporal BPFs peak at different times.

#### 3.7.3 Communication of the single event time scales to the history-dependent filters

In Section 3.3.3 we developed a descriptive envelope model for STD in response to periodic presynaptic inputs consisting of the functions *F* (*t*) for depression, *G*(*t*) for facilitation, and *H*(*t*) = *F* (*t*)*G*(*t*) for the synaptic update, given by eqs. (31)–(33) and (39). By approximating the model parameters using the results of our simulations for *x*(*t*) and *z*(*t*), we computed the filter time constants *σ_d_*, *σ_f_* using (34) and 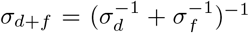. This approach is not strictly necessary for the DA model since the sequences *X_n_* and *Z_n_* can be computed analytically as well as the filter time constants *σ_dep_*, *σ_fac_* and *σ_dep_*_+*fac*_, which convey the same dynamic information as *σ_d_*, *σ_f_* and *σ_d_*_+*f*_. The calculation of these time constants is possible since the filter sequences involved a single *n*-dependent term. However, this is not the case for Δ*S_dep,n_* and Δ*S_fac,n_*, which are linear combinations of *n*-dependent terms. On the other hand, the shapes of Δ*S_dep,n_* and Δ*S_fac,n_* suggest these filters can be captured by the descriptive model by computing the appropriate parameters using the results of our simulations. We use the notation *F_dep_*, *F_fac_* and *H*^×^(*t*) = *F_dep_*(*t*)*F_fac_*(*t*). The solid lines in Figs. 10-A1 to -A3 and 11-A1 to A3 confirm this. In particular, parameter values can be found so that *F_dep_*(*t*), *F_fac_*(*t*) and *H*^×^(*t*) provide a very good approximation to Δ*S_dep,n_*, Δ*S_fac,n_* and 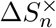, respectively (gray solid lines).

Using the descriptive model we computed the time constants 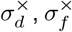 and 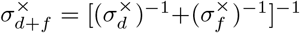. Figs. 10-B and 11-B (blue) show the dependence of these time constants with the single-event depression and facilitation time constant *τ_dep,_*_2_ (= *τ_fac,_*_2_) for two values of *τ_dep,_*_1_ (= *τ_fac,_*_1_) and Δ_*spk*_ = 10 (*f_spk_* = 100). These results capture the generic model behavior via rescalings of the type (28). A salient feature is the non-monotonicity of the curves for 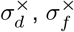 and 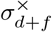 (blue) in contrast to the monotonicity of the curves for *σ_d,_*_2_, *σ_f,_*_2_ and *σ_d_*_+*f,*2_ for the depression and facilitation second component (red).

#### 3.7.4 Degeneracy

The fact that the same type of descriptive envelope models such as the one we use here capture the dynamics of the temporal filters for both single and multiple depression and facilitation processes show it will be difficult to distinguish between them on the basis of data on temporal filters. In other words, the type of temporal filters generated by the DA model (single depression and facilitation processes) are consistent with the presence of multiple depression and facilitation processes interacting as described by the cross (distribute) model.

### 3.8 Persistence and disruption of temporal filters for Δ*S_n_* = *X_n_Z_n_* in response to variable presynaptic spike trains

By design, the temporal filters discussed above emerge in response to periodic presynaptic spike trains (with period Δ_*spk*_). Naturally, a question arises as to whether these type of temporal filters emerge in response to non-periodic inputs and how their properties are affected by input variability. To address this issue, here we consider more realistic, irregular presynaptic spike trains with variable (*n*-dependent) ISIs represented by the sequence 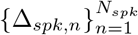. The natural candidates are Poisson spike trains (the ISI distribution follows a Poisson process with the parameter *r* representing the mean firing rate) [69, 80]. For Poisson spike trains both the mean ISI (< *ISI* >) and the standard deviation (SD) are equal to *r*^−1^ and therefore the coefficient of variation CV = 1. For Poisson spike trains with absolute refractoriness *ISI_min_*, < *ISI* >= *r*^−1^ + *ISI_min_* and CV = 1 *ISI_min_ < ISI* >^−1^ [80], making them more regular. We use here *ISI_min_* = 1 so that the irregularity remains high. As a first step, we consider variable presynaptic spike trains consisting of small perturbations to periodic spike trains.

#### 3.8.1 Perturbations to periodic presynaptic spike train inputs

To introduce some ideas, we consider perturbations of periodic presynaptic spiking patterns of the form

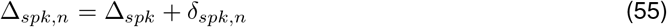

where Δ_*spk*_ is constant (*n*-independent) and 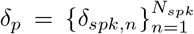 is a sequence of real numbers. The exponential factors in (7)-(8) and (26) read

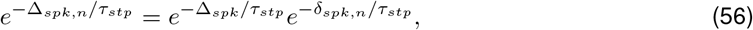

where *τ_stp_* represents *τ_dep_* or *τ_fac_*.

If we further assume |*δ_spk,n_/τ_stp_*| ≪ 1 for all *n*, then

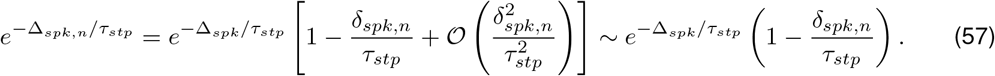

Eqs. (7)–(8) have the general form

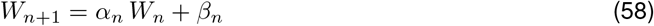

for *n* = 1, 2*,…, N_spk_* − 1 with

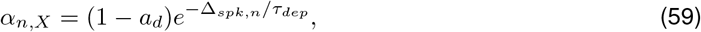

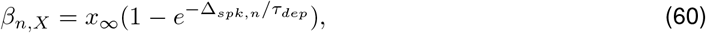

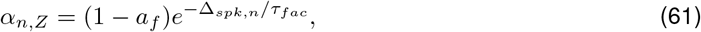

and

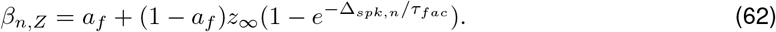

Substitution of (57) into these expressions yields

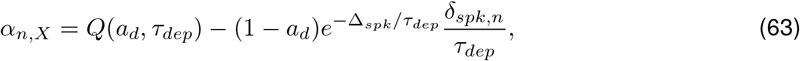

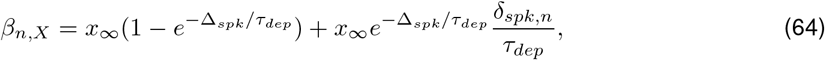

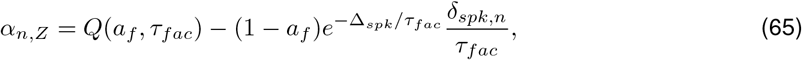

and

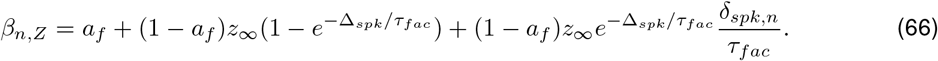

The last terms in these expressions are the 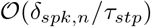 corrections to the corresponding parameters for the constant values of Δ_*spk*_ (*δ_p_* = 0, remaining terms) and contribute to the variance of the corresponding expressions. One important observation is that these variances monotonically increase with decreasing values of Δ_*spk*_ (increasing values of *f_spk_*). A second important observation is the competing effects exerted by the parameters *τ_stp_* (*τ_dep_* and *τ_fac_*) on the variance through the quotients

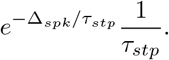

As *τ_stp_* decreases (increases), 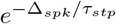 decreases (increases) and 1/*τ_stp_* increases (decreases). In the limit, 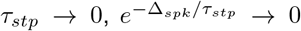 and 1/*τ_stp_* → ∞ and vice versa. Therefore, one expects the variability to change non-monotonically with *τ_dep_* and *τ_fac_*.

To proceed further, we use the notation

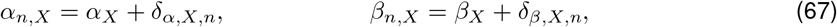

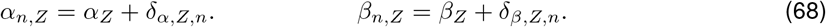

Substituting into (83) in the Appendix A.2 we obtain

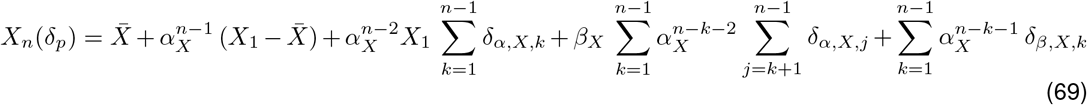

and

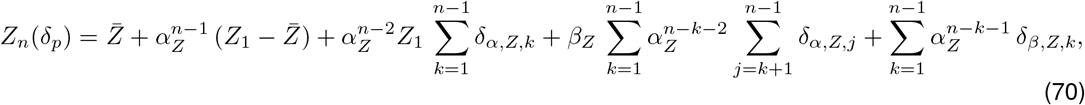

where *α_X_* = *Q*(*a_d_, τ_dep_*) and *α_Z_* = *Q*(*a_f_, τ_fac_*). In (69) and (70) the first two terms correspond to the solution (24) and (25) to the corresponding systems in response to a presynaptic spike train input with a constant ISI Δ_*spk*_ (*δ_p_* = 0), which were analyzed in the previous sections. The remaining terms capture the (first order approximation) effects of the perturbations *δ_p_* = {*δ_spk,n_*}, which depend on the model parameters through *α_X_*, *β_X_*, *α_Z_*, *β_Z_* and the sequences *δ_α,X,n_*, *δ_β,X,n_*, *δ_α,Z,n_* and *δ_β,Z,n_* defined by the equations above.

These effects are accumulated as *n* increases as indicated by the sums. However, as *n* increases, both *Q*(*a_d_, τ_dep_*)^*n*^ and *Q*(*a_f_, τ_fac_*)^*n*^ decrease (they approach zero as *n* → ∞) and therefore the effect of some terms will not be felt for large values of *n* → ∞ provided the corresponding infinite sums converge. On the other hand, the effect of the perturbations will be present as *n* → ∞ in other sums. For example, for *k* = *n* − 1 in the last sums in (69) and (70), 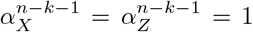 and therefore both *δ_β,X,n_*_−1_ and *δ_β,Z,n_*_−1_ will contribute *X_n_* and *Z_n_*, respectively, for all values of *n*. For small enough values of *δ_p_*, the response sequences *X_n_*(*δ_p_*) and *Z_n_*(*δ_p_*) will remain close *X_n_*(0) and *Z_n_*(0), respectively (the response sequences to the corresponding unperturbed, periodic spike train inputs) and therefore the temporal filters will persist. Fig. 12 shows that this is also true for higher values of *δ_p_*. There, the sequence *δ_p_* was normally distributed with zero mean and variance *D* = 1. In all cases, the mean sequence values computed after the temporal filter decayed (by taking the second half of the sequence points for a total time *T_max_* = 100000, *X_c_* and *Z_c_*) coincides to a good degree of approximation with fixed-point of the unperturbed sequences 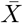 and 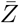 (compared the corresponding solid and dotted curves).

**Figure 12:**
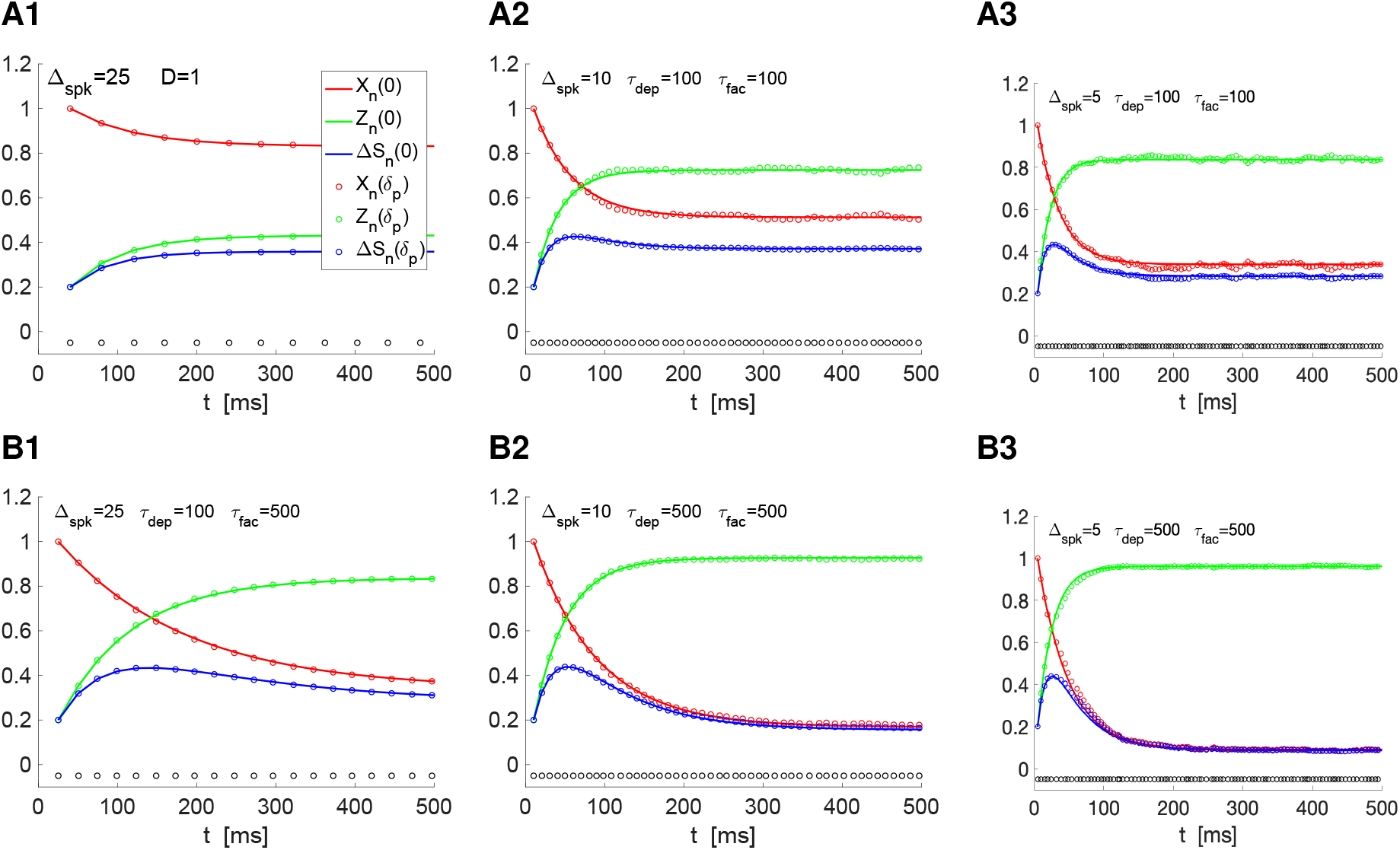
Temporal filters persist in response to variable presynaptic spike trains: Depression, facilitation and synaptic update response to normally distributed ISI perturbations to periodic spike train inputs. We used the recurrent equations (7) and (8) for *X_n_* and *Z_n_* respectively. The ISIs are perturbations around the ISI with constant Δ*spk* according to eq. (55) where the sequence 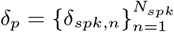 is normally distributed with zero mean and variance *D* = 1. Simulations were run for a total time *T_max_* = 100000. The sequences *X_c_*, *Z_c_* and Δ*S_c_* consist of the last half set of values for *X_p_*, *Z_p_* and Δ*S_p_*, respectively, after the temporal filters decay to a vicinity of 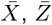 and 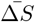. **A.***τ_dep_* = *τ_fac_* = 100. **A1.** var(*X_c_*) = 0.000015, var(*Z_c_*) = 0.000020, var(Δ*S_c_*) = 0.000002. **A2.** var(*X_c_*) = 0.000057, var(*Z_c_*) = 0.000065, var(Δ*S_c_*) = 0.000006. **A3.** var(*X_c_*) = 0.000152, var(*Z_c_*) = 0.000092, var(Δ*S_c_*) = 0.000053. **B.***τ_dep_* = *τ_fac_* = 500. **B1.** var(*X_c_*) = 0.000006, var(*Z_c_*) = 0.000003, var(Δ*S_c_*) = 0.000002. **B2.** var(*X_c_*) = 0.000012, var(*Z_c_*) = 0.000005, var(Δ*S_c_*) = 0.000008. **B3.** var(*X_c_*) = 0.000016, var(*Z_c_*) = 0.000006, var(Δ*S_c_*) = 0.000013. We used the following parameter values: *a_d_* = 0.1, *a_f_* = 0.2, *x*_∞_ = 1, *z*_∞_ = 0.

These results also confirm (by inspection) the previous theoretical observations. First, the variability is smaller for *τ_dep_* = *τ_fac_* = 500 than for *τ_dep_* = *τ_fac_* = 100. Second, as Δ_*spk*_ decreases *f_spk_* increases), the variability increases. For *τ_dep_* = *τ_fac_* = 100 (Fig. 12-A) the variability of Δ*S_c_* = *X_c_Z_c_* is smaller than the variabilities of both *X_c_* and *Z_c_*. For *τ_dep_* = *τ_fac_* = 100 (Fig. 12-B), the variability of Δ*S_c_* = *X_c_Z_c_* is smaller than the variability of *X_c_*, but not always smaller than the variability of *Z_c_*.

While this approach is useful to understand certain aspects of the temporal synaptic update filtering properties in response to non-periodic presynaptic spike train inputs, it is limited since it does not admit arbitrarily large perturbations, which could cause the perturbed ISI to be negative. One solution is to make phase-based perturbations instead of time-based perturbations. But it is not clear whether comparison among patterns corresponding to different Δ_*spk*_ are meaningful.

#### 3.8.2 Poisson distributed presynaptic spike train inputs

Fig. 13 shows the response of the depression (*X_n_*), facilitation (*Z_n_*) and synaptic update (Δ*S_n_*) peak sequences to Poisson distributed presynaptic spike train inputs for representative values of the spiking mean rate *r_spk_* and the depression and facilitation time constants *τ_dep_* and *τ_fac_*. Each protocol consists of 100 trials. A comparison between these responses and the temporal filters in response to periodic presynaptic spike inputs with a frequency equal to *r_spk_* (the Poisson rate) shows that collectively the temporal filtering properties persist with different levels of fidelity. Clearly, variability in the responses are present for each trial (Fig. S3). Figs. S3-B2 and -C2, the temporal band pass-filter is terminated earlier than the corresponding deterministic one. However, in Fig. S3-B1 the temporal band-pass filter is initiated earlier than the corresponding deterministic one.

**Figure 13:**
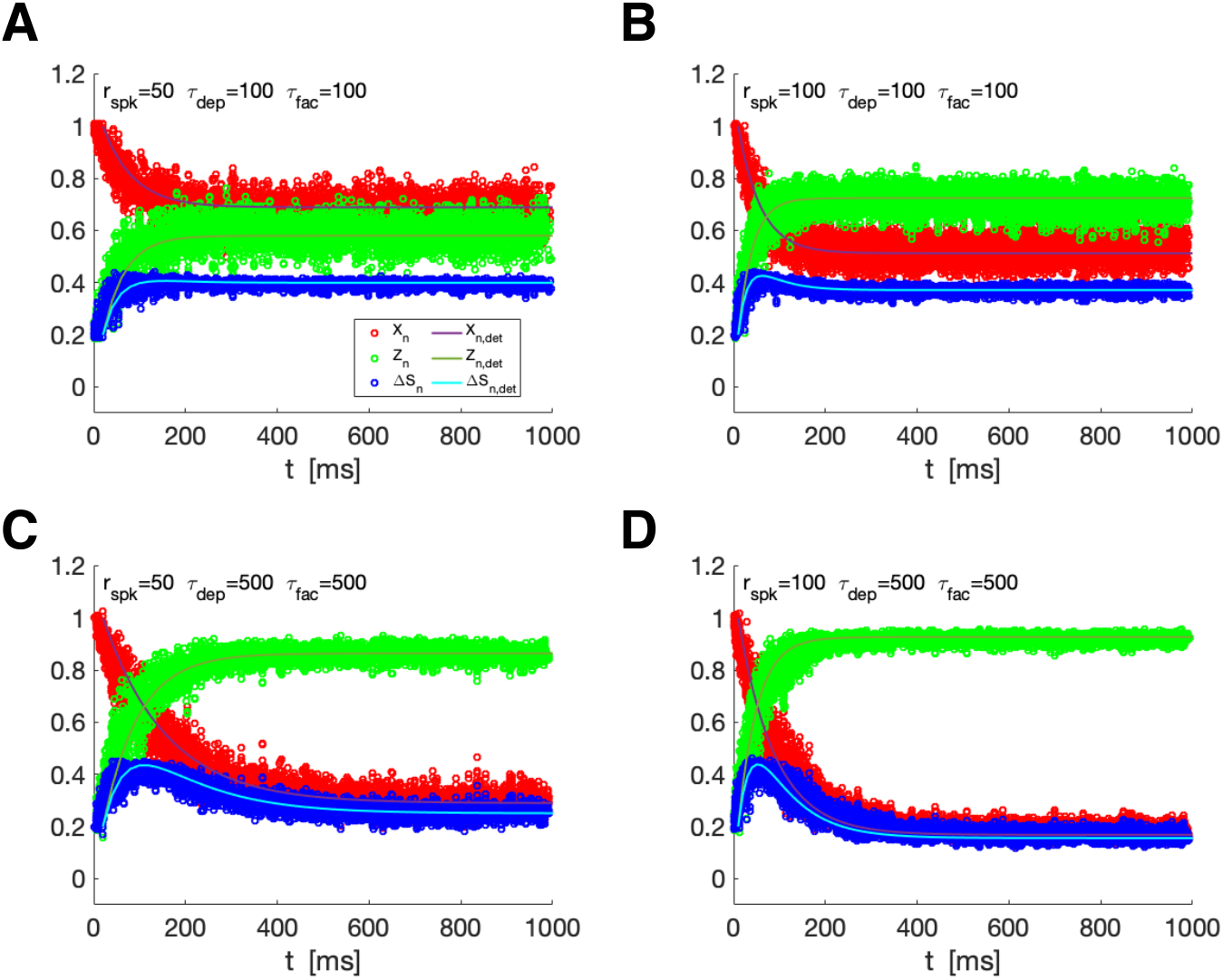
Temporal filters persist in response to variable presynaptic spike trains: Synaptic update response to Poisson distributed spike train inputs. We used the recurrent equations (7) and (8) for *X_n_* and *Z_n_* respectively. The ISIs have mean and standard deviation equal to *r_spk_*. Simulations were run for a total time *T_max_* = 2000 (Δ*t* = 0.01). We consider 100 trials and averaged nearby points. **A.***τ_dep_* = *τ_fac_* = 100 and *f_spk_* = 50. **B.***τ_dep_* = *τ_fac_* = 100 and *f_spk_* = 100. **C.***τ_dep_* = *τ_fac_* = 500 and *f_spk_* = 50. **D.***τ_dep_* = *τ_fac_* = 500 and *f_spk_* = 500. We used the following parameter values: *a_d_* = 0.1, *a_f_* = 0.2, *x*_∞_ = 1, *z*_∞_ = 0.

In Fig. 14 we briefly analyze the response variability of the *X_n_*, *Z_n_* and Δ*S_n_* sequences induced by the presynaptic ISI variability. For *X_n_* and *Z_n_*, the variability decreases with increasing values of *τ_dep_* and *τ_fac_* in an *r_spk_*-dependent manner (Fig. 14-A). For most cases, the variability also decreases with increasing values of *r_spk_* in a *τ_dep_*- and *τ_fac_*-manner (Fig. 14-B). An exception to this rule is shown in Fig. 14-B1 for the lower values of *r_spk_*.

**Figure 14:**
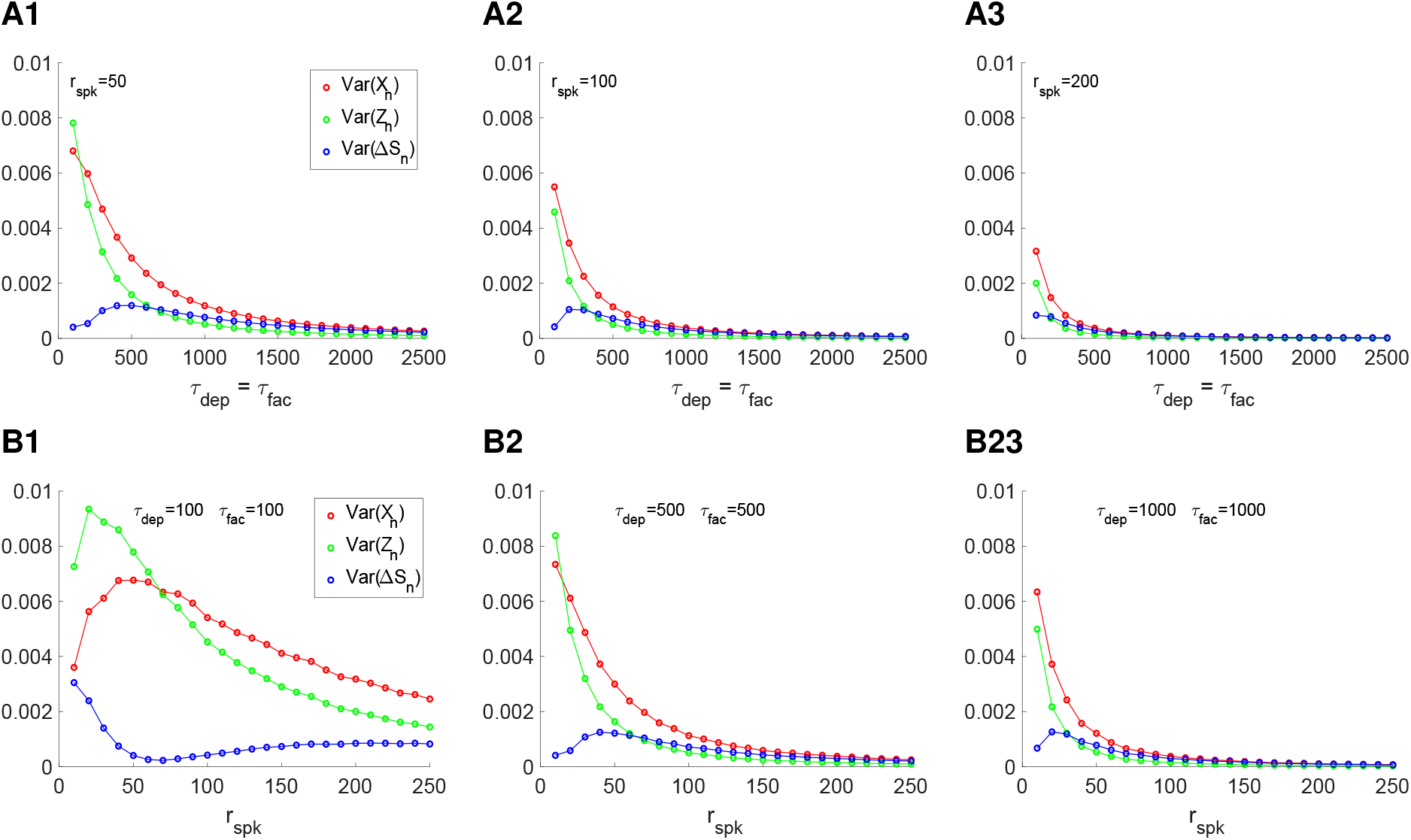
Response variability to non-periodic presynaptic spike trains: depression, facilitation and synaptic update response to Poisson spike train inputs. We used the recurrent equations (7) and (8) for *X_n_* and *Z_n_* respectively. The ISIs have mean and standard deviation equal to *r_spk_*. Simulations were run for a total time *T_max_* = 500000 (Δ*t* = 0.01). **A.** Variance as a function of *τ_dep_* and *τ_fac_* (*τ_dep_* = *τ_fac_*) for fixed values of the Poisson input spike rate. **A1.***r_spk_* = 50. **A2.***r_spk_* = 100. **A3.***r_spk_* = 200. **B.** Variance as a function of the Poisson spike rate for fixed values of *τ_dep_* and *τ_fac_* (*τ_dep_* = *τ_fac_*). **B1.***τ_dep_* = *τ_fac_* = 100. **B2.***τ_dep_* = *τ_fac_* = 500. **B3.***τ_dep_* = *τ_fac_* = 1000. We used the following parameter values: *a_d_* = 0.1, *a_f_* = 0.2, *x*_∞_ = 1, *z*_∞_ = 0.

The variability of the Δ*S_n_* sequences is more complex. Figs. 14-A1 and -A2 show examples of the peaking at intermediate values of *τ_dep_* and *τ_fac_*. Fig. 14-B1 shows an example of the variability reaching a trough for an intermediate value of *r_spk_* and relatively low values of *τ_dep_* and *τ_fac_*, while Figs. 14-B2 and -B3 show examples of the variability peaking at intermediate values of *r_spk_* for higher values of *τ_dep_* and *τ_fac_*. This different dependence of the variability with the model parameters and input rates emerge as the result of the different variability properties of the product of the sequences *X_n_* and *Z_n_*. A more detailed understanding of these properties is beyond the scope of this paper.

### 3.9 The MT model exhibits similar filtering properties as the DA model: Dynamics of the depression (*R*) and facilitation (*u*) variables and their interaction

The main difference between the DA model (5)-(6) and the MT model (12)-(13) is the update of the depression variables (*x* in the DA model and *R* in the MT model). Notation aside, the dynamics for depression and facilitation variables *x* and *z* in the DA model are completely independent both during the presynaptic ISI and the update. In the MT model, in contrast, while the dynamics of the depression and facilitation variables *R* and *u* are independent during the presynaptic ISI as well as the *u*-update, the *R*-update is dependent on *u*^+^. As a result, the difference equations describing the peak sequence dynamics for *R_n_* and *u_n_* (14)-(15) (peak envelope responses to periodic presynaptic inputs) are not fully independent, but the equation for *R_n_* is forced by the sequence *u_n_*, which is independent of *R_n_*. For the DA model, the difference equations for *Z_n_* and *Z_n_* are independent (7)-(8). Naturally, the steady-states (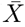 and 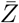) in the DA model are independent, while in the MT model, the steady state 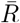 depends on the steady state 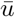. Here we show that despite these differences and the increased difficulty in the interpretation of the analytical solution for the MT model as compared to the DA model for the determination of the long-term depression and facilitation filter time constants, the two models describe similar dynamics.

Standard methods (see Appendix A) applied to these linear difference equations for Δ_*spk,n*_ = Δ_*spk*_ (independent of *n*) yield

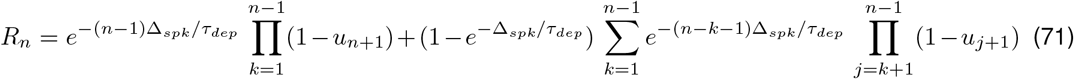

and

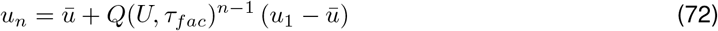

where *Q*(*U, τ_fac_*) is given by (26).

Because of the complexity of (71) we are not able to use the approach described in Section 3.3.1 to compute the (long-term) history-dependent time scales *σ_dep_* and *σ_fac_* in terms of the single event time scales (*τ_dep_* and *τ_fac_*) given by eq. (27). Instead, we use the descriptive modeling approach described in Section (3.3.3) by eqs. (31)–(34) and Section 3.3.3.

## 4 Discussion

The temporal and frequency-dependent properties of postsynaptic patterns are shaped by the presence of synaptic short-term plasticity (synaptic depression and facilitation; STP). In response to a presynaptic spike train, the postsynaptic membrane potential response may be amplified, attenuated or both, thus exhibiting a maximal response for an intermediate presynaptic spike (or spike sequence). This gives rise to the notion of STP-mediated temporal filtering: the response is optimal within a certain time window (or windows). During these temporal bands, sensory input is enhanced and the communication between neurons is facilitated.

We set out to understand the mechanisms of generation of temporal filters in response to presynaptic spike trains in the presence of STP. We focused on a feedforward network consisting of a presynaptic cell (modeled as a presynaptic spike train) synaptically connected to a passive cell (diagram in Fig. 2). This is the minimal network model that allows the systematic investigation of postsynaptic potential (PSP) temporal filters in response to presynaptic inputs. In our simulations we primarily used parameters consistent with AMPA excitation. We adopted the use of periodic spike trains as the reference presynaptic spiking input. This allowed us to conduct a systematic study of temporal filters. First, we characterized the three types of temporal filters that emerge: low-, high-, and band-pass filters (LPF, HPF, BPF, respectively). Second, we systematically investigated how their properties depend on the properties of the network building blocks, particularly the time constants involved in the sequence of concatenated processes: (i) the presynaptic spike train ISI Δ_*spk*_, (ii) the short-term depression and facilitation *τ_dep_* and *τ_fac_*, (iii) the synaptic decay time *τ_dec_*, and (iv) the membrane time constant *τ_m_*. We then showed that the reference temporal filters are preserved at the population (multiple trial) level in response to variable presynaptic spike trains. The degree of variability of these patterns within and across trials depends on the parameter values, but the temporal filtering properties remain. To our knowledge, this is the first systematic investigation of STP-mediated neuronal temporal filters. Our results have implications for the understanding of the mechanism underlying the temporal information filtering properties of neuronal systems discussed in the Introduction.

We used two biophysically plausible phenomenological models that have been widely used in the literature: the DA (Dayan-Abbott) and the MT (Markram-Tsodkys) models [21, 43, 62, 63, 67–73]. In the DA model [69], the depression and facilitation variables evolve independently, while in the MT model [63], the evolution of the depression variable is affected by the facilitation variable. We found no significant differences between the results for both models. The simplicity of the DA model allows for a number of analytical calculations that facilitate the analysis and the mechanistic understanding. From the differential equations describing the continuous evolution of the depression (*x*) and facilitation (*z*) processes one can extract the difference equations describing the discrete evolution of the peak sequences *X_n_* and *Z_n_*, respectively. These can be solved analytically providing the input Δ*S_n_* = *X_n_Z_n_* to the synaptic variable *S* at the arrival of each presynaptic spike. The solution to the difference equation for the synaptic peak sequences *S_n_* produces expressions for the synaptic peaks. The investigation of the MT model required the development of additional tools and numerical simulations since the difference equations for the depression variable (*R*) is nonlinear and not analytically solvable. Of particular importance is the development of a descriptive modeling approach to capture the shape of the temporal filters in terms of the model parameters or data (see Supplementary Material Section for the more detailed analysis). In contrast to the DA model where the temporal filter parameters (e.g., the filter time constants *σ_dep_* and *σ_fac_*) are derived from the single event parameters, for the MT model the temporal filter parameters (e.g., the filter time constants *σ_dep_* and *σ_fac_*) are inferred from the shapes obtained by simulating the equations for the depression and facilitations variables (*R* and *u*). An additional step is needed to relate the filter parameters to the single event parameters. This approach can be easily adapted to more complex models for which analytical solutions are not available and to experimental data following a similar protocol.

Dynamically, temporal BPFs can be considered as overshoot types of solution to a linear difference equation. Overshoots are not possible for one-dimensional linear difference equations (e.g., temporal LPFs and HPFs), but they are possible for two-dimensional linear difference equations. This implies that two time scales would be enough to explain the properties of BPFs for Δ*S_n_*. However, our results indicate that a third time scale is needed to explain the BPF properties for Δ*S_n_* in the general case, consistent with previous results [75]. This emergent time scale combines the first two and is further propagated to the higher levels of organization.

The interaction between the STP-mediated LPFs and HPFs with the synaptic and postsynaptic dynamics generates additional LPFs, HPFs and BPFs, with additional emergent time scales. For relatively low membrane time constants, the postsynaptic dynamics reflect the synaptic dynamics and the PSP filters are proportional to the synaptic filters (e.g., [21, 63, 67]). However, for higher membrane time constants, the PSP filters depart from this proportionality with the synaptic ones. Specifically, PSP BPF emerge in the presence of synaptic LPFs or in the presence of synaptic BPFs, but having different shapes and peaking at different times. This additional processing affects the communication between pre- and postsynaptic cells in the presence of STP.

In order to account for more realistic situations, we considered scenarios where more than one depression and facilitation processes with different time constants interact. The results are consistent with the ones for the single processes. However, the models we used (in the main body and in the Appendix) have been developed as natural extensions of the ones for single processes and are not based on observations or previous information about the presence of multiple depression and facilitation processes. More research is needed to determine whether these models and the resulting filters capture realistic situations.

An important question we addressed in our work is how the single event time constants (e.g., *τ_dep_*, *τ_fac_*, *τ_dec_*), which control the systems’ dynamics during the ISIs, are communicated to the temporal filters. In other words, how the temporal filters’ long-term time constants (*σ_dep_*, *σ_fac_*, *σ_sum_* for the DA model and *σ_d_*, *σ_f_*, *σ_sum_* for the MT model) depend on the single-event time constants for each presynaptic spike train ISI Δ_*spk*_. For the simplest synaptic model (one-dimensional linear dynamics for the variable synaptic variable *S* during the ISI and a constant update Δ*S*, independent of *S*), the single-event and temporal filter time constant coincide. For the slightly more complex models for depression and facilitation (one-dimensional linear dynamics for the variables *x* and *z* during the ISI, but the updates depend on the appropriate values of the variables at the arrival of the presynaptic spikes), there is a departure of the temporal filter time constants from the single event time constants. The dependence between the two types of time constants (filter and single event) is relatively complex and involves the presynaptic time scale Δ_*spk*_ and additional parameter values. This complexity is propagated to the PSP filters and is expected to be further propagated to higher levels of organization that are beyond the scope of the paper, but not unimportant.

While biophysically plausible, the phenomenological models of STP we used in this paper are relatively simple and leave out a number of important biological details that might contribute to determining the properties of STP-mediated temporal filters and their consequences for information processing. Further research is needed to understand the properties of these filters and how they emerge as the result of the interaction of the building blocks. An additional aspect that requires attention is the possible effect of astrocyte regulation of STP [10, 11] on the mechanisms of generation of STP-dependent temporal filters. Our work leaves out the mechanisms of generation and properties of the stationary low-, high- and band-pass filters and the associated phenomenon of synaptic and postsynaptic resonance. This will be discussed elsewhere.

The conceptual framework we developed in this paper allows the development of ideas on the properties of PSP temporal filters in response to presynaptic inputs in the presence of STP and the mechanism underlying their generation. An important aspect of this framework is the separation of the feedforward network into a number of building blocks, each one with its own dynamics. The emerging temporal filters can be analyzed in terms of the hierarchical interaction of these building blocks. This conceptual framework can be used to investigate the properties of low-, high- and band-pass stationary, frequency-dependent filters and the emergence of synaptic and postsynaptic resonances. It is conceived to be further extended to include a number of more complex scenarios, including non-periodic synaptic spike trains (e.g., Poisson spike inputs, bursting patterns with two or more spiking frequencies), more complex networks (e.g., two recurrently connected cells with STP in both synapses, three-cell feedforward networks with STP in both synapses), the modulatory effects of astrocytes, more complex postsynaptic dynamics involving ionic currents that have been shown to produce resonances [81–87], and the generation of postsynaptic spiking temporal filters. A first step in this direction is to extend the notion of STP-mediated temporal filters to the postsynaptic spiking domain and characterize the resulting firing rate temporal filters.

Our results make a number of predictions that can be experimentally tested both *in vitro* and *in vivo* using current clamp, and optogenetics [88, 89]. These primarily pertain to the dependence of the type and shape of the temporal PSP filters with the presynaptic spikes and the STP properties. These include our results in Figs. 8 and 9 (and analogous figures for the MT model) and extensions to additional results about the dependence of these filters with the model parameters (not presented here for lack of space) that can be obtained by using our modeling approach. In particular, we predict that PSP filters in the presence of STP are not proportional to the product of the synaptic depression and facilitation variables, but reflect the processing occurring at the postsynaptic level. The fact that temporal PSP filters persist in response to variable presynaptic spike inputs is important for this task. Our results using the simplified models also generate hypothesis to be tested in more detailed models of STP. From a different perspective, the PSP temporal filters can be used to infer the model parameters describing the single event processes and to extract biophysical and dynamic information from experimental data.

## Acknowledgments

This work was partially supported by the National Science Foundation grant DMS-1608077 (HGR) and an NSF Graduate Research Fellowship (YM). The authors are grateful to Allen Tannenbaum for useful comments and support, and to Farzan Nadim, Dirk Bucher and Nelly Daur for useful discussions.

## A 1D linear difference equations

## A.1 Constant coefficients

Consider the following linear difference equation

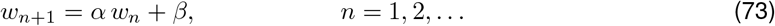

where *α* and *β* are constants. The steady-state for this equation, if it exists, is given by

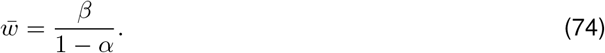

By solving (73) recurrently and using

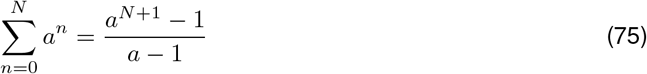

where *a* ≠ 1 is a real number, one gets

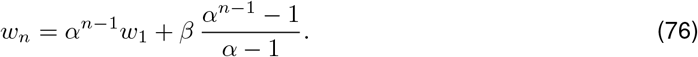

Substitution of (74) into this equation yields

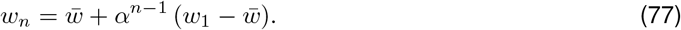

Application of formula (77) to the difference equations (7) and (8) gives, respectively,

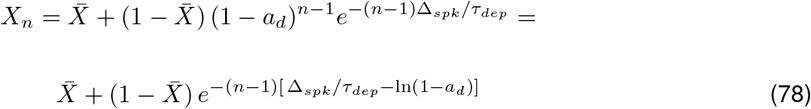

and

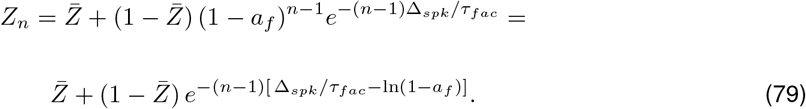

## A.2 Variable (*n*-dependent) coefficients

Consider the following linear difference equation

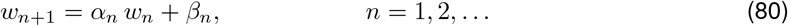

By solving (73) recurrently one gets

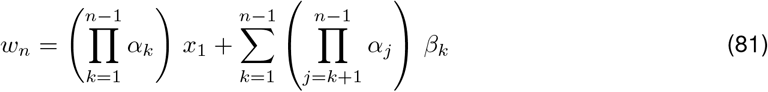

where we are using the convention 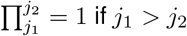. Eq. (81) reduces to eq. (79) if both coefficients in (81) are constant.

Consider now eq. (80) where the coefficients are expressed as small perturbations *δ_α,n_* ≪ 1 and *δ_β,n_* ≪ 1 (*n* = 1, 2*,…*), respectively, of constant coefficients

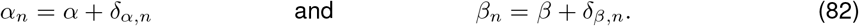

To the first order approximation, the solution (81) reads

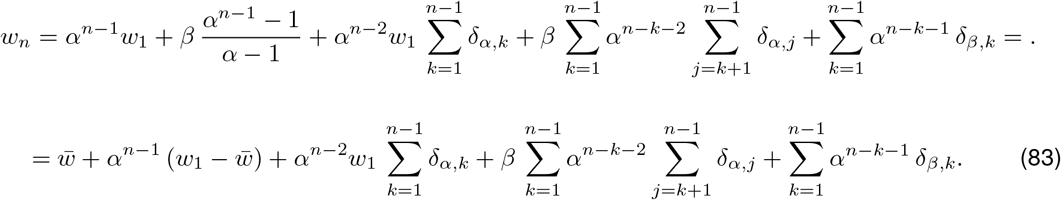

## B Some properties of 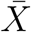 and 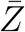 and their dependence with Δ_*spk*_ and *τ_dep/fac_*

Consider 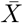 and 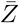 given by (9) and (10), respectively.

## B.1 Monotonic dependence of 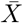 and 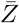 with Δ_*spk*_

If *a_d_* > 0 and *x*_∞_ > 0, then 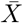 is an increasing function of Δ_*spk*_ and a decreasing function of *f_spk_*. This results from

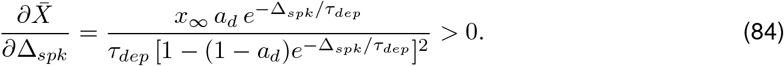

If *a_f_* < 1 and *z*_∞_ < 1, then 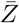 is a decreasing function of Δ_*spk*_ and an increasing function of *f_spk_*. This results from

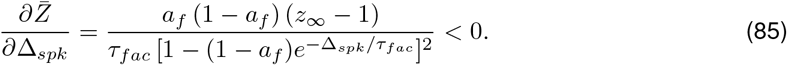

## B.2 Monotonic dependence of 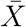 and 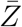 with *τ_dep/fac_*

If *a_d_* > 0 and *x*_∞_ > 0, then 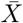 is an decreasing function of *τ_dep_*. This results from

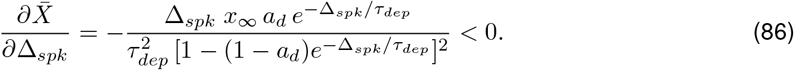

If *a_f_* < 1 and *z*_∞_ < 1, then 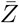 is a decreasing function of *τ_fac_*. This results from

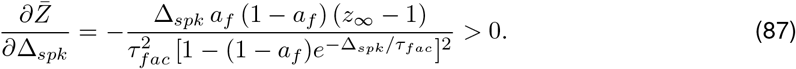

## C Models of synaptic depression and facilitation

## C.1 Depression - facilitation model used in [90]

Following [91, 92], the synaptic variables *S* obey a kinetic equation of the form

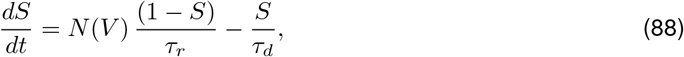

where *N* (*V*) (mM) representes the neurotransmitter concentration in the synaptic cleft. Neurotransmitters are assumed to be released quickly upon the arrival of a presynaptic spike and remain in the synaptic cleft for the duration of the spike (~ 1 ms). This can be modeled by either using a sigmoid function

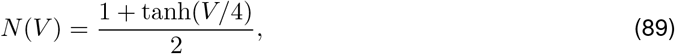

or a step function if the release is assumed to be instantaneous. The parameters *τ_r_* and *τ_d_* are the rise and decay time constants respectively (msec).

This model assumes *N* (*V*) is independent of the spiking history (the value of *N* (*V*) during a spike is constant, except possibly for the dependence on *V*). (There is evidence that this is not realistic [78, 93].) In [90], the “activated” time was 1 ms [94, 95].

In [90], they followed the description of the synaptic short-term dynamics following [43, 62] (Section 2.1.5). For the dynamics of the synaptic function *S*, they used a function [ *T*] = *κ* Δ*S_n_* during the release time and [ *T*] = 0 otherwise, instead of *N* (*V*). The combination of the two formulations yields

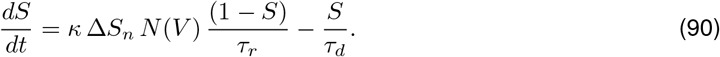

In the following alternative formulation [68] *κ* Δ*S_n_* does not affect the effective rise time of the synaptic function *S*

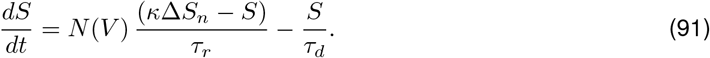

## C.2 Depression model used in [96]

Following experimental procedures described in [97], the synaptic current is described by *I_syn_* = *G_ex_a d*(*V E_ex_*) where *a* and *d* are variables that represent activation and depression processes, respectively. They follow the form:

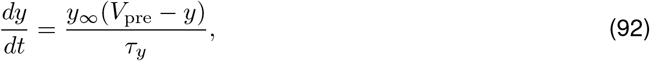

where *y* = *a, d*. The steady-state of *y* is given by

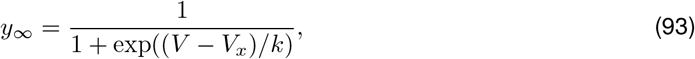

and its time constant follows

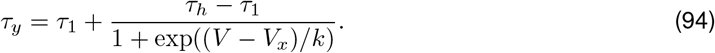

This model is used in [96] to describe bistability in pacemaker networks with recurrent inhibition and depressing synapses. Parameters in these equations are experimentally fitted from the pyloric network of the crab *Cancer borealis*.

## D Additional model formulations for multiple depression-facilitation processes

In Section 3.7 we discussed the model formulation (47)-(48) describing the interplay of two depression-facilitations processes. A number of additional, simplified formulations are possible based on different assumptions. The models we propose here are natural mathematical extensions of the single depression/facilitation processes discussed in the main body of this paper. They are phenomenological models, not based on any experimental observation or theoretical foundation, and they are limited in their general applicability. However, they are useful to explore the possible scenarios underlying the interplay of multiple depression and facilitation time scales affecting the PSP dynamics of a cell in response to presynaptic input trains.

## D.1 Additive and multiplicative segregated-processes models

In the additive and multiplicative *segregated* models, the variable *M* is given, respectively, by

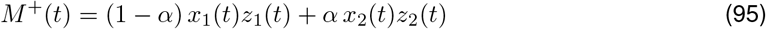

and

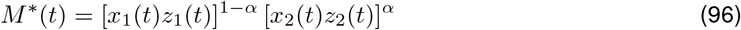

where the parameter *α* ∈ [0, 1] controls the relative contribution of each of the processes. Correspondingly, the updates are given by

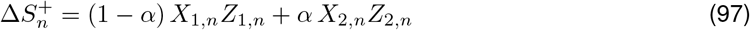

and

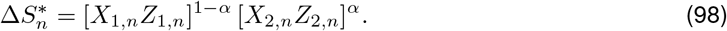

For *α* = 0, 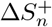 and 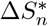 reduce to Δ*S*_1*,n*_ (single depression-facilitation process). This accounts for the regimes where *τ_dep,_*_2_, *τ_fac,_*_2_ ≪ 1. If the two processes are equal (*τ_dep,_*_1_ = *τ_dep,_*_2_ and *τ_fac,_*_1_ = *τ_fac,_*_2_), then 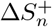 and 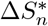 also reduce to Δ*S*_1*,n*_. However, these models fail to account for the reducibility in the situations where only *τ_dep,_*_2_ ≪ 1 or *τ_fac,_*_2_ ≪ 1, but not both. The option of considering depression to be described by *x*_1_ and facilitation by *z*_2_ (with *τ_fac,_*_1_, *τ_dep,_*_2_ ≪ 1) is technically possible in the context of the model, but it wouldn’t be consistent with the model description of single depression-facilitation processes, and it will make no sense to use the model in this way. In general, this model would be useful when the depression and facilitation time scales for each process 1 and 2 are comparable and the differences in these time scales across depression/facilitation processes should be large enough.

## D.2 Fully multiplicative model

One natural way to extend the variable *M* to more than one process is by considering

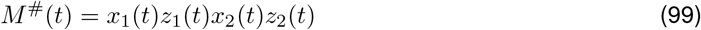

and the synaptic update, given by

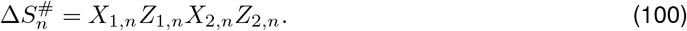

This formulation presents us with a number of consistency problems related to the reducibility (or lack of thereoff) to a single depression-facilitation process in some limiting cases when, for example, the two depression or facilitation time constants are very similar and therefore the associated processes are almost identical, or the depression or facilitation time constants are very small and therefore the envelopes of the associated processes are almost constant across cycles.

More specifically, first, if *τ_dep,_*_2_, *τ_fac,_*_2_ ≪ 1 (almost no STD), then 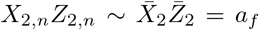 for all *n* after a very short transient and therefore 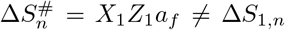. One way, perhaps the simplest, to address this is to divide the expressions (99) and (100) by 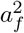 and redefine Δ*S_k,n_* for the single depression-facilitation process accordingly. Specifically,

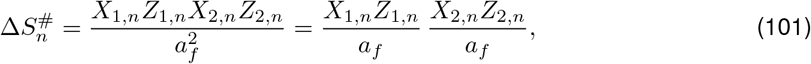

where we use the notation

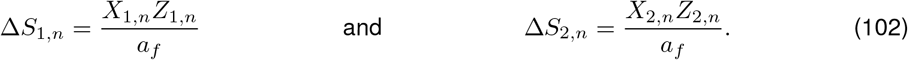

The effect of redefining Δ*S_k,n_* by dividing the original expression (used in the previous sections) does not affect the time constants and the differences in the values between the two formulations is absorbed by the maximal synaptic conductance.

Second, if *τ*_*dep*,1_ = *τ*_*dep*,2_ and *τ*_*fac*,1_ = *τ*_*fac*,2_, then *X*_1*,n*_ = *X*_2*,n*_ and *Z*_1*,n*_ = *Z*_2*,n*_ for all *n*, and 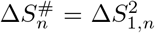 instead of 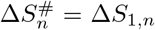. In order to address this, the synaptic update can be modified to

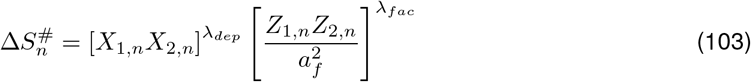

where

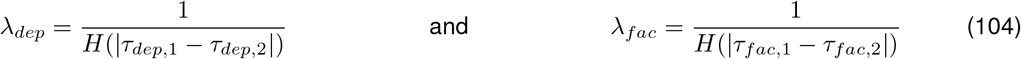

and *H*(Δ*τ*) is a rapidly decreasing function satisfying *H*(0) = 2 and 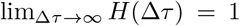. In our simulations we will use

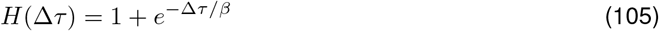

with *β* > 0. Correspondingly,

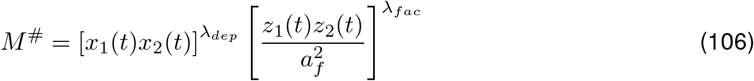

In this way,

- If *τ_dep,_*_1_ = *τ_dep,_*_2_, then *X*_1*,n*_ = *X*_2*,n*_ for all *n* and *λ_dep_* = 1/2. This gives

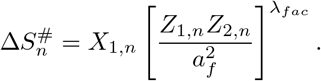 If, in addition, *τ_fac,_*_1_ ≠ *τ_fac,_*_2_ and |*τ_fac,_*_1_ − *τ_fac,_*_2_| > 0 is large enough, then *λ_fac_* = 1 and

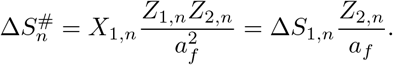
- If *τ*_*fac*,1_ = *τ*_*fac*,2_, then *Z*_1*,n*_ = *Z*_2*,n*_ for all *n*, *λ*_*fac*_ = 2 and

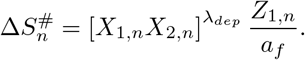 If, in addition, *τdep,*1 ≠ *τ_dep,_*_2_ and |*τ_dep,_*_1_ − *τ_dep,_*_2_| > 0 is large enough, then *λ_dep_* = 1 and

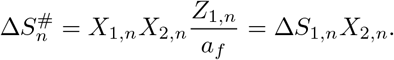
- It follows that if both *τ_dep,_*_1_ = *τ_dep,_*_2_ and *τ_fac,_*_1_ = *τ_fac,_*_2_, then *X*_1*,n*_ = *X*_2*,n*_ and *Z*_1*,n*_ = *Z*_2*,n*_ for all *n*, *λ_dep_* = *λ_fac_* = 2 and

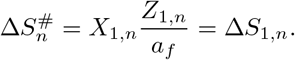
- If *τ_dep,_*_2_ ≪ 1 and *τ_dep,_*_1_ − *τ_dep,_*_2_ is large enough, then *X*_2*,n*_ = 1 for all *n* (after a very short transient), *λ_dep_* = 1, and then

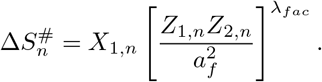 If, in addition, *τ_dep,_*_1_ ≪ 1 and *τ_dep,_*_2_ ~ *τ_dep,_*_1_ (|*τ_dep,_*_1_ − *τ_dep,_*_2_| ~ 0 not large enough), then *X*_1*,n*_ = 1 for all *n* (after a very short transient), *λ_dep_* = 2, and then

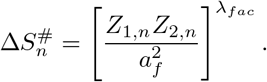
- If *τ_fac,_*_2_ ≪ 1 and *τ_fac,_*_1_ − *τ_fac,_*_2_ is large enough, then *Z*_2*,n*_ = *a_f_* for all *n* (after a very short transient), *λ_fac_* = 1, and then

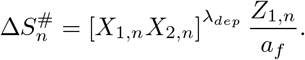 If, in addition, *τ_fac,_*_1_ ≪ 1 and *τ_fac,_*_2_ *τ_face,_*_1_ (*τ_fac,_*_1_ *τ_fac,_*_2_ 0 not large enough), then *Z*_1*,n*_ = *a_f_* for all *n* (after a very short transient), *λ_fac_* = 2, and then

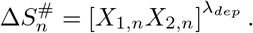
- It follows that if *τ_dep,_*_1_, *τ_dep,_*_2_ ≪ 1 (|*τdep,*1 − *τ_dep,_*_2_| ~ 0 not large enough) and *τ_fac,_*_1_, *τ_fac,_*_2_ ≪ 1 (|*τ_fac,_*_1_ − *τ_fac,_*_2_| ~ 0 not large enough), then

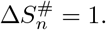

## E Descriptive rules for the generation of temporal (envelope) band-pass filters from the interplay of the temporal (envelope) low- and high-pass filters

From a geometric perspective, temporal band-pass filters in response to periodic presynaptic inputs arise as the result of the product of two exponentially increasing and decreasing functions both decaying towards their steady-state (e.g., Fig. 6). At the descriptive level, this is captured by the temporal envelope functions (*F*, *G* and *H* = *FG*) discussed above whose parameters are not the result of a sequence of single events but are related to the biophysical model parameters by comparison with the developed temporal filters. These functions provide a geometric/dynamic way to interpret the generation of temporal filters in terms of the properties of depression (decreasing functions) and facilitation (increasing functions) in response to periodic inputs, although this interpretation uses the developed temporal filters and therefore is devoid from any biophysical mechanistic interpretation.

In order to investigate how the multiplicative interaction between *F* (*t*) and *G*(*t*) given by eqs. (31)–(32) give rise to the temporal band-pass filters *H* = *FG*, we consider a rescaled version of these functions

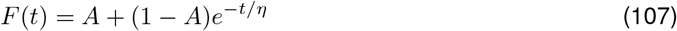

and

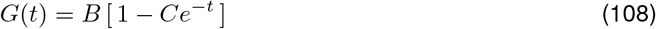

where *B* = 1 and

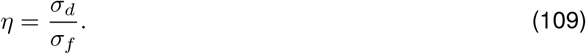

The function *G* transitions from *G*(0) = 1 − *C* to lim_*t*→∞_ *G*(*t*) = 1 with a fixed time constant (Fig. 15, green curves). The function *F* transitions from *F* (0) = 1 to lim_*t*→∞_ *F* (*t*) = *A* with a time constant *η* (Fig. 15, red curves). It follows that *H* transitions from *H*(0) = 1 − *C* to lim_*t*→∞_ *H*(*t*) = *A B* = *A* (Fig. 15, blue curves). A temporal band-pass filter is generated if *H* raises above *A* for a range of values of *t*. This requires *F* to decay slow enough so within that range *H* = *FG > A* (Fig. 15-A) or *A* to be small enough (Fig. 15-B). In fact, as *A* decreases, the values of *η* required to produce a band-pass temporal filter increases (compare Fig. 15-A2 and -B2).

**Figure 15:**
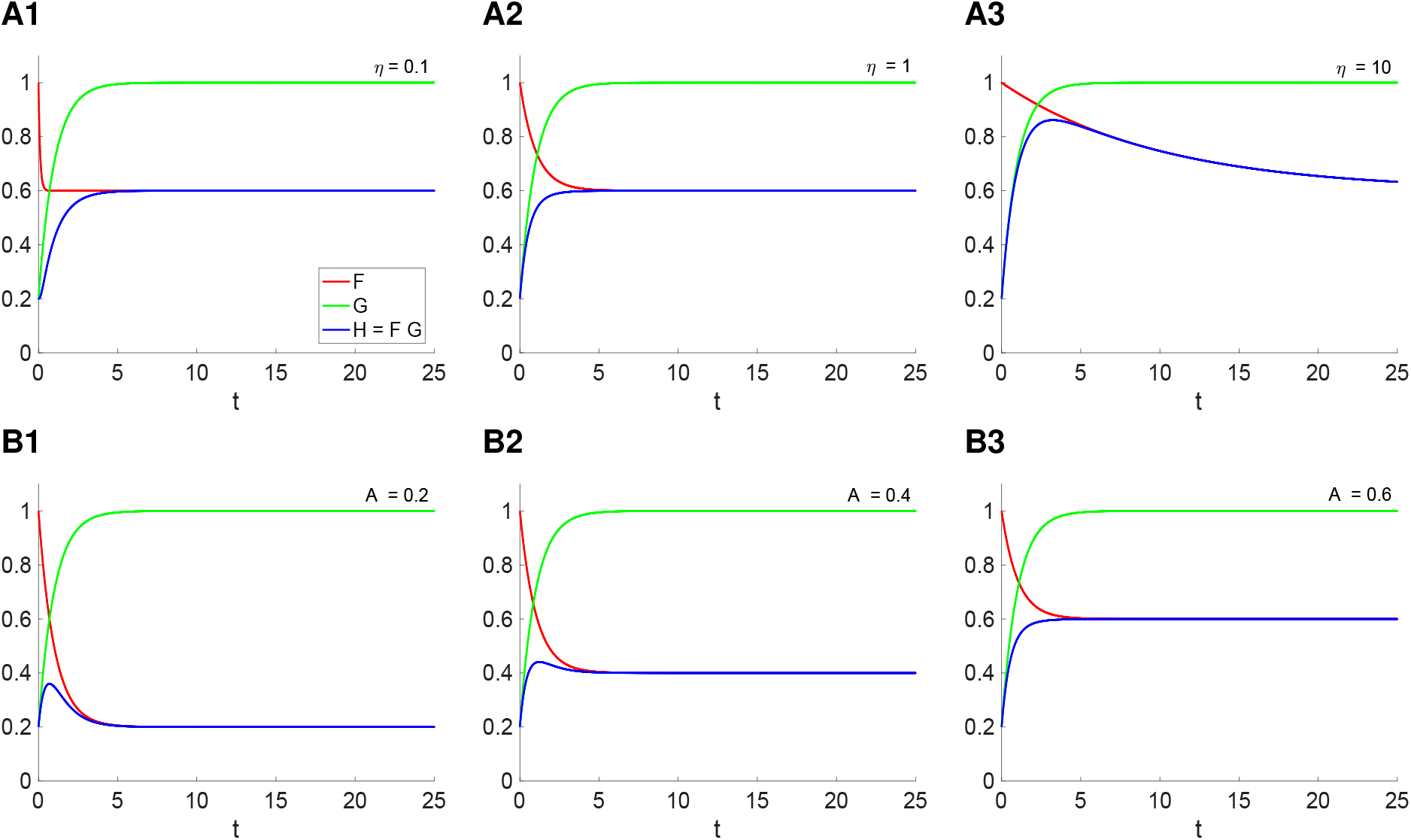
Temporal band-pass filters generated as the result of the multiplicative interaction of temporal low- and high-pass filters: envelope functions approach. We used the envelope functions *F* and *G* defined by (107) and (108), respectively, and *H* = *FG*. **A.** Increasing *η* contributes to the generation of a band-pass temporal filter. We used *A* = 0.5, *C* = 0.8 and **A1.***η* = 0.1. **A2.***η* = 1. **A3.***η* = 10. **B.** Decreasing *A* contributes to the generation of a band-pass temporal filter. We used *η* = 1, *C* = 0.8 and **B1.***A* = 0.2. **B2.***A* = 0.4. **B3.***A* = 0.6.

Changes in the parameter *B* in (108) affect the height of the band-pass temporal filter, but not the generation mechanism described above. However, for certain ranges of parameter values *H* is a temporal low-pass filter (not shown).

## Supplementary Material

### Extended analysis

**Figure S1:**
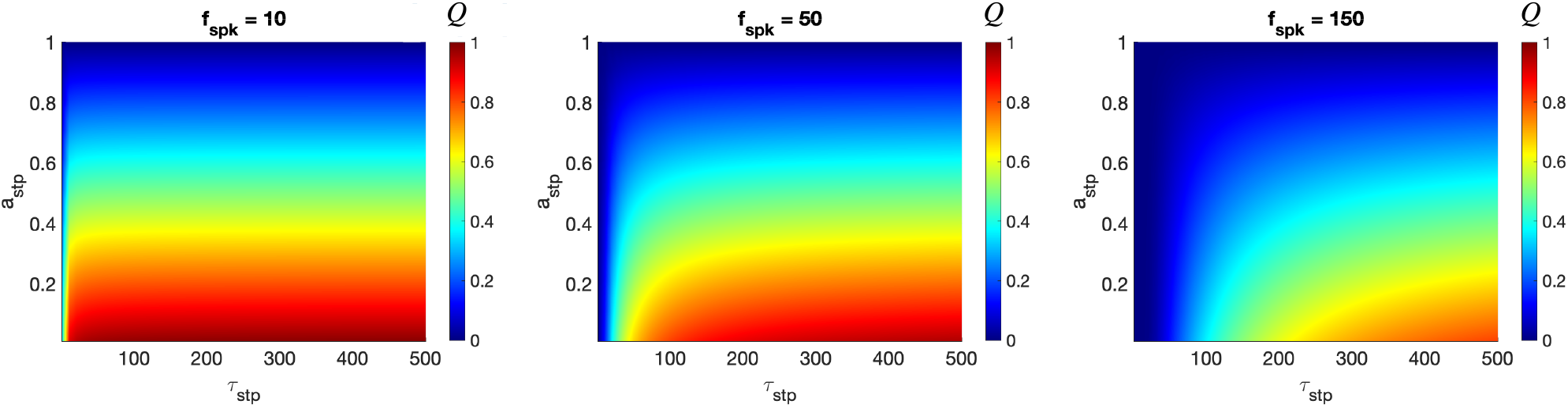
**Colormap of***Q*(*a_stp_, τ_stp_*) The colormaps show how *Q* (see eq 26) spanned over different values of *a_stp_* and *τ_stp_* behave. We consider *a_stp_* in the range [0:1] and *τ_stp_* in the range [0:500]. Every panel is computed for a different value of *f_spk_*. Notice that *Q*(*a_stp_, τ_stp_*) has, in general, lower values for higher *f_spk_*.

**Figure S2:**
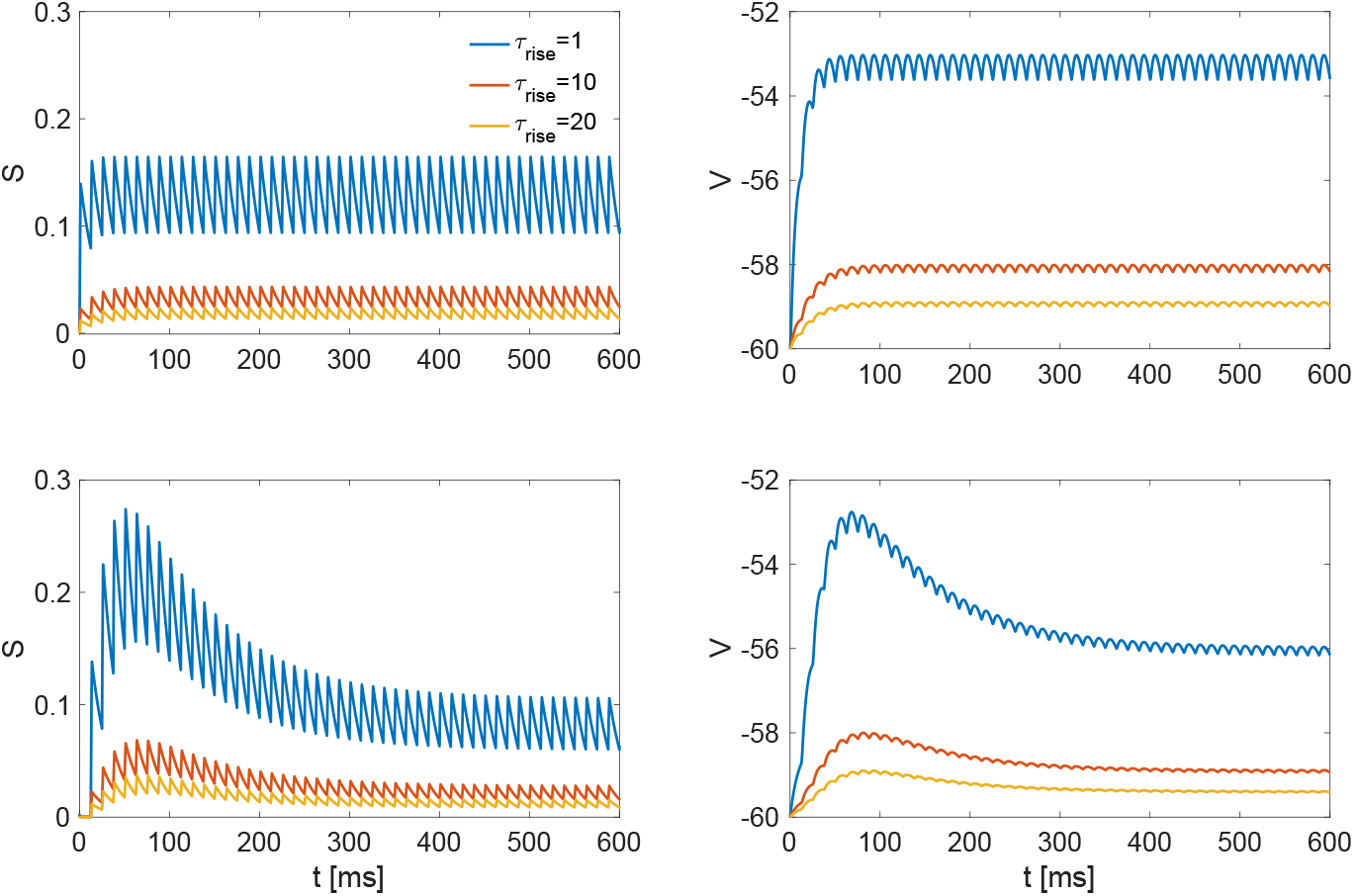
DA model combined with synaptic rise times. In every row we show the variable *S* and the variable *V* for a given rise time (see legend) without STP (first row) and with STP from the DA model (second row). Notice that for very small *τ_rise_* the increments in *S* and *V* are fast. When combined with STP, a band-pass filter shows up in *S* (see eqs 4–6). In this paper, we considered that rise times are very fast such as the ones found in AMPA and GABA *A*. However, in cases where *τ_rise_* increases our observations indicate that the band-pass filter is suppressed until it can no longer be observed. This effect happens because the presynaptic spikes take longer and longer to increase the value of *S* as *τ_rise_* increases until they become unnoticeable. We used the following parameters: *τ_dec_* = 20, *τ_dep_* = 400, *τ_fac_* = 50, *a_dep_* = 0.1, *a_fac_* = 0.2, and *f_spk_* = 80. The model for the rise times is taken from eq. (91) and assumes that a presynaptic spike has a time window of 1 ms which will be the time the postsynaptic membrane voltage takes to rise. We assume *k* = 1 mM.

**Figure S3:**
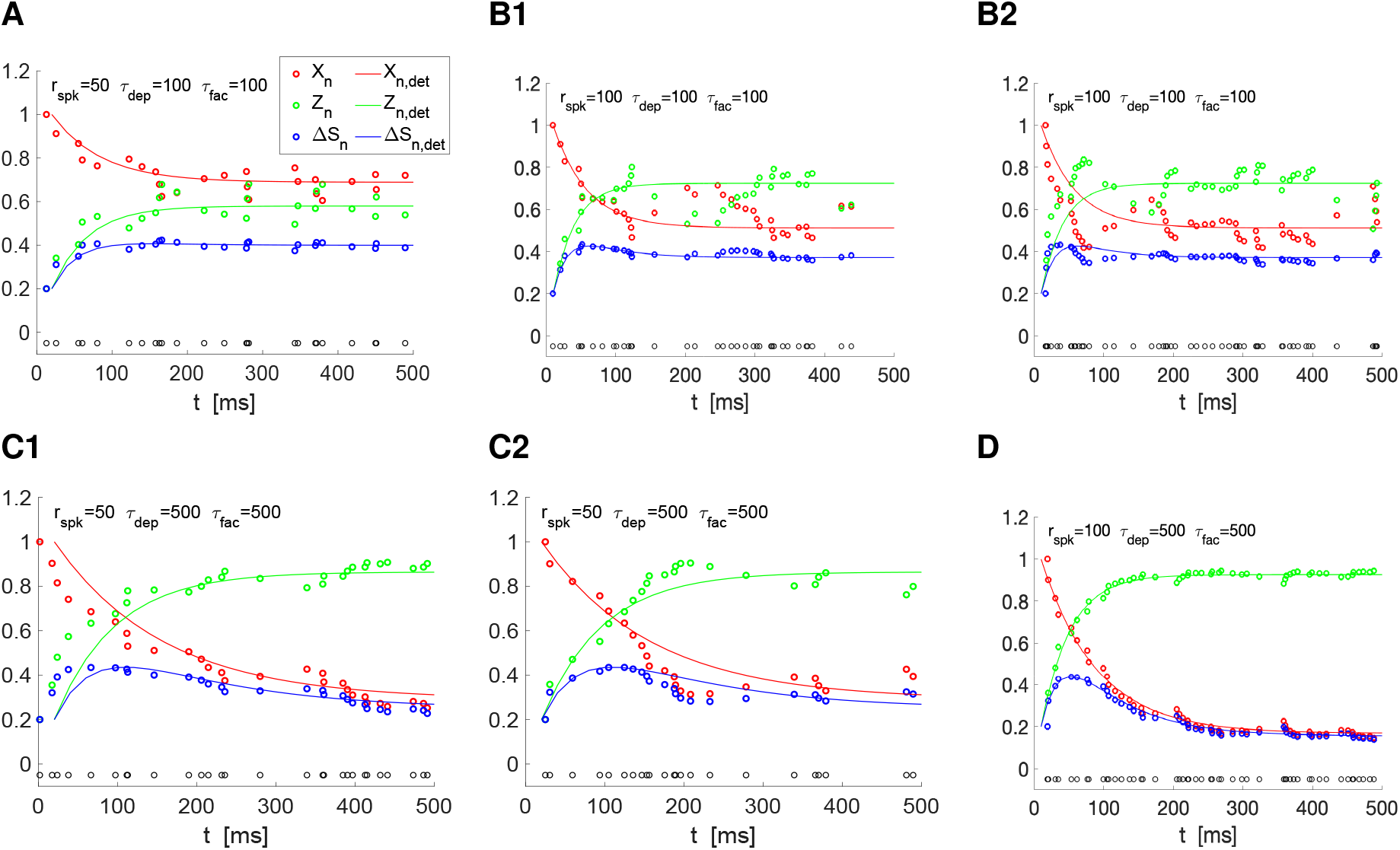
Temporal filters persist in response to variable presynaptic spike trains: Synaptic update response to Poisson distributed spike train inputs. We used the recurrent equations (7) and (8) for *X_n_* and *Z_n_* respectively. The ISIs have mean and standard deviation equal to *r_spk_*. Simulations were run for a total time *T_max_* = 200000 (Δ*t* = 0.01). **A, B.***τ_dep_* = *τ_fac_* = 100. **A.***f_spk_* = 50. **B.***f_spk_* = 100. **C, D.***τ_dep_* = *τ_fac_* = 500. **C.***f_spk_* = 50. **D.***f_spk_* = 100. We used the following parameter values: *a_d_* = 0.1, *a_f_* = 0.2, *x*_∞_ = 1, *z*_∞_ = 0.

### The Markran-Tsodyks (MT) model

First, we briefly remark on the notation used in this section. We conduct analysis using continuous extensions of Δ*S_n_*, *S_n_*, and *V_n_* – denoted as Δ*S*, *S*, *V* in the forthcoming figures and text. Time scales of temporal HPFs and LPFs in *S* are denoted *σ_f,S_* and *σ_d,S_*. The third time scale in temporal BPFs in *S* are denoted *σ_d_*_+*f,S*_. All temporal filters are fitted using gradient descent of a quadratic cost function.

The analysis for the MT model proceeds similarly to the DA models’. As seen in Section 3.3, the interaction between *X* and *Z* produces low-, high-, and band-pass temporal filters in Δ*S*. Similarly, *R* and *u* produce low-, high-, and band-pass temporal filters in Δ*S*. Again, we find Δ*S* temporal LPFs and HPFs not only develop in synapses exhibiting exclusively STD and STF, respectively. Indeed, Δ*S* temporal LPFs (HPFs) can develop in synapses where the time scale of depression (facilitation) dominates facilitation (depression). However, as in the case of the DA model, the exact ranges of *f_spk_* over which LPFs and HPFs develop depend on the balance between facilitation and depression. Figure S4-A1 shows that almost exclusively depressive synapses exhibit LPFs for 0 < *f_spk_* < 150, whereas a dominantly depressive synapse may stop producing LPFs for *f_spk_* > 100 (compare to Figure S4-A5). A similar situation arises in facilitating synapses (Figure S4-A2 and -A5).

*R_n_* and *u_n_* are well described by exponential decays, much in the same way *X_n_* and *Z_n_* are observed to be (Section 3.4). As such, as we did in the DA model, in the MT model we imagine that temporal filters in Δ*S* are heuristically the product of two exponentials. Despite the non-linearity present in MT model which complicates the analysis of how the long-term time scales of *R_n_* and *u_n_* are passed through their product Δ*S*, Δ*S* temporal LPFs (examples in Figures S5-B2,-B3) and HPFs (examples in Figures S5-A2,-A3) are still well described by a single time scale exponential. The time scales extracted from HPFs at dominantly facilitating and exclusively facilitating synapses are summarized in Figure S5-A1. Figure S5-B shows analogous results for LPFs of the MT model. A careful reader will note that Figure S5 refers to temporal filters of *S*, rather than Δ*S*. However, *τ_dec_* = 3 for these figures so that the contribution of the synaptic HPF implemented by synaptic decay is inconsequential for this discussion. The same remark also applies to Figures S6, S9, and S10.

In Section 3.6, BPFs in the DA model are shown to arise from 3 time scales, 2 of which can be extracted from corresponding LPFs and HPFs. A similar result is true for the MT model. Figure S6 outlines how these three time scales vary as the input frequency varies. Band-pass temporal filters become more sharply peaked as the input frequency increases. This reflects itself as observable decreases in the scenario where three time scales characterize band-pass temporal filters. Figure S6-B show that this third time scale is not superfluous – that removing the *σ_d_*_+*f,S*_ time scale from the model drastically alters the temporal filter fit.

Finally, we briefly review how Δ*S* temporal LPFs, HPFs, and BPFs are propagated to *S* and PSP. We note that *τ_dec_* and *τ_mem_* implement temporal HPFs. Figure S7 summarizes the types of filters that result in *S* from different incident Δ*S* temporal filters. Similarly, Figure S8 summarizes the types of filters that result in *V* from different incident *S* temporal filters. As in the DA model, we note there are instances where BPFs are passed through levels of organization and others where they arise due to an interaction of filters at different levels of organization (see Section 3.7). The communication through BPFs between levels of organization is exemplified by Figure S7-A3 (Δ*S* to *S*) and Figure S8-A3 (*S* to *V*). BPFs arising from interactions of filters between levels of organization are exemplified by Figure S7-A1 (Δ*S* to *S*) and Figure S8-A1 (*S* to *V*).

BPFs in the PSP arise in two ways – either passed through from the incident BPF filter or as the result of interacting an incident LPF and HPF implemented by the post-synaptic cell. These BPFs can be distinguished by analyzing the three time constants used to fit BPFs. First we consider synaptic BPFs that transfer to the post-syntactic cell’s response.

In this case, the procedure to extract the three time constants of the PSP BPF differs slightly from the procedure used to extract the three time constants of BPFs in *S* and Δ*S*. Instead of fixing LPF and HPF time scales and finding the best fitting third time constant, as was done in the Δ*S* BPFs, all three time constants for the PSP BPF are allowed to vary. In this way, we find a triple of time constants that fit the

**Figure S4:**
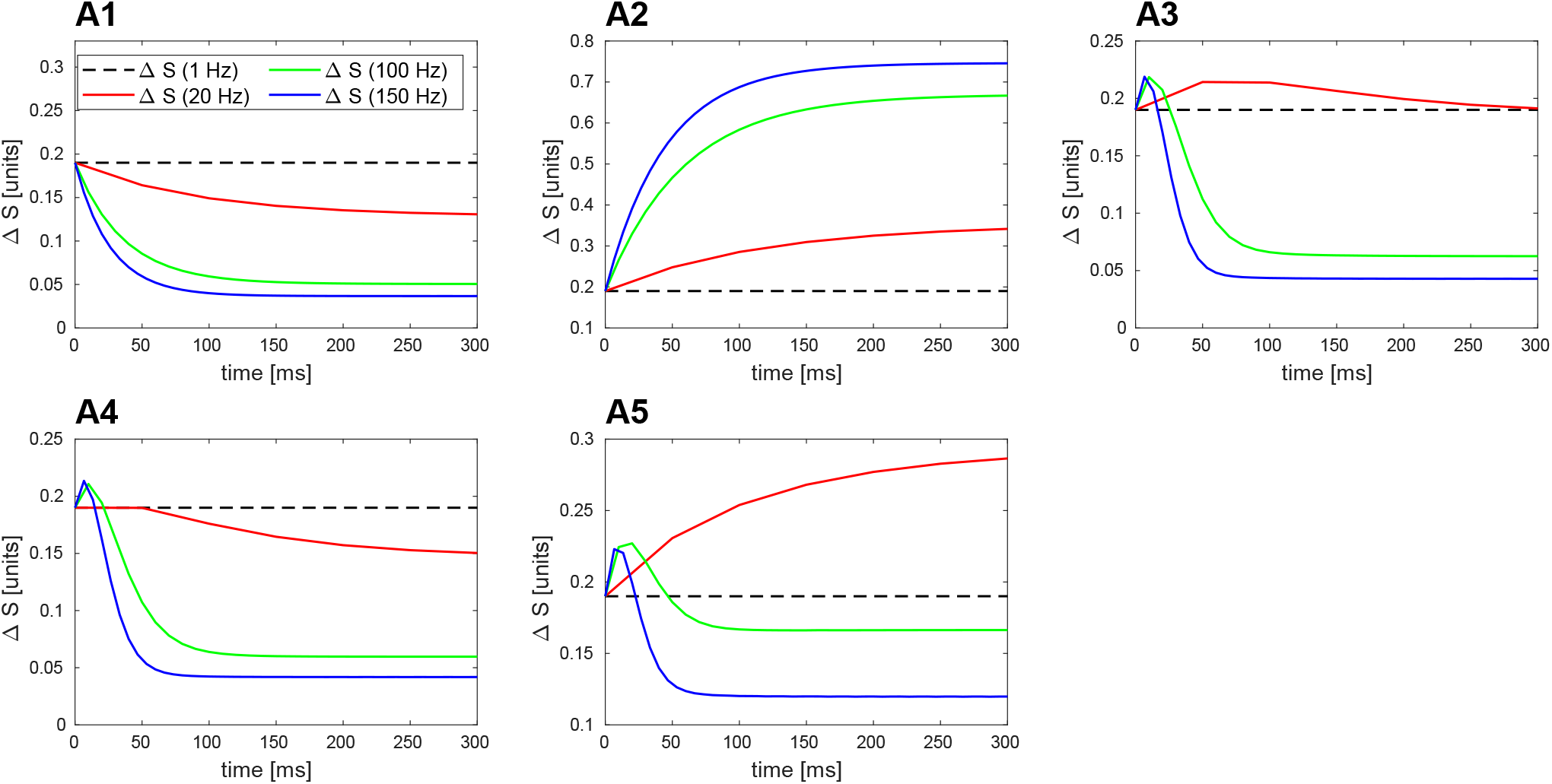
Temporal Filters in Δ*S* for the MT model. The interaction of presynaptic spiking and STP timescales create only high-pass, low-pass, and band-pass temporal filters. For these simulations *τ_dec_* = .01 to suppress summation. **A1.** Low-pass temporal filters appear for all input frequencies. As the input frequency increases, the low-pass temporal filters decay more aggressively (*τ_dep_* = 150, *τ_fac_* = 1). **A2.** High-pass temporal filters appear for all input frequencies. As the input frequency increases, the high-pass temporal filters rise more aggressively (*τ_dep_* = 1, *τ_fac_* = 150). **A3.** Band-pass temporal filters appear for all input frequencies. As the input frequency increases, the band-pass temporal filters become more sharply peaked (*τ_dep_* = 150, *τ_fac_* = 150). **A4.** Low-pass temporal filters appear for low input frequencies but then band-pass temporal filters develop for higher input frequencies (*τ_dep_* = 150, *τ_fac_* = 30). **A5.** High-pass temporal filters appear for low input frequencies but then band-pass temporal filters develop for higher input frequencies (*τ_dep_* = 30, *τ_fac_* = 150). In all simulations for the synapse: *U*_0_ = .1.

**Figure S5:**
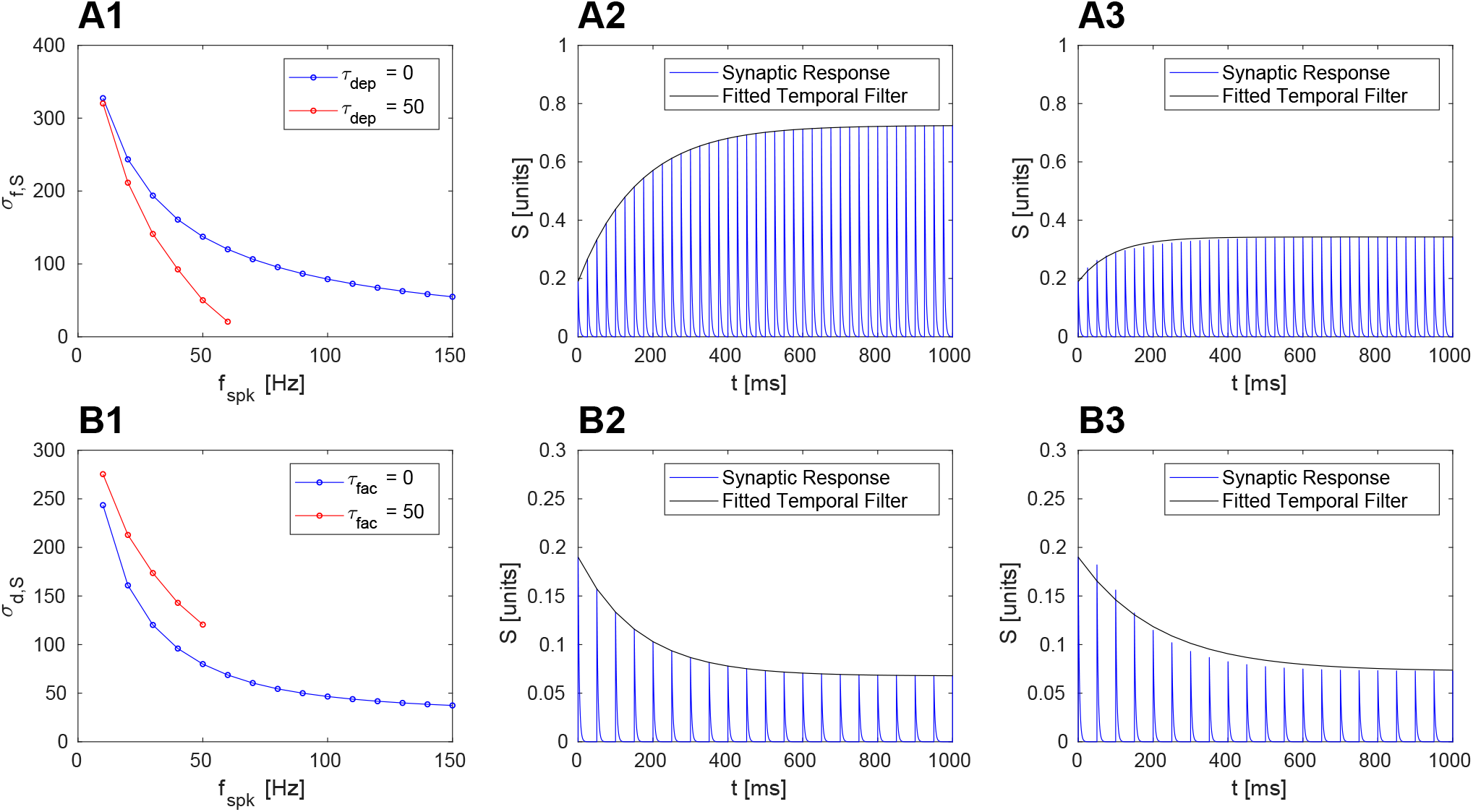
**A.** STF dominated synapses exhibit high-pass temporal filter. (*τ_fac_* = 500) **A1.** The dependence of high-pass temporal filter’s time scale on input frequency: a comparison of a synapse with no STD and fast STD. **A2.** Example of high-pass temporal filter at synapse with no STD. (*τ_dep_* = 0, *f_spk_* = 40) **A3.** Example of high-pass temporal filter at synapse with fast STD. (*τ_dep_* = 50, *f_spk_* = 40) **B.** STD dominated synapses exhibit low-pass temporal filter: a comparison of a synapse with no STF and fast STF. (*τ_dep_* = 500) **B1.** The dependence of low-pass temporal filter’s time scale on input frequency. **B2.** Example of low-pass temporal filter at synapse with no STF. (*τ_fac_* = 0, *f_spk_* = 20) **B3.** Example of low-pass temporal filter at synapse with fast STF. (*τ_fac_* = 50, *f_spk_* = 20) In all simulations for the synapse: *U*_0_ = .1 and *τ_dec_* = 3. Upper bound of RMSE on all low- and high-pass temporal fits: .012.

**Figure S6:**
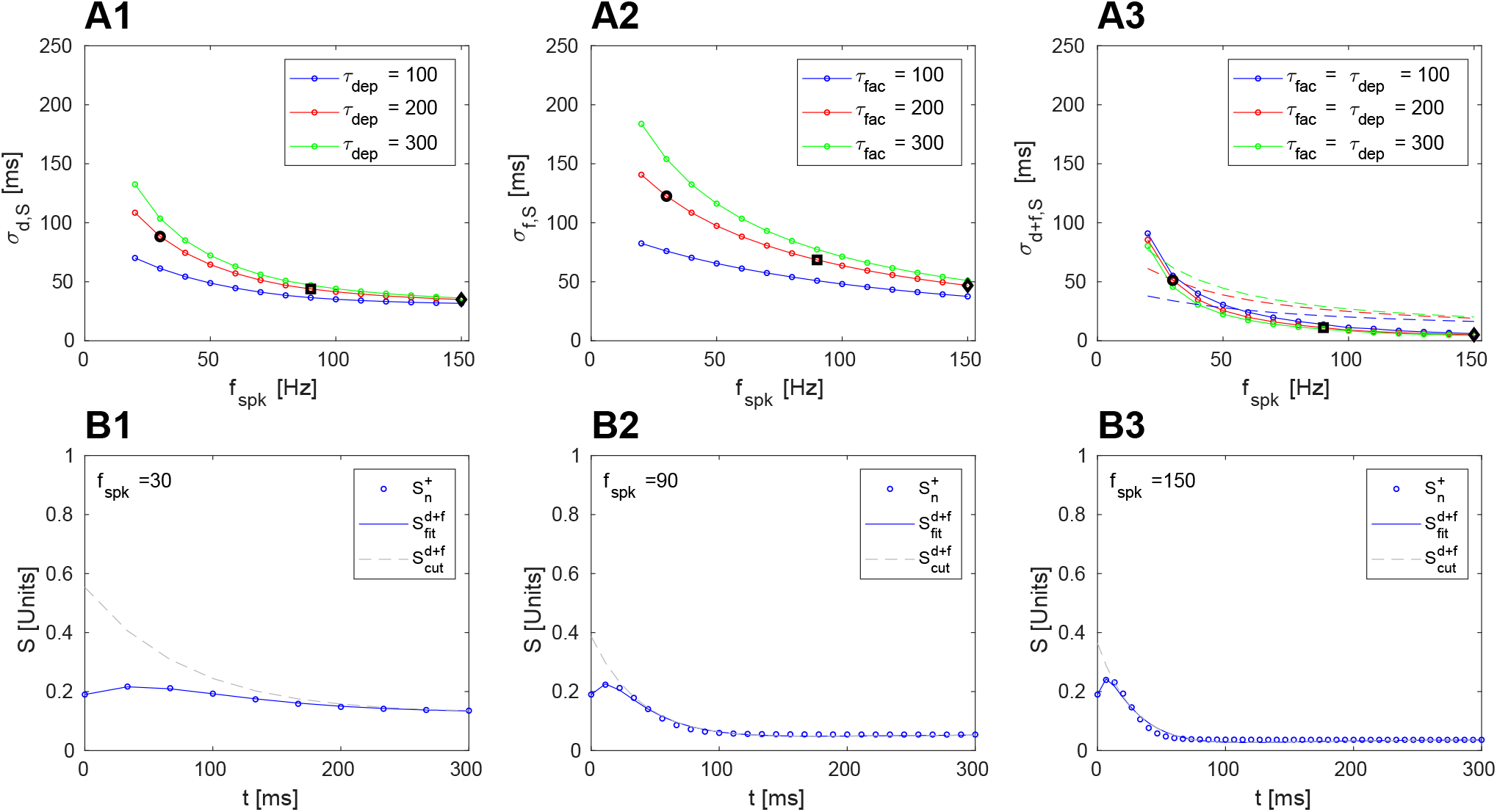
The time scales of band-pass temporal filters are determined by the mix of STF’s and STD’s time scales at the synapse along with input frequency. **A.** The time scales of STD and STF impact the three time scales characterizing band-pass filters in different ways. **A1.** The impact of STD’s time scale, *τ_dep_*, on the band-pass temporal filter’s time scale related to low-pass temporal filters. (*τ_fac_* = 0) **A2.** The impact of STF’s time scale, *τ_fac_*, on the band-pass temporal filter’s time scale related to high-pass temporal filters. (*τ_dep_* = 0) **A3.** The impact that both STD’s and STF’s time scale have on third time scale of temporal filtering. Solid lines are fit *σ_d_*_+*f,S*_ fitted from three time scale model of temporal BPFs. The dashed lines are the corresponding values of (1/*σ_d_* + 1/*σ_f_*)^−1^. **B.** The third time scale is not redundant. Without it, the temporal band-pass filter will fail to fit. (*τ_dep_* = *τ_fac_* = 200). The three time scales for the temporal filters in the following figures can be obtained from the foregoing three figures. 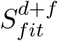 is the temporal BPF extracted using the three time scale model. 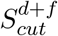 is obtained by setting the coefficient on the third time scale to zero. **B1.** Temporal BPF fit with and without third time scale when *f_spk_* = 30. The circles in A represent the time scales extracted using the three time scale model BPF model. **B2.** Temporal BPF fit with and without third time scale when *f_spk_* = 90. The squares in A represent the time scales extracted using the three time scale model BPF model. **B3.** Temporal BPF fit with and without third time scale when *f_spk_* = 150. The diamonds in A represent the time scales extracted using the three time scale model BPF model In all simulations for the synapse: *U*_0_ = .1 and *τ_dec_* = 3. Upper bound of RMSE on all temporal filter fits used in this figure: .01.

**Figure S7:**
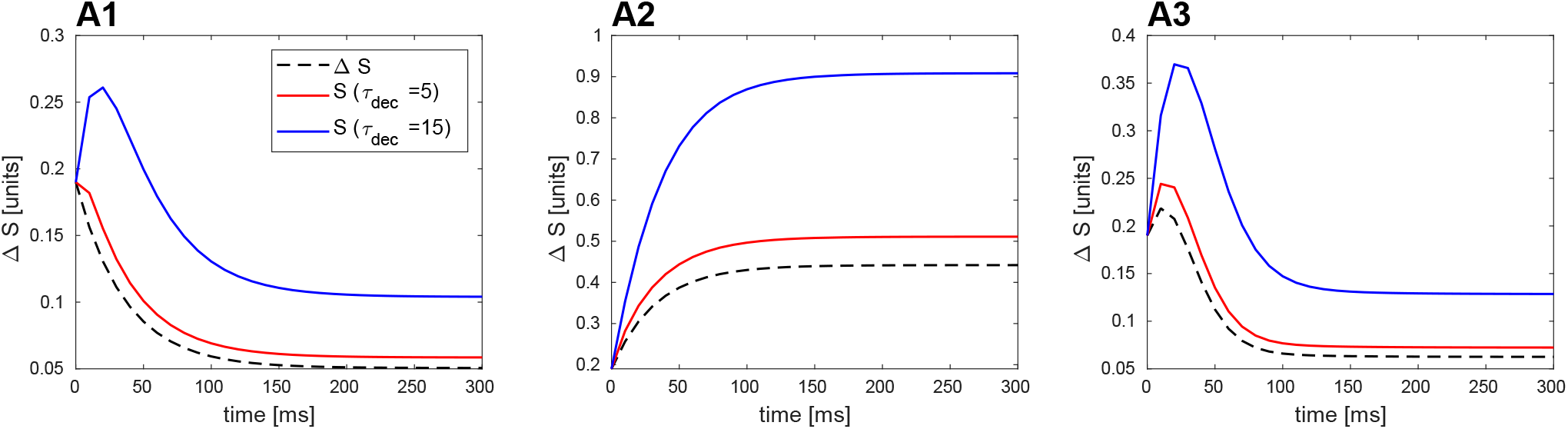
Temporal Filters in *S* for the MT model. Δ*S* create only low-pass, high-pass, and band-pass temporal filters. Now we examine the effect that summation has on these temporal filters. The effect of summation can be understood to be a high-pass temporal filter interacting with a temporal filter created from the interaction of input and STP time scales (shown here for *f_spk_* = 100). **A1.** The interaction of a low-pass temporal filter in Δ*S* with the time scale of summation, *τ_dec_* is shown. We observe that low-pass temporal filters become band-pass temporal filters for longer time scales of summation (*τ_dep_* = 150, *τ_fac_* = 1). Note that for extreme, unphysiological time scales of summation (*τ_dec_* > 150), high pass temporal filters in *S* may also develop (not shown). **A2.** The interaction of a high-pass temporal filter in Δ*S* with the time scale of summation, *τ_dec_* is shown. We observe that high-pass temporal filters remain high-pass temporal filters for any time scale of summation (*τ_dep_* = 1, *τ_fac_* = 150). **A3.** The interaction of a band-pass temporal filter in Δ*S* with the time scale of summation, *τ_dec_* is shown. We observe that band-pass temporal filters remain band-pass temporal filters (*τ_dep_* = 150, *τ_fac_* = 150). Longer time scales of summation increase the size of the band-pass peak (both in height and width). Note that for extreme, unphysiological time scales of synaptic decay (*τ_dec_* > 150), high pass temporal filters in *S* may also develop (not shown). In all simulations for the synapse: *U*_0_ = .1.

**Figure S8:**
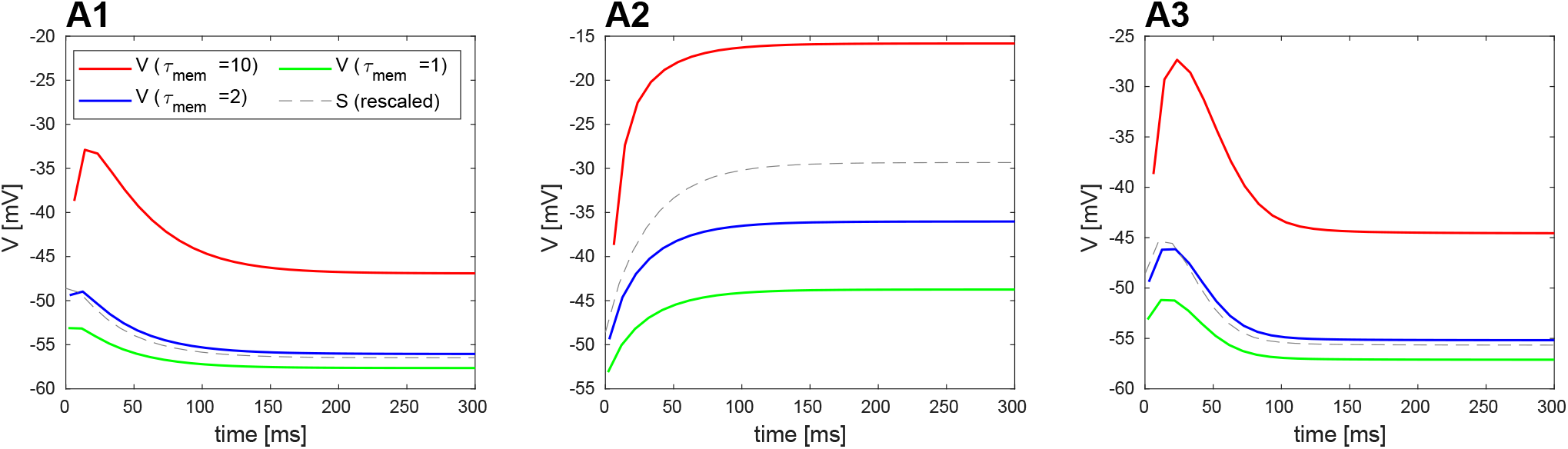
Temporal Filters in *V* for the MT model. The interaction of input, STP, and summation time scales combine to create only low-pass, high-pass, and band-pass temporal filters. Now we examine the effect that the membrane time constant of the post-synaptic cell has on the temporal filters incident from the synapse. For fast enough membrane time constants (*τ_mem_* < 1 ms, *g_L_* > 1) the post-synaptic temporal filter reflects the synaptic temporal filter (*f_spk_* = 100, *τ_dec_* = 5). As the membrane time constant slows, the synaptic temporal filter interacts with the post-synaptic cell. Biophysically, the effect of a slow membrane time constant is to create post-synaptic summation – resulting in a high-pass post-synaptic temporal filter. The development of an interaction between the high-pass post-synaptic temporal filter with the synaptic temporal filter is seen in these figures as the membrane time constant slows. For these figure *G_L_* = 0.1, 0.5, and 1. These correspond to *τ_mem_* = 10 ms, 2 ms, and 1 ms, respectively. **A1.** The interaction of a low-pass synaptic temporal filter in *S* (rescaled and shown in gray) with the membrane time constant is shown. We observe that low-pass synaptic temporal filters can become band-pass temporal filters as the membrane time constant slows (*τ_dep_* = 150, *τ_fac_* = 1). **A2.** The interaction of a high-pass synaptic temporal filter in *S* (rescaled and shown in gray) with the membrane time constant is shown. We observe that high-pass synaptic temporal filters remain high-pass temporal filters in the post-synaptic response as the membrane time constant slows (*τ_dep_* = 1, *τ_fac_* = 150). **A3.** The interaction of a band-pass synaptic temporal filter in *S* (rescaled and shown in gray) with the membrane time constant is shown. We observe that band-pass synaptic temporal filters remain band-pass temporal filters in the post-synaptic cell (*τ_dep_* = 150, *τ_fac_* = 150). In particular, as the membrane time constant slows, the size of the band-pass peak increases (both in height and width). We remark that for all cases, for extremely slow membrane time constants (*τ_mem_* > 1 sec or *g_L_ < .*001), high-pass post-synaptic temporal filters develop (not shown). In all simulations for the synapse: *U*_0_ = .1, *τ_dec_* = 5. In all simulations of the post-synaptic cell: *G_ex_* = 1, *C* = 1, *E_L_* = −60, *E_ex_* = 0.

PSP BPF: *ρ_a_, ρ_b_* and *ρ_c_* (Example shown in Figure S9-A3). Then we introduce the following quantities:

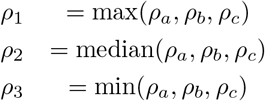

These time constants are then compared to the magnitude of the time constants obtained from the synaptic BPF. In Figure S9-C we note that the synaptic BPF has time constants such that *σ_d,S_ > σ_f,S_ > σ_d_*_+*f,S*_. Assuming the post-synaptic cell maintains this relation, we associate *ρ*_1_ with *σ_d,S_*, *ρ*_2_ with *σ_f,S_*, and *ρ*_3_ with *σ_d_*_+*f,S*_. Figure S9-C suggests that the HPF implemented by the membrane time constant is modifying all three time constants of the incident synaptic BPF.

BPFs in the post-synaptic cell are implemented also by the interaction with synaptic LPFs and the HPF implemented by the membrane time constant. One may use the same three parameter model to fit these PSP BPFs. However, we find that there are actually two time scales describing the shape of these BPFs. Figure S10-B show explicit examples of the third time scale’s insignificance. Furthermore, one of the time scales describing the PSP BPF is shown to be inherited from the incident LPF’s time scale of decay.

Figure -B1 and -B2 show how the membrane time constant also modifies incident LPF and HPF time constants in the post-synaptic cell. Here, the same models to extract time constants of rise for HPFs and time constants of decay for LPFs in Δ*S* also work well for extracting time constants for LPFs and HPFs of the PSP (example fits in Figure S9-A1 and -A2). In this figure, *σ_f,V_* is the time constant of rising in PSP HPFs and *σ_d,V_* is the time constant of decay in PSP LPFs, respectively.

In Section 3.9.2, we discuss how the temporal filters implemented by *Z_n_*, *X_n_*, and Δ*S_n_* persist in the presence of Poisson spiking. Here we review a potential consequence of Poisson inputs interacting with temporal filters: gain control. The data in the following figure, Figure S11, uses the MT model, however, the foregoing discussion suggests that the DA and MT model both exhibit temporal filters, albeit via slightly differing quantitative mechanisms. Figure S11 plots average amplitude of voltage response over 1100 trials where the input Poisson rates change over time. As the spiking rates change over time, different steady states are achieved. Overshooting transient features develop between rate changes when STD time scales increase. The fact that these overshoots depend on the time scale of STD suggests that there may be a connection between the overshoot magnitudes and the LPFs that STD implement. Furthermore, the magnitude of these average rate changes and their dependence on the initial and final rates was studied as a mechanism for gain control by Abbott *et. al.*. Figure S11 show that average amplitude also depends on the time scale of STD and membrane time scales - ergo, it follows that filters that STD and membrane time constants implement may also play an important role in determining the average amplitude of voltage response.

**Figure S9:**
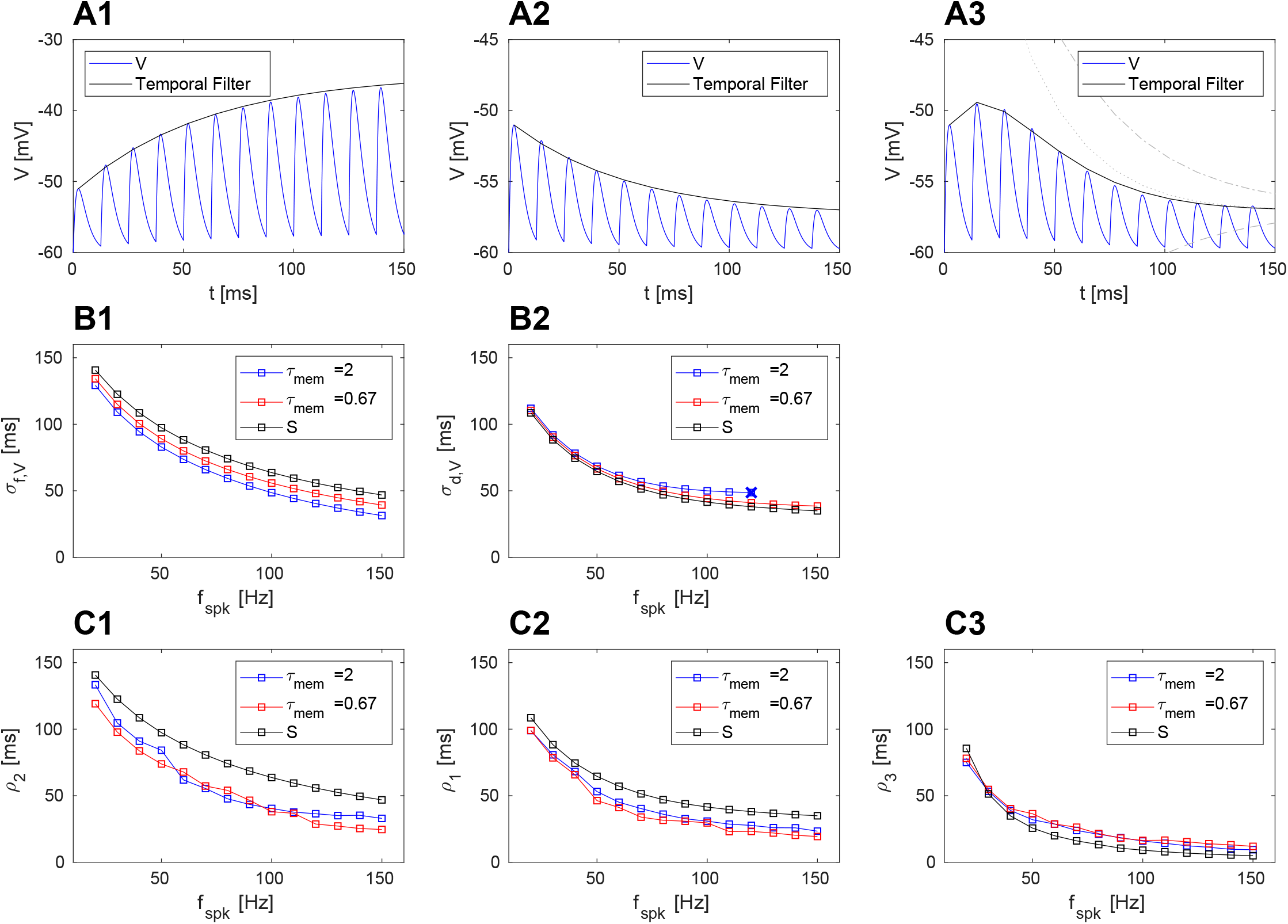
**A.** Representative fits of low-, high-, and band-pass temporal filters in passive post-synaptic cell. (*G_L_* = .5) **A1.** Example of high-pass post-synaptic temporal filter. (*τ_dep_* = 0, *τ_fac_* = 200, *f_spk_* = 80) **A2.** Example of low-pass post-synaptic temporal filter. (*τ_dep_* = 200, *τ_fac_* = 0, *f_spk_* = 80) **A3.** Example of band-pass post-synaptic temporal filter. (*τ_dep_* = 200, *τ_fac_* = 200, *f_spk_* = 80) **B1.** The impact that membrane time constant has on the time scale of post-synaptic high-pass temporal filters. The incident synaptic temporal filter is high-pass and given by *τ_dep_* = 0, *τ_fac_* = 200. **B2.** The impact that membrane time constant has on the time scale of post-synaptic low-pass temporal filters. The incident synaptic temporal filter is low-pass and given by *τ_dep_* = 200, *τ_fac_* = 0. “X” marks the input frequency at which the post-synaptic temporal filter transitions from low-pass to band-pass. These band-band pass temporal filters are analyzed in Figure S10. **C.** The impact that membrane time constant has on the time scales of post-synaptic band-pass temporal filters. The incident synaptic temporal filter is band-pass and given by *τ_dep_* = 200, *τ_fac_* = 200. The way the time scales, *ρ*_1_, *ρ*_2_ and *ρ*_3_, are obtained are outlined in Methods. **C1.** The impact of the membrane time constant on *ρ*_1_. **C2.** The impact of the membrane time constant on *ρ*_2_. **C3.** The impact of the membrane time constant on *ρ*_3_. All simulations were performed using MT Model with *U*_0_ = .1 and *τ_dec_* = 3. The parameters for the passive cell are *G_ex_* = 1, *C* = 1, *E_L_* = −60, *E_ex_* = 0. The upper bound on the RMSE for all temporal filters (low-, high-, and band-pass) is .4 mV. The upper bound on the maximum difference between a fit and the temporal filters for all voltage responses is 1 mV.

**Figure S10:**
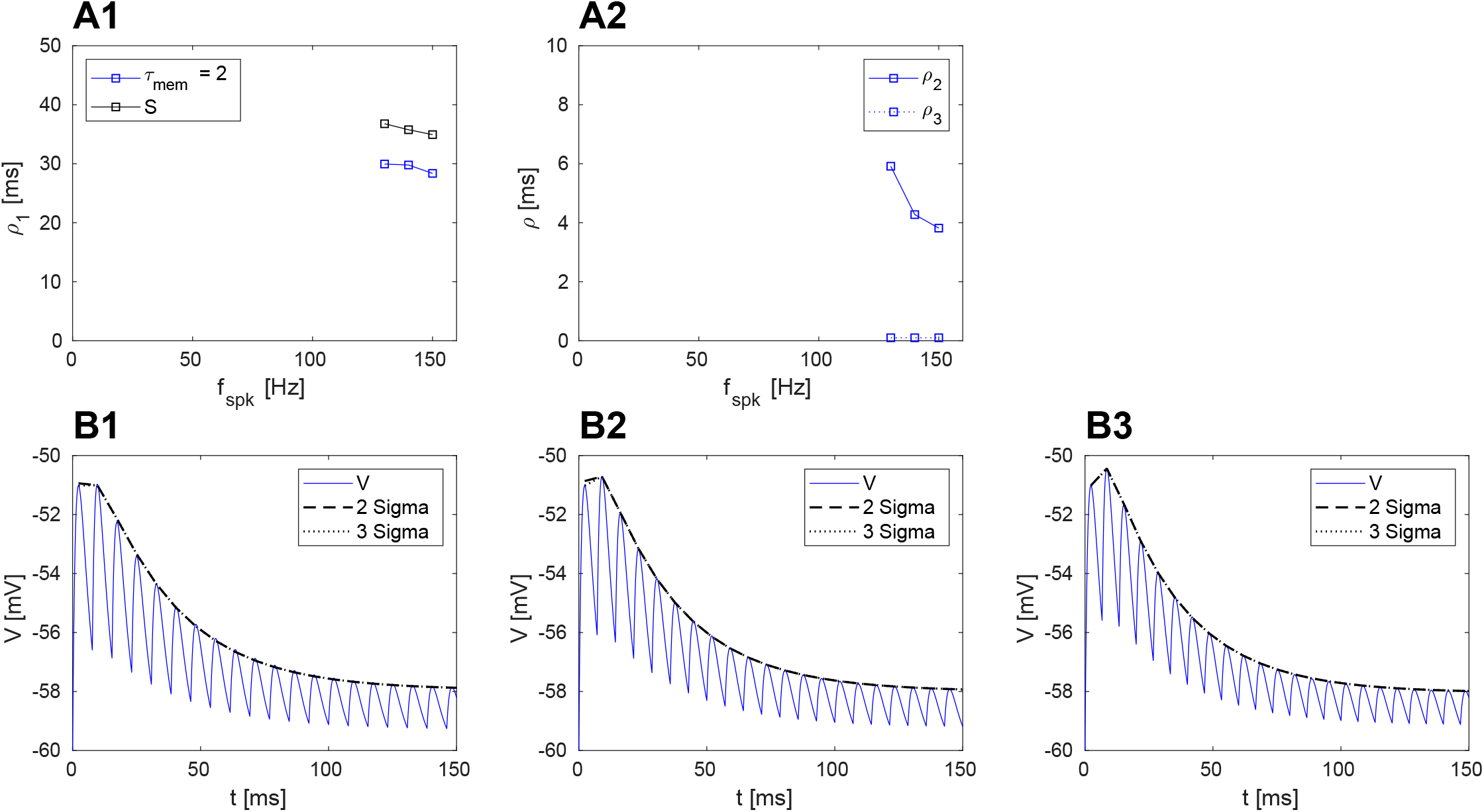
**A.** Time scales of band-pass temporal filters formed by incident low-pass synaptic temporal filters. The incident synaptic temporal filter is low-pass and given by *τ_dep_* = 200, *τ_fac_* = 0. **A1.***ρ*_1_ plotted as a function of frequency and compared to *σ_d_*. **A2.***ρ*_2_ and *ρ*_3_ plotted as a function of frequency. **B.** The three temporal band-pass temporal filters plotted in A. “3 sigma” is the fit obtained using BPF filter model letting all three time scales vary. “2 sigma” is the fit obtained omitting the fastest time scale, *ρ*_3_, from the fit. (*G_L_* = .4) **B1.** Temporal band-pass temporal filters plotted in A with its fits. (*f_spk_* = 130) **B2.** Temporal band-pass temporal filters plotted in A with its fits. (*f_spk_* = 140) **B3.** Temporal band-pass temporal filters plotted in A with its fits. (*f_spk_* = 150) All simulations were performed using MT Model with *U*_0_ = .1 and *τ_dec_* = 3. The parameters for the passive cell are *G_ex_* = 1, *C* = 1, *E_L_* = −60, *E_ex_* = 0. The upper bound on the RMSE for all temporal filters (low-, high-, and band-pass) is .4 mV. The upper bound on the maximum difference between a fit and the peaks of voltage response is 1 mV.

**Figure S11:**
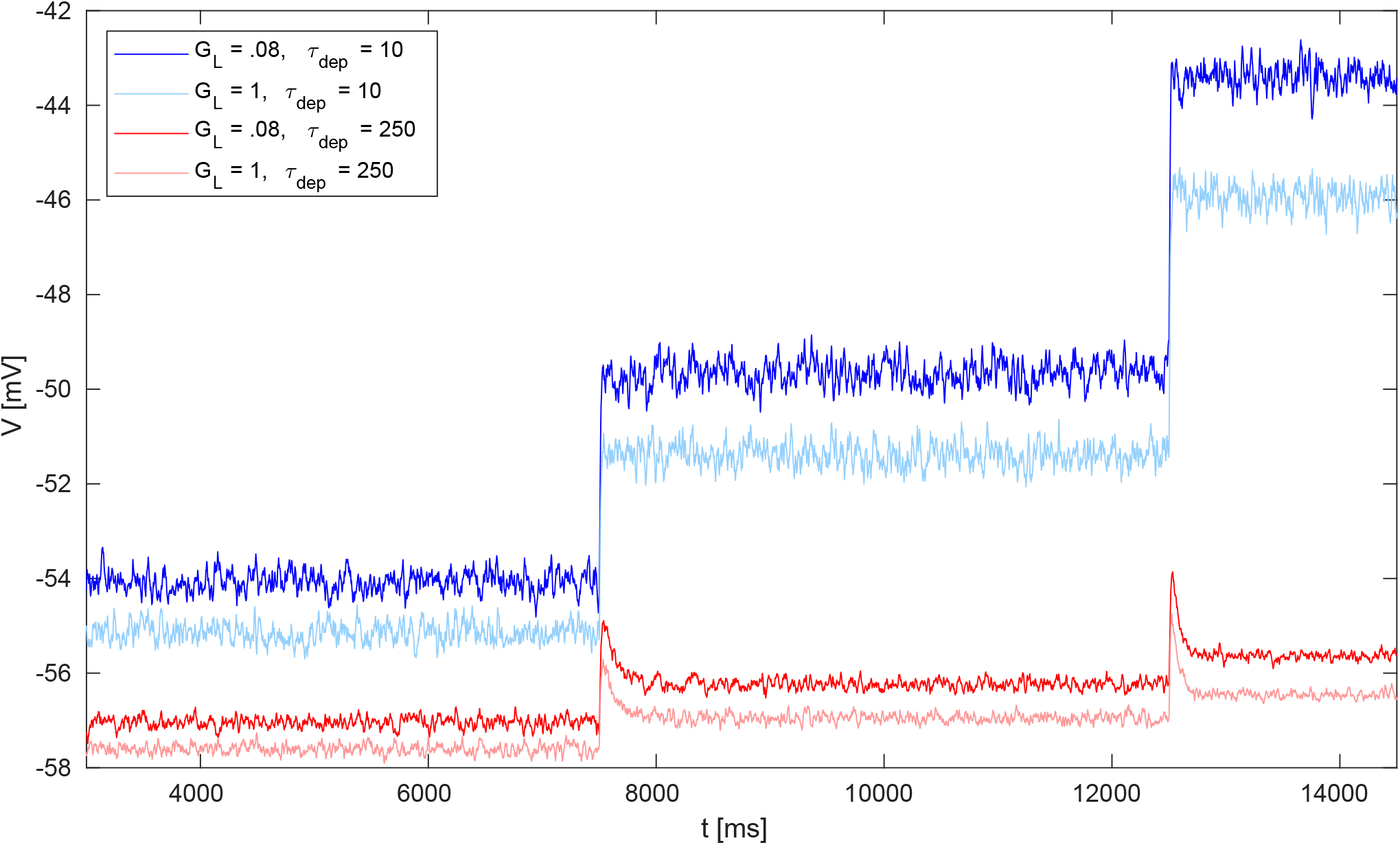
Each trace in the figure is the average of 1100 trails. A trial consists of the following: 7.5 secs of 20 Hz Poisson stimulus, 5 secs of 40 Hz Poisson stimulus, and 2 secs of 80 Hz Poisson stimulus. The first second of the simulation is cut off to remove the transient behaviors from the initialization of the simulation. All simulations were performed using MT model with parameters: *τ_fac_* = 0, *τ_dec_* = 3, *U*_0_ = .1. Passive post-synaptic cell parameters were *C* = 1, *G_ex_* = .1, *E_L_* = −60, *E_ex_* = 0.

